# Incomplete references leave bulk deconvolution targets non-identifiable, but identification has a computable precision price

**DOI:** 10.64898/2026.09.10.750775

**Authors:** Hanyu Jiang, Fei Gao, Peixi Liu, Yaheng Wu, Ying Jie, Ying Li, Yang Jiang

**Affiliations:** Department of Ophthalmology, Peking Union Medical College Hospital, Chinese Academy of Medical Sciences and Peking Union Medical College, Beijing 100730, China; Beijing Key Laboratory of Fundus Diseases Intelligent Diagnosis & Drug/Device Development and Translation, Beijing 100730, China; Key Laboratory of Ocular Fundus Diseases, Chinese Academy of Medical Sciences, Beijing 100730, China; Eight-Year Medical Doctor Program, Chinese Academy of Medical Sciences and Peking Union Medical College, Beijing 100730, China; CAMS Key Laboratory of Translational Research on Lung Cancer, State Key Laboratory of Molecular Oncology, Department of Medical Oncology, National Cancer Center/National Clinical Research Center for Cancer/Cancer Hospital, Chinese Academy of Medical Sciences and Peking Union Medical College, Beijing 100021, China; Beijing Institute of Ophthalmology, Beijing Tongren Eye Center, Beijing Tongren Hospital, Capital Medical University, Beijing 100730, China

**Keywords:** deconvolution, reference incompleteness, cell-type composition, measurement operator, identifiability, partial identification, provenance

## Abstract

**Background:** Reference-based deconvolution estimates cell-type proportions from bulk profiles. Incomplete references compromise these estimates, yet many remedies return point estimates. We separate the operator provenance that reproduces an estimate from the information and precision that identify its target.

**Results:** Two operator histories at one reduced reference give different estimators: 28.2% of 196,420 sample-deletion pairs differed by >0.1 total variation; locking every learned component made paths identical. Across nine cohorts, operator history reversed 160 of 952 association signs, 31 with a significant path, and changed significance for 126. Observationally equivalent completions can fill the open simplex and reverse retained-type rankings. Under a joint zero-exposure condition, shared structure leaves inherited bounds unchanged; one to six uncalibrated views also gave identical bounds. A profile library contracted estimator-output envelopes by 98.93% yet covered the full-reference effect for 42.77%, with 19 wrong-sign certificates; conditional sharp bounds stayed at [−1, 1]. Treating RNA yields as exact collapsed intervals to points covering none of seven flow-measured targets: contraction without coverage is false certainty. Calibrated cross-modal anchors contract width to 0.51 at six types but certify no valid sign. Exact RNA yields identify cell fractions from RNA contributions; a decisive sign in blood requires ±2.6% proxy accuracy with near-total contraction of donor-heterogeneity and dynamic-range envelopes.

**Conclusions:** Incomplete-reference deconvolution is an identification problem, not only an estimation problem. Remedies must be scored on shrinkage, coverage, certification and false certification against held-out targets. fitdrop implements this scoring and a precision frontier for planning decisive measurements.

## Background

Bulk-tissue deconvolution is a workhorse of genomics [1–3]: given a bulk profile and a reference of cell-type signatures, it estimates the proportions of the constituent types. Downstream analyses treat those proportions as measurements, comparing them across cohorts, regressing them on phenotypes and tracking them across studies. The references, however, are moving targets. Single-cell atlases differ in which populations they resolve, dissociation and preservation protocols deplete fragile types before a reference is built [4], and updated atlas versions refine, change and add types [5]. Whenever two analyses use references with different column sets, the question of whether their composition estimates are comparable is unavoidable.

In systematic benchmarking, an incomplete reference degraded results regardless of every other pipeline choice [6], and methodological reviews list it as an open challenge [2, 3, 7]. The field’s responses fall into two families. One is to ensure that the reference is complete, but this is not operational, because populations are missing from references precisely when they are rare, fragile or unresolved. The other comprises semi-reference and completion-recovery methods that introduce latent unknown profiles under explicit optimization or modeling assumptions [8–14]. Residual-based recovery can reconstruct missing-type profiles when the number of missing types is known [15]. Every member of this family returns a point estimate. None of them reports, and no study we could find asks, how much of that number is determined by the observed data and how much by the chosen assumptions.

Prior work has measured what an incomplete reference does to accuracy. Deletion experiments in bulk RNA benchmarking established that removing a cell type redistributes its signal non-uniformly across the remaining types [6, 16]. In DNA methylation, leave-one-type-out retraining inflates a dedicated error metric monotonically with the missing type’s abundance [17], and deep-learning frameworks extend incomplete-reference deconvolution across omics [18]. The sensitivity of composition estimates to the choice of reference is well documented [19, 20]. That unrepresented cell types leave the inverse problem under-determined has been acknowledged at most qualitatively. Recent identifiability theory for supplied-reference deconvolution and geometric analyses of non-negative factorization concern different ambiguity classes [21, 22].

An identified set is the set of target values that some admissible completion makes compatible with the observations. We found no formal identified-set analysis of the retained-composition target over admissible incomplete-reference completions in genomic deconvolution. Existing work on this axis stops at point estimates and does not connect completion ambiguity to operator histories crossed at a fixed terminal reference.

Prior omission designs, moreover, perform different reference edits. Some delete a signature profile from a fixed matrix and re-run the solver [16]. Others rebuild reduced references with method-dependent internal steps [6]. Some carry feature support derived from the full reference into depleted-reference runs while other branches reconstruct their references natively [15], and the original MuSiC paper itself contains an incomplete-reference robustness experiment [23]. No study we found crosses the two ends of this spectrum, a fully frozen operator state and a fully rebuilt one, while holding the observed terminal endpoint fixed. A benchmarking framework that decomposes pipelines into reference-construction and regression components [24] could express the contrast but does not report it. The deletion experiment and the operator refit are bundled, in varying and mostly unreported proportions, into a single arm.

This bundling hides a distinction that matters, because a reference is two things at once. It is the design matrix of the solve, but for a large class of estimators it is also the training data for the measurement operator. A marker library [25], per-gene calibration centers and scales [26], cross-subject variance weights [23] and solver hyperparameters [27] are learned quantities, some from the reference alone and some jointly with the mixtures during fitting. Deleting a column therefore changes the estimation problem twice: it removes one unknown, and it changes the operator that will measure all remaining unknowns. Beyond the operator, the reference is also a completion assumption, a claim that its columns suffice to explain the bulk, and its failure raises an identification question that no amount of algorithmic improvement can settle from the observed data alone.

Here we treat these as two separate failures of uniqueness and study each with the tool it requires. For the operator, we run every deletion twice on the identical reduced reference. A **frozen** path (F) carries operator state learned on the full reference, with deletion applied only at the solve step. A **refit** path (R), which we also call rebuilt, re-learns the operator state on the reduced reference and is the arm closest to what prior deletion studies ran. Both paths use the same bulk, terminal reduced reference, constraint set and solver, differing only in the history of the operator state.

For the target, we construct exact observationally equivalent completions and ask what they do to the quantity a practitioner would report. Reference incompleteness has been treated as an empirical bias [6, 15–17, 28]. We treat it as an identification problem with a computable price. Endpoint-matched histories show that identical bulk data and an identical reduced reference yield different compositions and different disease associations because the measurement operator retains its history, and orthogonal imaging shows that neither history is consistently more accurate. Exact completions then show that fixing the operator does not identify the retained-composition target. We score shared multi-sample structure, additional measurement views, external profile libraries, calibrated cross-modal anchors and absolute quantification by whether they narrow conclusions while retaining coverage. Inverting the measurement-uncertainty models then states the precision a declared conclusion requires and the uncertainty source that must be reduced first. The result separates reproducing an estimate, identifying its target and designing the measurement that resolves the remaining ambiguity.

## Results

### Fitting and deleting do not commute

The first failure of uniqueness is operator memory: the same terminal reduced reference can carry different learned measurement states. We formalize the paths through the operator state, using the term broadly. It includes both static components learned from the reference (feature support, transformations, weights, hyperparameters) and, for iterative estimators, bulk-conditioned solver state induced jointly by the bulk and the reference. Let *θ_A_*(*y*) denote this complete history-carrying state (for estimators with purely reference-learned state the dependence on y drops out) and g the solver. The frozen path carries *θ_A_*(*y*) through deletion, restricted to the retained system: *g*(*y, A_−d_; θ_A_*(*y*) restricted). The refit path recomputes 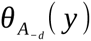. Both consume the same bulk profile *y* and the same terminal reduced reference *A_−d_*. Operator memory is present when the restriction of *θ_A_*(*y*) fails to equal 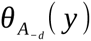. Operator provenance is the record of which history produced the state in use. The same contrast admits a second reading that matters for practice: the frozen path is exactly the history “a type was added to the reference and later removed” while the refit path is “the type was never there”. Every reported F/R disagreement therefore also measures the operator scar left by a round-trip reference edit.

The two paths disagree (Fig. 1), so deletion changes the estimator as well as the problem. BisqueRNA’s reference-based branch has the simplest learned stage: it learns per-gene affine calibration from reference-derived donor pseudo-bulk [26] and, as run here with `markers = NULL`, performs no marker selection. On this estimator, the operator-history separation, the distance between frozen and refit compositions, had median Aitchison values [29] from 0.70 to 1.37 across cornea, dorsolateral prefrontal cortex (DLPFC), lung and blood (labelled PBMC after its peripheral blood mononuclear cell reference panel) source panels (Additional file 1: Table S1; 73,588 paired sample solutions). Fewer than 1.6% of samples in any tissue fell below 10⁻⁸. On the probability scale, the frozen-versus-refit displacement exceeds 0.1 in total variation (TV) for 28.2% and 0.25 for 8.7% of sample-deletion pairs (196,420 pairs over the 11 of 17 tissue-estimator groups with stored per-sample vectors). Median TV was 0.0433, 0.0578 and 0.0409 for the calibration, support-vector regression (SVR) and non-negative least squares (NNLS) estimators. A TV displacement above 0.1 means that reconciling the two history-specific estimates would move more than one-tenth of the total estimated composition, at identical bulk and terminal reference. Across the full evidence base (784 deletion checkpoints, 308,617 sample-by-deletion solutions, five estimators, 17 tissue-estimator groups), reference history changes the measurement itself, with estimator-dependent magnitude.

**Fig. 1.**
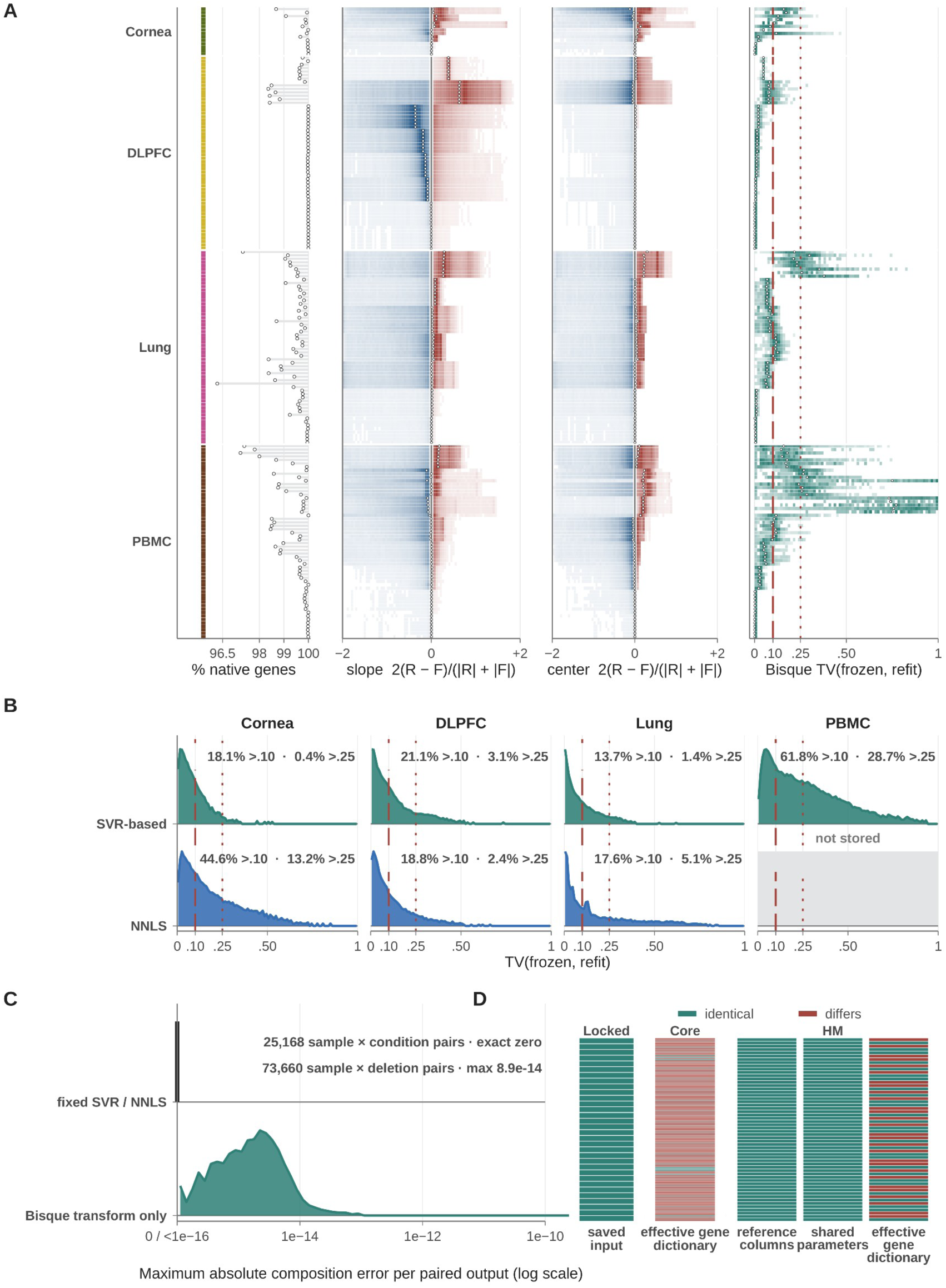
Estimator-state scars persist at identical reduced endpoints. **(A)** Learned-state change is shown across 182 tissue-by-source-by-deletion base units. “Same endpoint” means that the raw reduced reference and bulk mixture are identical; feature support and transformation belong to learned operator state and may differ between the frozen and refit paths. The operator-state summaries and output distances comprise 73,588 Aitchison-comparable sample-by-deletion pairs. **(B)** Total-variation (TV) distances between frozen and refit outputs are summarized within seven non-Bisque tissue-by-estimator groups, comprising 122,760 sample-by-deletion pairs. Across all 11 groups with stored vectors, 196,420 pairs were available; 28.2% exceeded TV 0.10 and 8.7% exceeded TV 0.25. These pooled percentages are not the denominator for the seven within-group displays. **(C)** The commuting control gives exact output identity for 25,168 paired outputs when the operator is fixed. The Bisque transformation-only comparison instead contains 73,660 composition-error pairs; this denominator is distinct from the Aitchison-comparable set in A. The displayed errors quantify numerical composition agreement, not componentwise scalar counts. **(D)** Endpoint-state tiles report identical or differing saved input, effective gene dictionary, reference columns and shared parameters across three audit sets with their own denominators, which are never pooled: 32 locked-control conditions (terminal dictionary and saved-input identity; 32/32 identical), 364 core-deletion conditions (the 182 tissue-by-source-by-deletion base units of A, each audited under two dictionary-matching modes; frozen-versus-refit terminal dictionary; 18 identical and 346 differing) and 56 Huuki-Myers imaging-benchmark conditions audited on three items each (shared parameters, gene dictionary and reference columns; 140 identical and 28 differing out of 168 item-level checks). Teal and brick red in D denote identical and differing states, respectively. Elsewhere, the blue-to-brick diverging fields encode the signed *R− F* relative slope and center contrasts shown on their axes and do not denote path superiority. HM, Huuki-Myers; NNLS, non-negative least squares; SVR, support-vector regression.

Two controls isolate the source of non-commutation. A controlled branch for the calibration estimator keeps the gene space, basis and bulk fixed and re-learns only the transformation after deletion. It reproduces the native refit output to numerical identity (median Aitchison distance 10⁻¹⁴; 99.99% of samples below 10⁻⁸, a compositional-distance criterion distinct from the per-component maximum error reported in Fig. 1C). Conversely, a commuting negative control that pre-locks every learned component predicts F ≡ R exactly, and delivers it: the maximum frozen-refit difference was numerically zero in all 32 checked conditions. In the two controls, re-learning of operator state was the only difference channel, and deletion arithmetic alone contributed nothing. Larger drift in the learned transformation accompanies larger output displacement: across the 182 tissue-by-deletion units of the calibration estimator, the Spearman correlation between the relative drift of the learned calibration slope and the frozen–refit Aitchison distance was 0.82–0.91 within tissue.

“Identical terminal reduced reference” means the raw reduced reference and bulk are fixed. Feature support, transformations and scaling are defined as components of the operator state and are therefore permitted to differ between paths where the estimator learns them. We substantiated this boundary with an endpoint comparison of the estimator outputs. Terminal reference columns, sample dictionaries and locked parameters are identical between paths in every evaluated condition (56/56 in the imaging-benchmark experiment; 32/32 in the commuting control, where the full input identity holds byte-for-byte). The effective gene dictionary, by contrast, differs between paths in 28/56 imaging-benchmark conditions and 173/182 evaluated core base units, which is exactly where estimators filter or select features as part of the learned state.

### Reference memory travels through distinct carriers and survives real edits

Reference memory resides in distinct carriers, and ordinary reference edits preserve the resulting state difference. Support-level auditing, the standard way to ask whether a reference change was consequential, asks whether the marker genes survived. By this audit, deletion appears benign.

Tissue-balanced median native-gene retention after deleting a type ranged from 99.63% to 99.99% (96.28–100% across individual deletion units). The learned operator shows the opposite. Deleting one type renormalizes each donor’s composition over the retained types, on which Bisque’s calibration targets are built, and the learned per-gene affine map moves substantially.

Depending on tissue, 35–61% of genes change their calibration slope by more than 10% (intercepts: 18–58%). The slope drift is analytically equal to the drift of the calibration-target dispersion, and the calibration center drifts independently of the dispersion (their median relative drifts differ by a factor of 19 in DLPFC). The operator moves in two dimensions while the marker list barely moves at all. Near-complete feature retention therefore does not establish operator stability.

Across estimators, we found reference history stored in at least three distinct carriers (Fig. 2). In BisqueRNA it lives in continuous, reference-learned calibration parameters, and re-learning only the transformation reproduces the rebuilt output to numerical identity. In MuSiC, reference history is carried by the bulk-dependent state generated during iterative fitting [23]. In identical-support deletions (17,794 genes after MuSiC’s internal filtering), the mean-expression basis, cell sizes, cross-subject variances and donor sets are bit-identical between the frozen and rebuilt paths. Holding that basis fixed and recomputing the solver state reproduces the rebuilt output exactly (residual 0). The magnitude of this endogenous refit is itself deletion-dependent: deleting inhibitory neurons barely perturbs it (maximum prediction change 3×10⁻⁵) while deleting astrocytes rewrites it (≈0.33). In the DNA-methylation pipeline, history is stored discretely, through feature-support turnover under IDOL-style selection (identifying optimal DNA methylation libraries) [25]. These are estimator-specific carriers of the same abstract failure: equality of the terminal reference matrix does not imply equality of the measurement state.

**Fig. 2.**
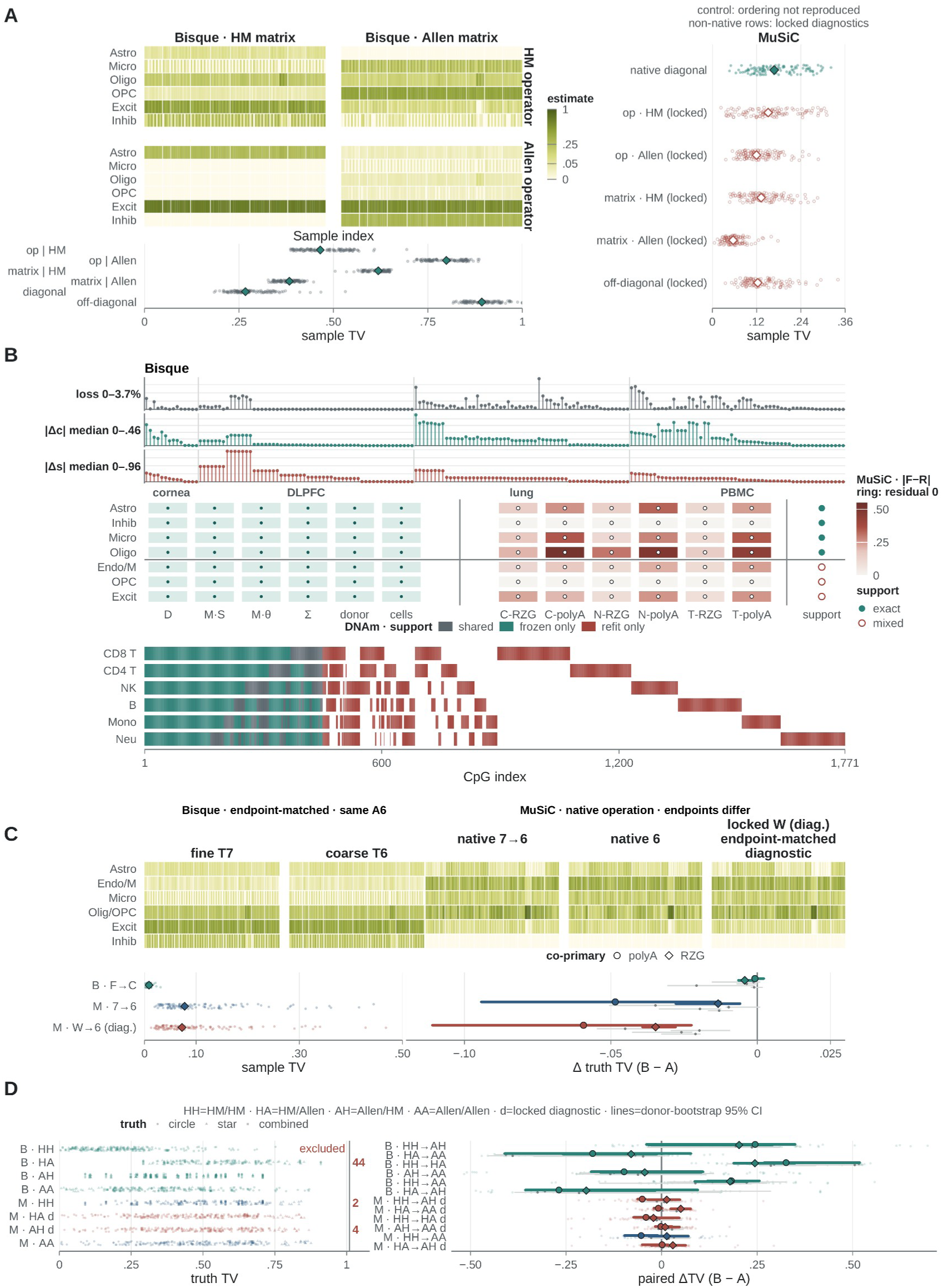
Reference operations expose estimator-specific state dependence. **(A)** Huuki-Myers and Allen reference matrices are crossed with their learned operator states on the same 110 libraries. The Bisque matrix-by-operator square displays observed total-variation (TV) separations. MuSiC native diagonal cells are operational comparisons; its locked-weight off-diagonal cells are mechanism diagnostics, not official native paths. The co-adaptation ordering observed for Bisque is not reproduced by MuSiC. **(B)** Three operator-state carriers are shown: Bisque transformation and centering change, MuSiC support and residual-weight behavior, and DNA-methylation feature support. The DNA-methylation display contains 1,771 unique CpGs across 4,809 deletion-by-CpG memberships, with 450 CpGs per arm per deletion; unique features, memberships and per-arm counts are not interchangeable denominators. **(C)** Merge operations separate an endpoint-matched Bisque comparison from a MuSiC native operational comparison whose endpoints differ and a locked-weight endpoint-matched diagnostic. Horizontal contrasts use *Δ* truth TV (*B− A*), so positive values mean greater truth loss for operation B than A. **(D)** Cross-atlas transfer compares two independent reference systems, Huuki-Myers and Allen, rather than successive versions. Bisque uses the full operator-source-by-reference-matrix square; MuSiC off-diagonals remain locked-weight diagnostics. Lines are donor-bootstrap 95% confidence intervals. Factor-swap contrasts are read one swap at a time. Color keys are panel-local and are not a single figure-wide scale. In the B DNA-methylation support strip, gray, teal and brick red denote shared, frozen-only and refit-only support; in the B Bisque traces the two colors separate center drift from slope drift; in the B MuSiC support matrix, filled and open symbols denote exact and mixed support. In C–D, colors distinguish the displayed native, endpoint-matched and locked diagnostic operations rather than a universal winner. C, cytoplasmic; CpG, cytosine–phosphate–guanine site; cpg_index (panel B), CpG index; Endo, endothelial; HM, Huuki-Myers; Mono, monocyte; N, nuclear; Neu, neutrophil; NK, natural killer; op, operator; RZG, RiboZeroGold; T, total.

The memory survives the reference operations practitioners actually perform. In a merge operation on the imaging-benchmark data, we rebuilt the identical six-type terminal reference by relabeling oligodendrocytes and oligodendrocyte precursor cells (OPC) as one type and rebuilding. For Bisque, the endpoint-matched contrast between seven-type-learned and six-type-learned operator states gives a median TV separation of 0.009. For MuSiC, the native operational comparison yields 0.078, and a locked-weight counterfactual on the byte-identical six-type reference produces a separation of the same order (0.073), so the inherited weight state alone suffices to generate the divergence.

Cross-atlas transfer crossed the donor-matched benchmark reference with the official Allen Institute DLPFC atlas (Allen region label DFC) [30] on a strict six-type common core. Bisque operator-source swaps at a fixed matrix produced median TV separations of 0.47–0.80, and matrix swaps at a fixed operator state 0.38–0.62. These are factor-swap contrasts, read one swap at a time. Fully crossed combinations reached 0.89. The two matched native Bisque systems were closer to one another (median TV 0.267) than any mismatched hybrid was to its native counterpart (0.383–0.799). This ordering is consistent with partial co-adaptation between each reference matrix and the operator state learned from it. MuSiC’s native diagonals differ by 0.168, its off-diagonal cells are locked-weight mechanism diagnostics, and no co-adaptation inference is drawn from MuSiC. A reference system behaves as a coupled measurement instrument, matrix and learned state together. Comparing composition estimates across independent reference systems, or across versions when an update changes both matrix and learned state, is therefore a cross-instrument comparison, shown here by direct measurement on two independent systems.

### Against orthogonal measurement, path advantage depends on the deleted type

Neither operator history is consistently closer to orthogonally measured composition; which one wins depends on the deleted type. We evaluated both paths against our strongest external benchmark: the Huuki-Myers human DLPFC multi-assay resource, in which cell composition was orthogonally measured by RNAScope/immunofluorescence (IF) on sections of the same tissue blocks that were bulk-sequenced [31]. The resource comprises 110 post-quality-control (QC) RNA-seq libraries from aliquots of 19 tissue blocks nested within 10 donors, six library preparations, and a donor-matched 56,447-nucleus single-nucleus RNA-seq (snRNA-seq) reference.

Neither path was consistently closer (Fig. 3). Of 24 prespecified stratified contrasts (6 deletions × 2 estimators × 2 co-primary strata; nominal 90% donor-bootstrap intervals), 13 excluded zero: 10 favoring the frozen path, 3 the rebuilt path. The largest contrast sits within a single estimator and preparation (Bisque, Total×RiboZeroGold): deleting astrocytes favors the frozen path (median paired TV difference +0.044, 90% confidence interval (CI) 0.035 to 0.050). Deleting inhibitory neurons instead favors the rebuilt path (−0.065, 90% CI −0.091 to −0.015). Under max-|t| simultaneous adjustment, 9 of 24 contrasts remain nonzero at 90% (8 frozen-favoring, 1 rebuilt-favoring) and the inhibitory-neuron interval individually crosses zero. A direct astrocyte-versus-inhibitory contrast, specified post hoc after the family-level analysis, is positive with donor-bootstrap 95% CIs excluding zero in both co-primary strata (0.116, 95% CI 0.009 to 0.134; 0.068, 95% CI 0.007 to 0.123).

**Fig. 3.**
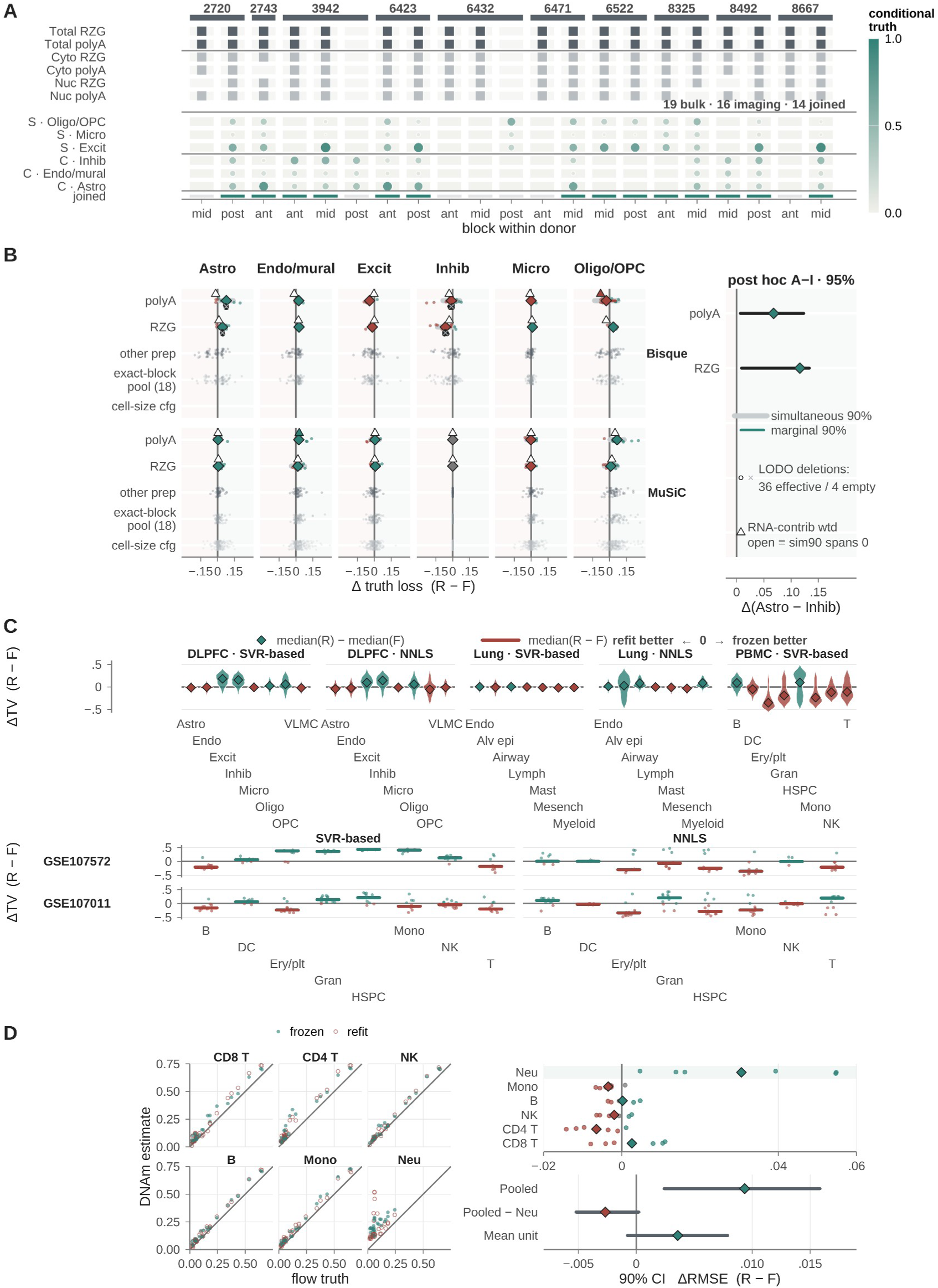
Truth-benchmarked path advantage changes with deletion and estimand. **(A)** The Huuki-Myers imaging benchmark aligns conditional composition measurements with bulk preparations by block within donor. The 110 post-quality-control libraries are aliquots of 19 blocks nested within 10 donors, not independent truth replicates. **(B)** Frozen–refit truth-loss contrasts are *Δ* = *R− F*: positive values indicate lower loss for frozen, whereas negative values indicate lower loss for refit. Sample/block pairings are the observational units and donors the resampling clusters; ‘other prep’, ‘exact-block pool (18)’ and ‘cell-size cfg’ rows are sensitivity rows (other library preparation, exact-block-matched 18-block reference pool, cell-size weighting). Astrocyte and inhibitory-neuron deletions have opposite nominal signs, but the inhibitory-neuron simultaneous 90% confidence interval spans zero. The Astro-minus-Inhib contrasts were post hoc and equal 0.116 [0.009, 0.134] for Total×RiboZeroGold and 0.068 [0.007, 0.123] for Total×polyA, using donor-bootstrap 95% confidence intervals. They support a deletion-dependent difference in path advantage rather than two separate family-level effects on opposite sides of zero. Under RNA-contribution weighting, all four headline simultaneous intervals span zero. **(C)** Truth-loss contrasts across independent synthetic-mixture and flow-cytometry benchmarks, *R− F* direction retained; the difference-of-medians and median-of-paired-differences summaries are distinct estimands. **(D)** DNA-methylation estimates against flow truth, with 90% confidence intervals for *Δ R M S E* = *R− F*; the overall direction depends on the pooled versus neutrophil-conditioned estimand. Color keys are panel-local: in C and in the directional summaries of B and D, teal denotes frozen-favoring and brick red refit-favoring contrasts, gray intervals or states selecting neither path; the B line key distinguishes simultaneous from marginal 90% intervals; the D scatter key identifies the frozen and refit estimates and does not encode direction. Alv epi, alveolar epithelial; cfg, configuration; CI, confidence interval; DC, dendritic cell; Endo, endothelial; Ery/plt, erythroid/platelet; Gran, granulocyte; HSPC, hematopoietic stem and progenitor cell; LODO, leave-one-donor-out; Mesench, mesenchymal; Mono, monocyte; Neu, neutrophil; NK, natural killer; op, operator; RMSE, root-mean-square error; RZG, RiboZeroGold; wtd, weighted.

The family-level evidence therefore establishes the frozen-favoring astrocyte effect and a deletion-dependent difference in path advantage, with the inhibitory-neuron effect individually unresolved. As a prespecified sensitivity, re-weighting the benchmark by cell size preserves the sign of 20 of 24 contrasts but widens all four headline intervals to cross zero. The pattern is thus established for the prespecified primary estimand, unweighted conditional cell composition, and attenuates under RNA-contribution weighting. Direction is a joint property of deleted type, estimator and assay: across the six preparations, the Bisque contrast reverses between strata for five of six deletions, MuSiC for two of six.

The same heterogeneity replicates in every other truth domain available. In three constructed-truth domains (1,500 Dirichlet pseudo-mixtures each for PBMC, lung and DLPFC, built from cells held out of the reference), the sign of the rebuilt-minus-frozen error difference varies across deleted types in all five domain-estimator combinations. Freezing is better for some deleted types and refitting for others, within every combination. In two independent flow-measured blood cohorts (GSE107572, nine samples; GSE107011, twelve donors, mapping prespecified before comparison with flow cytometry), both estimators split across the eight deletions under the unified paired-difference estimand. The deletion-level sign agrees between the two cohorts in only five of eight (SVR-based) and four of eight (NNLS) conditions.

In whole-blood DNA methylation with an independent flow-characterized whole-blood test set (36 sample-deletion units), the ranking is estimand-dependent. The pooled root-mean-square error (RMSE) difference favors freezing (+0.0094, 90% CI +0.0024 to +0.0158, complete cluster enumeration) but is carried entirely by the neutrophil deletion. Excluding it reverses the point estimate. The per-unit win count leans the other way (refit 18, frozen 14, ties 4). Declaring the paths interchangeable would require equivalence bounds of 20.9% and 10.4% of baseline RMSE for the two estimands. Under either estimand, the data support neither a fixed direction nor demonstrated equivalence, and the reversal pattern is domain-dependent.

### Operator history reverses cohort associations

Operator history changed the sign of 160 of 952 association estimates across nine cohorts (Fig. 4). The cohorts are non-longitudinal disease cohorts in four tissues (952 paired frozen–refit associations; tissue-matched references; two-sided Wilcoxon, Benjamini–Hochberg (BH) false discovery rate (FDR) within each path). Of the 160 opposite-signed pairs, 31 were significant on at least one path, and 126 pairs changed significance status. The ten-cohort tally (1,064 pairs; 180 opposite-signed; 33 significant reversals; 144 status changes) is reported descriptively, because the tenth cohort’s repeated draws invalidate independent-sample inference (Methods). In the most extreme case, in the SVR-based analysis, monocyte/macrophage proportions associate with COVID-19 positively on the frozen path (FDR = 0.030) and negatively on the rebuilt path (FDR = 0.006): two significant, opposite conclusions from identical data and an identical reduced reference. Among the 37 path-inconsistent associations in the cornea cohort, the built-in discovery/replication split gives 24 concordant signs, 12 reversals and one tie on the frozen path. The rebuilt path gives 12 concordant signs, five reversals and 20 ties (Fig. 4D). Operator history changes the sign-replication summary as well as the association estimate. The path difference itself is statistically supported by direct between-path inference.

**Fig. 4.**
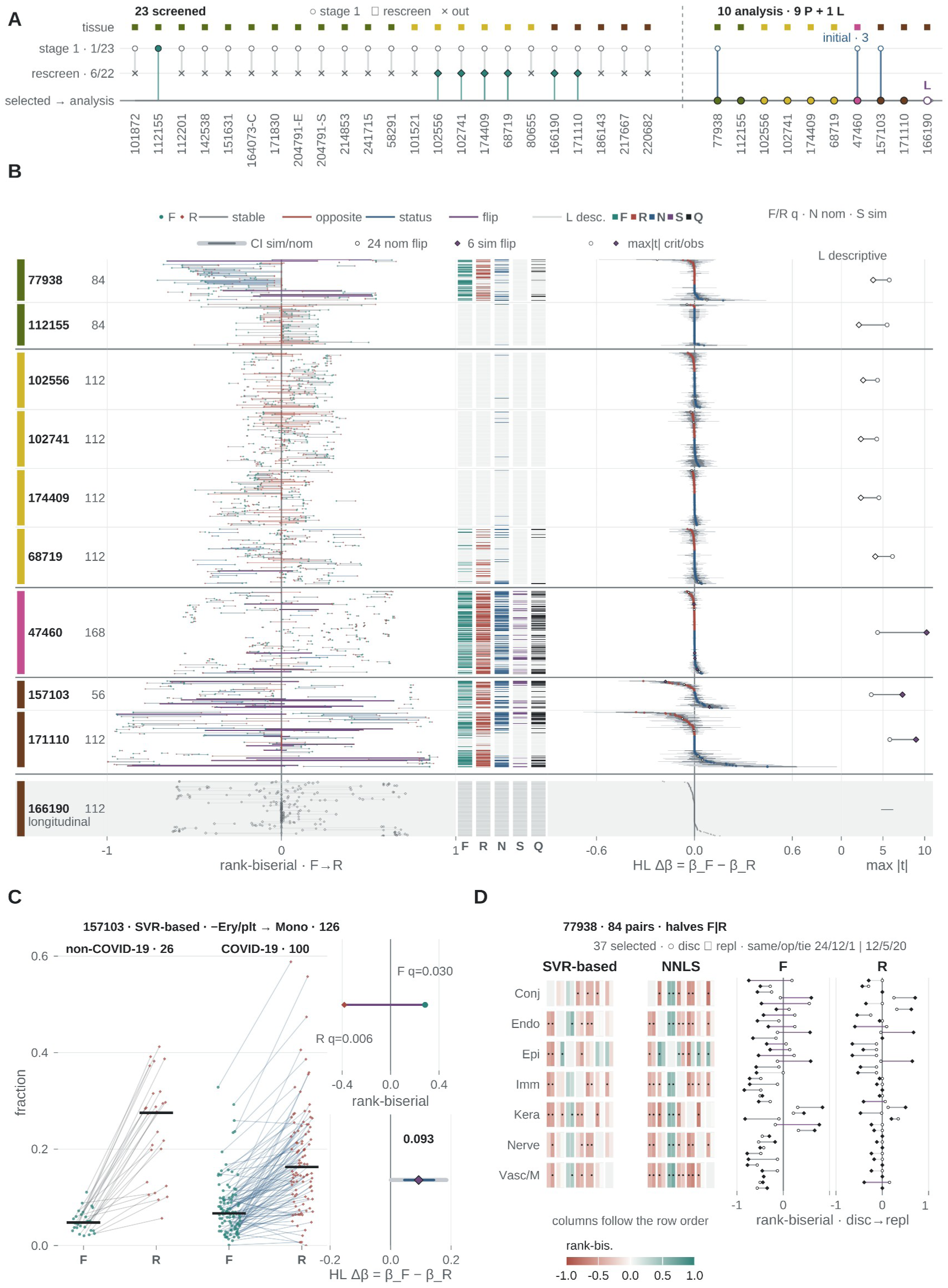
Association direction and significance depend on operator history. **(A)** Two-stage cohort screening identifies the analysis set under prespecified eligibility rules. **(B)** Frozen and refit association effects and between-path inference are shown for 1,064 association IDs, partitioned into the 952 primary pairs from the nine non-longitudinal cohorts and the 112 longitudinal pairs from GSE166190, which are reported descriptively and do not enter primary family counts. For the 952 primary pairs, *Δ β* = *β_F_ − β_R_*: 220 nominal 95% intervals excluded zero; 24 of these pairs had opposite-signed path estimates. Separately, 160 tests passed global Benjamini–Hochberg correction, 177 passed within-cohort Benjamini–Hochberg correction, three of nine cohort-level omnibus tests rejected, and six opposite-signed pairs retained nonzero simultaneous 95% intervals. These are distinct nominal, global, within-cohort, omnibus and simultaneous decision rules. **(C)** The GSE157103 monocyte example, shown for the SVR-based analysis, has opposite frozen and refit association directions and a nonzero rank-biserial [74] contrast. This unadjusted path-association comparison targets operator-history dependence. **(D)** The corneal discovery–replication display compares path-specific sign consistency among selected association pairs and does not add a cohort to the primary universe. Color and shape keys are panel-local. Teal circles and brick-red diamonds denote frozen and refit effects in the panels that display path-specific estimates. In the D heatmaps, teal denotes positive and brick-red negative association effects, and purple connectors mark discovery–replication sign flips; panel B instead uses its own categorical line key for stable, opposite-signed, significance-status-change and sign-flip pairs. Continuous fields reproduce the displayed signed estimands, including *β_F_ − β_R_*, and are not a path-victory key. BH, Benjamini–Hochberg; Conj, conjunctival; disc, discovery; Endo, endothelial; Epi, epithelial; Ery/plt, erythroid/platelet; HL, Hodges–Lehmann; Imm, immune; Kera, keratocyte; Mono, monocyte; nom, nominal; repl, replication; sim, simultaneous; SVR, support-vector regression; Vasc/M, vascular/mural.

Estimating Δβ = β_F − β_R for every association pair (Hodges–Lehmann location shift; subject-level stratified bootstrap), 220 of 952 nominal 95% intervals exclude zero, 24 of them with opposite-signed point estimates. Under global and within-cohort BH correction, respectively, 160 and 177 between-path tests remain significant. Three of nine cohorts reject a vector-level omnibus null, and 6 of the 24 opposite-signed pairs retain nonzero within-cohort simultaneous 95% intervals. Across the 392 association pairs of GSE47460, GSE157103, GSE77938 and GSE112155, the absolute between-path effect shift |Δβ| was 0.52 times the frozen-path bootstrap standard error at the median and 3.9 times at the 90th percentile. Operator history therefore moves estimated associations on the scale of sampling uncertainty and, in the upper tail, several times beyond it. Prespecified two-stage screening of the 23 remaining cohorts admitted seven (one in stage 1 and six in stage 2 under a metadata amendment applied uniformly). Each was analyzed under the identical protocol, with every inclusion and exclusion decision reported in Additional file 1: Table S2. The longitudinal cohort GSE166190 enters only as a supplementary analysis (Additional file 2: Fig. S2; Additional file 1: Table S3).

Deletion effects also need a correct null for where a deleted type’s signal lands among the retained types (Fig. 5). This structural layer asks which retained types receive the deleted signal, and the conventional null assumes the signal lands uniformly across receivers. Replacing the uniform receiver null with a null conditioning on the realized receiver margins cut significant structural-layer units from 12 to 3 (104 units; no unit gained significance). Empirical type-I error assessment confirms anti-conservatism rather than a mere difference in stringency: under no-effect simulations consistent with the realized margins, the conventional uniform null rejects at 12.1% against a nominal 5%, reaching 26–34% in individual tissue-estimator cells. The margin-aware null is conservative at 2.2%. Across nominal levels the calibration slopes are 1.87 versus 0.45. At these margins, the conventional omission null inflates significance calls.

**Fig. 5.**
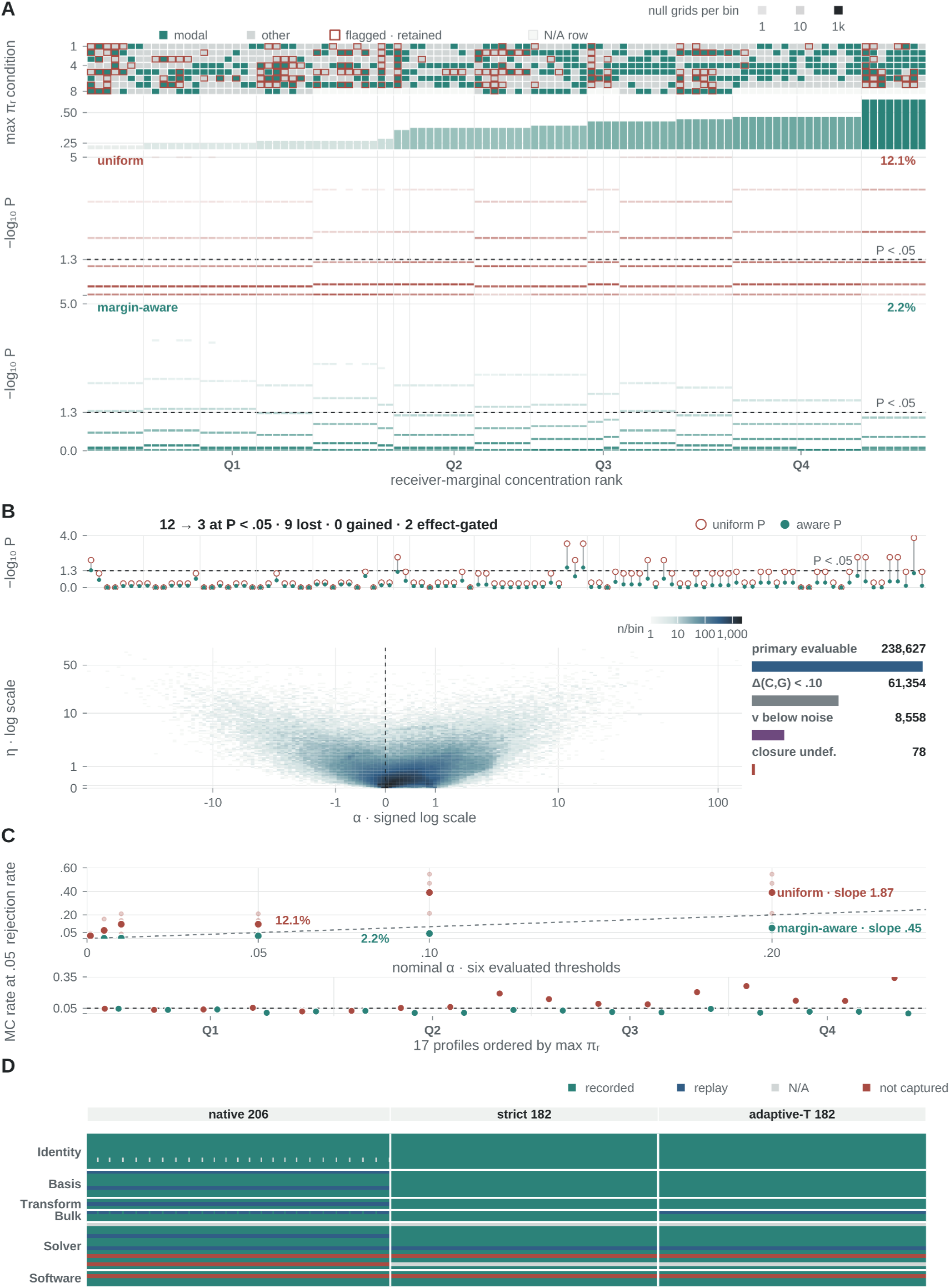
Uniform receiver nulls inflate false positives to 12.1%, and margin-aware recalibration is conservative. **(A)** Structural receiver margins and Monte Carlo null *P* values are aligned by source unit. The 104 source units reuse 17 receiver profiles and therefore do not represent independent profile estimations; each null comprises 104,000 Monte Carlo grids, and the uniform and receiver-margin-aware nulls reuse the same grids. **(B)** Recalibration changes nominal calls from 12 to 3 at *P* < 0.05, with no gained calls and cases excluded by the effect-size threshold (‘effect-gated’ in the panel) shown separately. The lower field maps the evaluable primary grid by signed *α*, scale and observation density; it is a calibration surface, not additional independent replication. **(C)** At nominal 0.05, empirical rejection is 12.12% for the uniform null and 2.17% for the receiver-margin-aware null. Across the evaluated thresholds, the fitted calibration slopes are 1.87 and 0.45, respectively; both are intercept-inclusive fits, not through-origin slopes. Unit-weighted quartiles and profile-wise Monte Carlo rates remain descriptive summaries of the prespecified empirical receiver-margin data-generating process. **(D)** The Bisque reference-state manifest covers 570 operator objects across 38 fields and serves as the Bisque-specific worked reporting exemplar among the five estimators. Recorded, replay-derived, not-applicable and not-captured fields are displayed separately; in particular, N/A is not merged with not captured. Color keys are panel-local. In the null and P-value elements of A–C, brick red denotes the uniform null and teal the receiver-margin-aware null; the A structural matrix instead separates modal from flagged-and-retained source units, and D uses its own four-state key for recorded, replay-derived, not-applicable and not-captured fields. Teal intensity in A and density shading in B encode the displayed receiver-margin or bin-count scale, not frozen/refit path superiority. MC, Monte Carlo; N/A, not applicable.

### The full-reference target is not point-identified, and the two failures are independent

Even with the operator fixed, the same observed bulk and incomplete reference support different retained-composition targets. Distinct missing-type profiles reconstruct the same bulk exactly while reversing the retained-type ranking, so the ambiguity attaches to the target itself, and Theorem 1 below strengthens the reversal to maximal non-identification (Fig. 6).

**Fig. 6.**
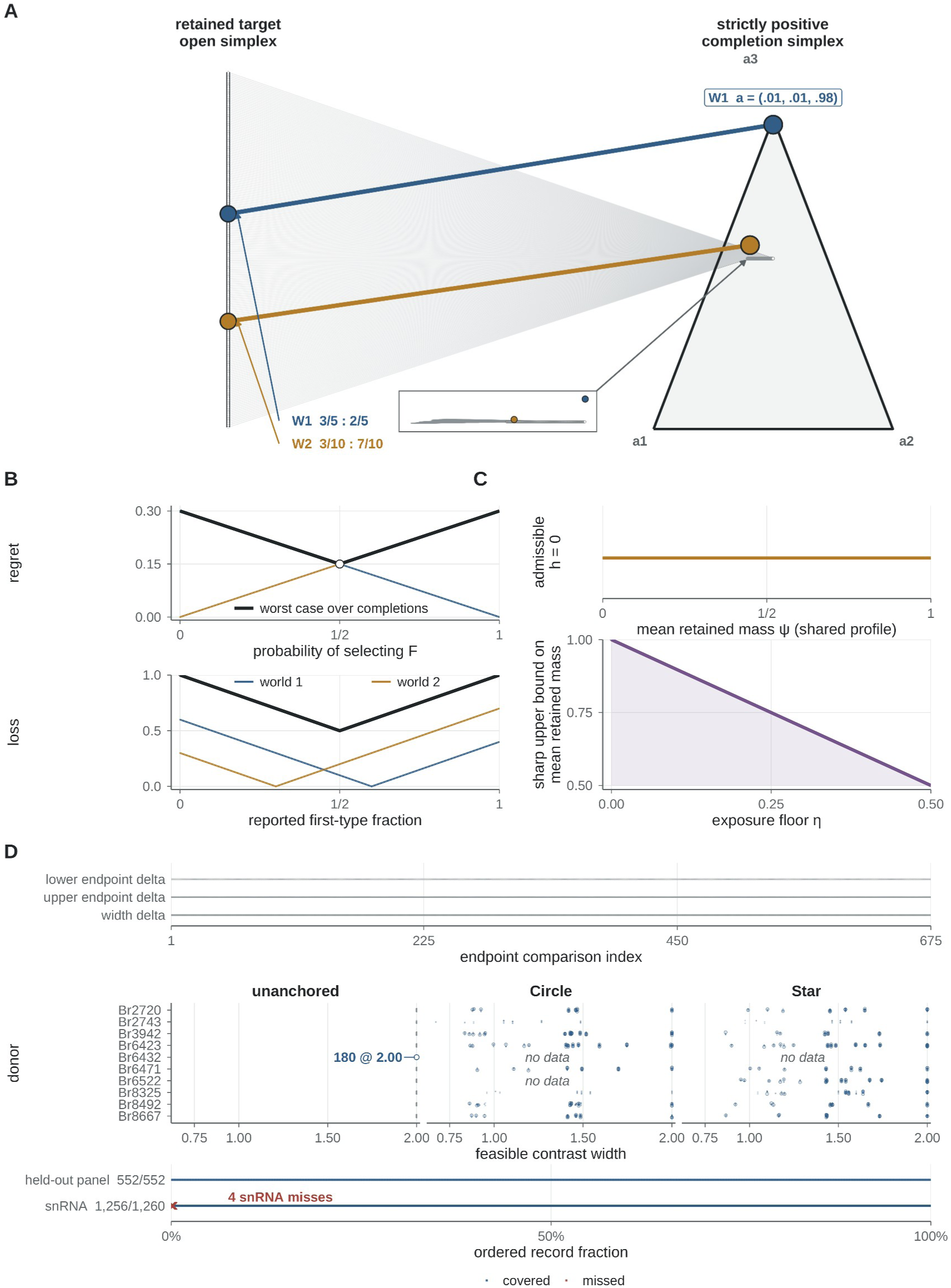
The retained-composition target is not point-identified, and neither shared structure nor additional views repair it. **(A)** Two admissible completions of the same strictly positive observation (*P* = 3, *K* = 2) reverse the retained-type ranking, (3/5, 2/5) versus (3/10, 7/10), both reconstructing the bulk exactly with simplex-interior solutions. The identified set fills the open simplex via a strictly positive family with nonsingular completed references (*κ*_1_ *≤* 1313/485 < 2.71); opposite rankings persist on relative *ℓ_∞_* neighborhoods of radius 1/200. **(B)** No observed-data rule universally selects the better candidate: at *δ* = *d_TV_* = 3/10, minimax regret over the full completion class is *δ* for deterministic binary selectors and *δ*/2 for randomized ones (upper plot, probability of selecting F: *δ* at either endpoint, *δ*/2 at the midpoint); the full-class minimax worst-case loss over composition-valued outputs is 1/2 at the simplex midpoint (lower plot, reported first-type fraction; black, class-wide worst case). Eighty exact-arithmetic checks verify every displayed witness. **(C)** Under the joint zero-slice condition (Z), hidden exposure *h* = 0 annihilates the shared profile *a*, so cohort-sharing it, or fixing it by external calibration, tightens no bound inherited from the zero-exposure submodel (upper strip: every target stays admissible at *h* = 0). Deleting the zero slice is what moves an inherited bound: in the scalar model 1 = *x_i_* + *h_i_*, an exposure floor *h_i_ ≥η* lowers the sharp upper bound on the mean retained mass from 1 to 1 *−η* (lower plot). **(D)** In Huuki-Myers blocks, mean feasible coordinate widths from one, two, three and six library preparations are identical at 1.0000, as constructed by the prespecified class with a separate positive-diagonal operator for every block-by-view observation, whereas one three-type imaging panel used as a partial anchor contracts the width to 0.7604 with 100% held-out coverage. Coordinate widths (0–1, donor-balanced over 11 blocks and 8 donors) and per-block contrast widths (0–2) are on their own scales. TV, total variation.

The construction is displayed twice, in two parameterizations that share the same conclusion. For readability we state it on the plain standard-basis pair *A_−d_* =[*e*_1_ *, e*_2_] with a bulk profile *y* =(3/10, 2/10, 5/10) on three features. Fig. 6A and the theorem itself use the smoothed pair *A_−d_* =[*S e*_1_ *, S e*_2_], with *S* =(97/100) *I* +(1/100)11*^⊤^* (columns (98, 1, 1)/100 and (1, 98, 1)/100), at *y* =(301, 204, 495)/1000. The smoothed reference is strictly positive, and the strict-positivity, rank and conditioning guarantees of Theorem 1 attach to it. Both yield the same two retained-type targets and the same reversal.

For the standard-basis pair, two completions, appending column (0, 0, 1) or appending column (3/10, 1/10, 3/5), are both admissible: non-negative, unit-sum, and reconstructing y exactly with simplex-interior solutions. Under the first completion the retained-type conditional composition is (3/5, 2/5). Under the second it is (3/10, 7/10). The ranking of the two retained types reverses. All quantities are exact rationals. No noise, estimator, or operator refit is involved.

The example is robust to perturbation, and we establish it as a theorem, with an exact-arithmetic script checking every displayed witness and constant (80 checks) and the universal claims proved analytically (Additional file 2: Note 1; Additional file 3: Data S1). **Theorem 1 (maximal non-identification).** There exists a strictly positive observation with P = 3 features and K = 2 retained types whose identified set of retained-type conditional compositions is the *entire open simplex*: every target composition (t, 1−t) with 0 < t < 1 is attained by an explicit strictly positive realizing family a(t), each of whose completed references is nonsingular with 1-norm condition number at most 1313/485 < 2.71, and opposite rankings persist on relative ℓ∞-open neighborhoods of radius 1/200. The strictly positive two-world ranking-reversal construction embeds into every dimension setting P ≥ 3, K ≥ 2. When P ≥ K+1, a separate embedding can additionally be chosen with full column rank. Point identification thus fails as completely as it can fail, with positivity, rank and conditioning all controlled. **Theorem 2 (no universal observed-data selector).** At that observation, with q_F = (3/5, 2/5) and q_R = (3/10, 7/10) as the two operator histories’ candidates and δ = d_TV(q_F, q_R) = 3/10: the minimax regret over the full admissible completion class (not merely over two chosen worlds) is exactly δ for deterministic binary selectors and exactly δ/2 for randomized ones; the minimax oracle-relative regret over composition-valued outputs is likewise exactly δ/2, attained at the midpoint (it is the minimum over outputs, not a value every output attains), while the full-class minimax raw worst-case risk over composition-valued outputs equals exactly 1/2, attained at the simplex midpoint, because the identified set fills the simplex. Because the ambiguity is algebraic, arising at the completion step itself, replication of the same observed-data law cannot remove it under the unrestricted model. Resolution requires additional information or restrictions: constraints on the missing profile, informative multi-sample structure, or an anchor reference.

Conditionalization standardizes the reporting scale. It does not identify the target. A sum-to-one output on the observed reference can at most target the conditional composition, never absolute proportions and hidden mass simultaneously, so re-normalizing onto retained types is necessary for any comparison. The witness shows that even the ranking on this common scale reverses across admissible completions.

The two failures are logically independent, and neither implies the other. A fixed, externally supplied NNLS operator commutes with deletion by construction, as our pre-locked control demonstrates, yet the completion ambiguity remains untouched. Conversely, an adaptive operator is path-dependent even in benchmarks where the target is externally measured and thus identified for evaluation purposes, as the imaging benchmark shows. The regime in which both failures operate, an adaptive estimator with an incomplete reference and no external truth, is the regime the methodological reviews describe [2, 3]. This 2×2 structure also fixes the scope of each claim: operator provenance concerns estimators with reference-learned state. Completion non-identification is broader and applies to every reference-based method, fixed or adaptive. The two failures also compound, and Theorem 2 is their formal connection: once two operator histories emit different candidates at the same observed input, unknown completion can reverse which candidate sits closer to the fixed estimand. Truth can validate a path locally, within a measured tissue, preparation and deletion, but no observed-data rule can be uniformly correct over an unrestricted completion class.

Theorem 2 already rules out a universal selector. We used a synthetic deletion spectrum to ask the narrower, average-case question of whether common diagnostics discriminate the lower-error history under one generating distribution. In this spectrum, built to avoid oracle circularity (400 dictionary instances × 6 deletions; mixtures generated from K+2 columns while the estimator saw only K), direction heterogeneity persisted (71% frozen-better, 29% refit-better). None of five candidate diagnostics (deleted-column distance to the retained hull, inter-column coherence, deleted-column weight, operator drift magnitude and hidden mass) discriminated the lower-error path (area under the receiver operating characteristic curve, AUC, 0.445–0.574; Additional file 2: Fig. S1).

### Replication and additional views are not identification

Repeated samples and additional uncalibrated views, under the completion and operator classes stated below, leave different target values compatible with the same observations. This ambiguity is the gauge, and only information that breaks the underlying observational equivalence can contract the identified set. Theorem 2 names the escape routes: constraints on the missing profile, multi-sample structure, an anchor. Replication of the same kind, samples or assays observed through the same uncalibrated map, adds observations without adding the information that separates admissible completions; the Proposition below states exactly when it adds nothing. That is why the operative response has been to model the unknown component and return a point estimate [8, 11].

#### Proposition (shared completion structure)

Consider multiple samples constrained to share a single completion profile a. If the admissible class satisfies the joint zero-slice condition (Z), under which every retained composition in the base identified set admits a completion with joint zero hidden exposure, h = 0, then the shared-profile constraint tightens no bound inherited from the zero-exposure submodel. At h = 0 the product h·a annihilates a: even fixing the missing profile exactly, by external calibration, adds nothing (Additional file 2: Note 3). Under (Z), constraints on the missing profile alone remove no target value that is feasible at zero exposure; a bound inherited from the zero slice moves only when they are paired with information that deletes that slice. A per-sample exposure floor h_i ≥ η > 0 is one sufficient example (in the scalar model 1 = x_i + h_i it lowers the sharp upper bound on the mean retained mass from 1 to 1 − η), as are cross-sample average-exposure lower bounds, hidden-type-specific anchors, and absolute-mass constraints. A PBMC pilot evaluates both sides. Under a model class consistent with (Z) at its zero-exposure-floor setting, sharp bounds remained exactly [−1, 1] with zero contraction, and remained so at positive floors, an additional empirical failure the Proposition does not predict. A rank-one shared-structure class produced feasible witnesses of both signs; across the two pilot classes, 273 of 384 optimization extrema were solved, 107 remained unresolved at the prespecified gap and 4 were infeasible, and all 36 main-profile extrema of the rank-one class remained unresolved.

Additional measurement views obey the same logic on the operator side. When each block-by-view observation carries its own unknown positive diagonal operator and no constraint ties these operators across views or blocks, the operators absorb any completion, and the identified set cannot contract. The Huuki-Myers resource realizes this exactly (Fig. 6). On the same tissue blocks, feasible-width bounds computed from one, two, three or all six library preparations are identical at width 1.0000, with bound-by-bound differences exactly zero, the consequence the operator class predicts. Adding a single three-type imaging anchor on the same blocks instead contracts the mean coordinate width to 0.7604 (24.0% contraction) with 100% held-out coverage. Identification requires an independently constrained measurement map, and no number of measurements substitutes for one. Independently calibrated views can constrain composition even when each view alone leaves it ambiguous. We next asked whether observable-fit budgets and finite external-profile libraries supply identifying restrictions of this kind or merely stabilize estimator outputs.

### Contraction without coverage is false certainty

Observable fit leaves both association signs feasible in every one of the 952 primary contrasts, and the external profile library narrows estimator outputs without covering the continuity oracle. Every restriction is therefore scored jointly on shrinkage, coverage, certification and false certification against a declared oracle, with results from different oracle families reported separately. Between the theorem’s unrestricted completions and any imported information stands the restriction every practitioner already has: observable fit. We fixed per-sample tolerances τ_i before inspecting results. Each tolerance is the sample’s full-reference model-misfit floor (its best-achievable total-variation reconstruction residual) plus one history-independent slack term, the larger of a held-out-gene residual and a constructed-truth pseudobulk floor (λ = 1; Methods). We then computed sharp linear-programming (LP) feasible-effect intervals for the Hodges– Lehmann association effect [32] over all 952 cohort × analysis-unit × deletion × retained-type contrasts of the nine non-longitudinal cohorts. All 44,856 per-sample LP bound calculations completed without solver failures. The exactness proof is in Additional file 2: Note 2.

Observable fit identifies the direction of none of the 952 association effects: in every contrast the feasible interval contains both signs. The budget itself is the obstruction: across the nine cohorts the median model-misfit floor alone is 0.33–0.65, approaching the maximal total variation of 1, and the full budget exceeds that maximum for 783 of 937 samples (83.6%). Observable fit, calibrated additively on absolute TV, never becomes discriminating in these cohorts.

A prespecified library of biologically anchored completions sharply contracts estimator-output envelopes where the budget could not. For every tissue and deleted type we completed the reference with each eligible type-matched external donor pseudobulk profile, re-ran the estimator under the prespecified refit-on-completed strategy, and recorded the finite-library effect envelope, the min–max hull of the resulting association effects. Of 2,049 profile fits, 46 ended in solver failure. Across the 952 contrasts, 390 (41%) are library-conditionally sign-invariant, meaning that every evaluated profile leads the estimator to the same association direction. A further 95 (10%) have an evaluated library sign flip, 345 have an evaluated near-zero effect, 56 lack a type-exact library, and 66 are incompletely evaluated (Additional file 4: Data S2). Under the finite-library assumption we first read that 41% as usable evidence. Under the four-quantity criterion, that 41% measures agreement among estimator outputs over the evaluated profiles, and whether such agreement supports a sign conclusion is decided by coverage and false certification.

The criterion is that added information must be scored on four quantities at once: the shrinkage ratio (how much the interval contracted), coverage (how often the contracted interval still contains its oracle), certification (how often it excludes zero) and false certification (how often it excludes zero on the wrong side). They are scored against three oracle families that are never merged. Constructed truth (known generating proportions) scores point-estimator error only, since the estimator outputs there carry no matching interval bounds. Orthogonal external measurement truth is flow cytometry. The full-reference continuity oracle, the effect recovered when the deleted column is restored, is a consistency target, not independent biological truth.

Scored this way, the library envelope contracts by 98.93% on average yet covers the continuity oracle for only 42.77% of evaluable contrasts (355/830). Its 390 certificates are 41.0% of the 952 planned contrasts and 47.0% of the 830 oracle-evaluable ones, and 19 of them carry the wrong sign (2.0% of planned contrasts, 2.3% of evaluable contrasts, 4.9% of certificates; Fig. 7). The identification benchmark on the same data is unambiguous. Conditional sharp bounds (the identified set given the linear mixture model, the prespecified budget and the library-derived completion class) are the complete [−1, 1] at all 30 prespecified scan points of library size × inflation: shrinkage 0, coverage 100% (all 934 evaluable contrasts at the main setting; 18 unavailable), certificates 0. The 41% is therefore a conditional phenomenon of finite estimator-output sensitivity, a fact about which profiles the library happens to contain, and not a property the identified set supports. Nor can its robustness to profile provenance be assessed on these data: 210 of the 390 invariant contrasts (53.85%) rest on a single profile source. For the remainder, a leave-one-source-out check is vacuous by construction: the envelope is a min–max hull, so any complete subset of an all-same-sign union is necessarily same-sign, and deleting one source can only preserve, never falsify, an invariant call. The asymmetry is visible directly: deleting one source leaves all 392 evaluable states unchanged, deleting the other changes 58, and none of the 58 changes starts from the sign-invariant state.

**Fig. 7.**
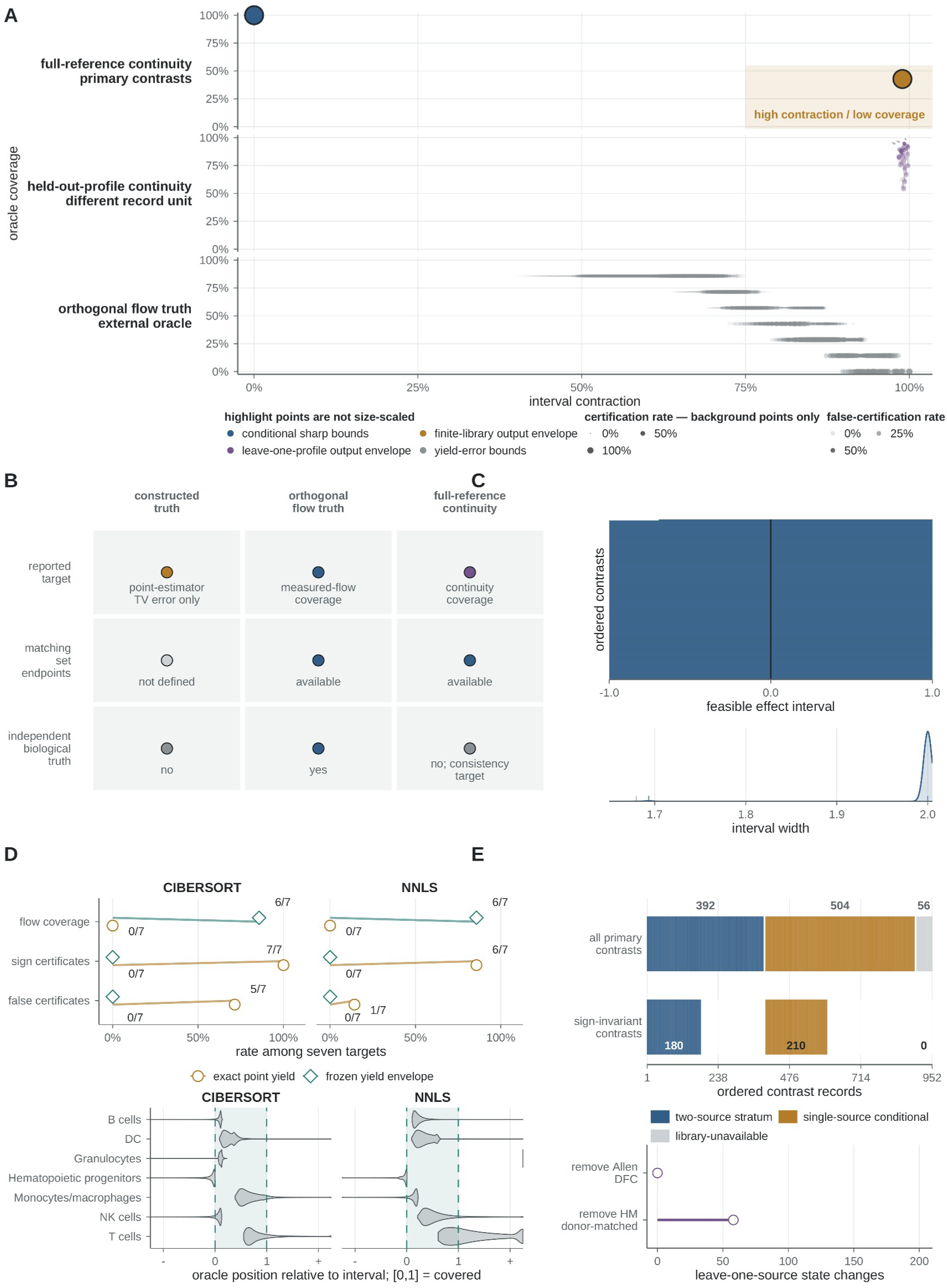
Contraction without coverage is false certainty. **(A)** Contraction–coverage plane. Background point size encodes certification rate; false certification has a separate key; highlighted points are not size-scaled. The finite-library output envelope contracts effect intervals by 98.93% yet covers the full-reference continuity oracle for only 42.77% of evaluable contrasts (355/830); its 390 certificates are 41.0% of 952 planned and 47.0% of 830 evaluable contrasts, and 19 carry the wrong sign. Conditional sharp bounds on the same data sit at zero contraction, 100% coverage (934/934 evaluable) and zero certificates. The leave-one-profile-out frontier, whose record unit and oracle differ from the primary contrast, traces the trade from 66.76% coverage with 48.60% certification (library size two, 11,691 records) to 90.66% with 41.32% (size eight, 20% inflation, 11,229 records). **(B)** Three separate oracle families: constructed truth scores point-estimator error only; orthogonal flow cytometry is external measurement truth; full-reference continuity is a consistency target. **(C)** With prespecified tolerances, all 952 primary contrasts admit both signs, the nine-cohort median model-misfit floor is 0.33–0.65 against a maximal total variation of 1, and the full budget exceeds that maximum in 783 of 937 samples. **(D)** With a flow oracle, treating per-type yields as exact contracts the interval by 100% and certifies 6/7 (NNLS) and 7/7 (CIBERSORT) targets with flow coverage 0/7, five CIBERSORT certificates pointing the wrong way (false certification 71.43%); re-inflating to the prespecified envelope (‘frozen yield envelope’ in the panel) restores coverage to 6/7 and sends all certificates to zero. **(E)** Sign-invariant contrasts: 210 of 390 (53.85%) rest on one profile source; for the remainder a leave-one-source-out check is vacuous, since a min–max hull over an all-same-sign union stays same-sign on any subset. Deleting one source changes 0/392 evaluable states, the other 58/392, none from the sign-invariant state.

A cleaner demonstration uses the absolute-scale system, where an orthogonal flow oracle exists. Treating measured per-type RNA yields as exact collapses the feasible interval to a point (shrinkage 100%) and produces sign certificates for six of seven (NNLS) and seven of seven (CIBERSORT, the SVR-based signature estimator; Methods) downstream targets, each a mapped type’s cell fraction minus its RNA contribution, taken as the median across donors. Against the flow oracle, coverage is 0 of 7 for both estimators, and five of CIBERSORT’s seven certificates point the wrong way (false certification 71.43%). Re-inflating the yields to their realistic prespecified error envelope restores coverage to 6 of 7 and sends certificates, true and false alike, to zero for both estimators. The point-yield precision came from excluding the truth. Structurally, exact yields do close the proportion-versus-RNA-contribution gauge, and the mechanism works as designed. Externally, the current yield measurements are miscalibrated enough that the closed gauge points at the wrong answer. Even exact yields act on the conversion, not on the completion: giving both witness worlds the same yields leaves their observations identical and their retained targets different, for every positive yield vector (Additional file 2: Note 1, Corollary N1.1).

On the leave-one-profile-out envelope, which uses a different record unit and oracle from those underlying the 42.77% result above, expanding the library and inflating tolerances buys coverage at modest certification cost. Coverage rises from 66.76% with 48.60% certification at the smallest library setting (11,691 evaluable held-out records) to 90.66% with 41.32% at library size eight with 20% tolerance inflation (11,229 records). Any completion method should report certification–coverage frontiers of this kind alongside shrinkage.

### Three ceilings, and the measurement frontier

Identification, model fit and estimator boundary zeros impose three distinct ceilings on what these data can certify (Fig. 8). The precision frontier prices the measurement that would lift the first, and the other two belong to the model and to the estimator. The **identification ceiling** is set by the information itself: conditional sharp bounds are [−1, 1] throughout, so the sign-certificate ceiling of the library class is zero, and any certificate the envelope emits is estimator-conditional. The **model ceiling** is set by how far real mixtures sit from the linear model and reference class. Its diagnostics live on different scales and are reported item-wise, never compressed into one number. Across the 12 PBMC donors the median full-reference relative RMSE is 0.200 and the residual-to-signal L1 ratio 0.462; across the six Huuki-Myers library preparations the median reconstruction TV tolerance is 0.393. Across the ten-cohort diagnostic panel of 1,035 samples the median marker-fit floor is 0.558, the held-out-gene residual 0.596 and the pseudobulk floor 0.519.

**Fig. 8.**
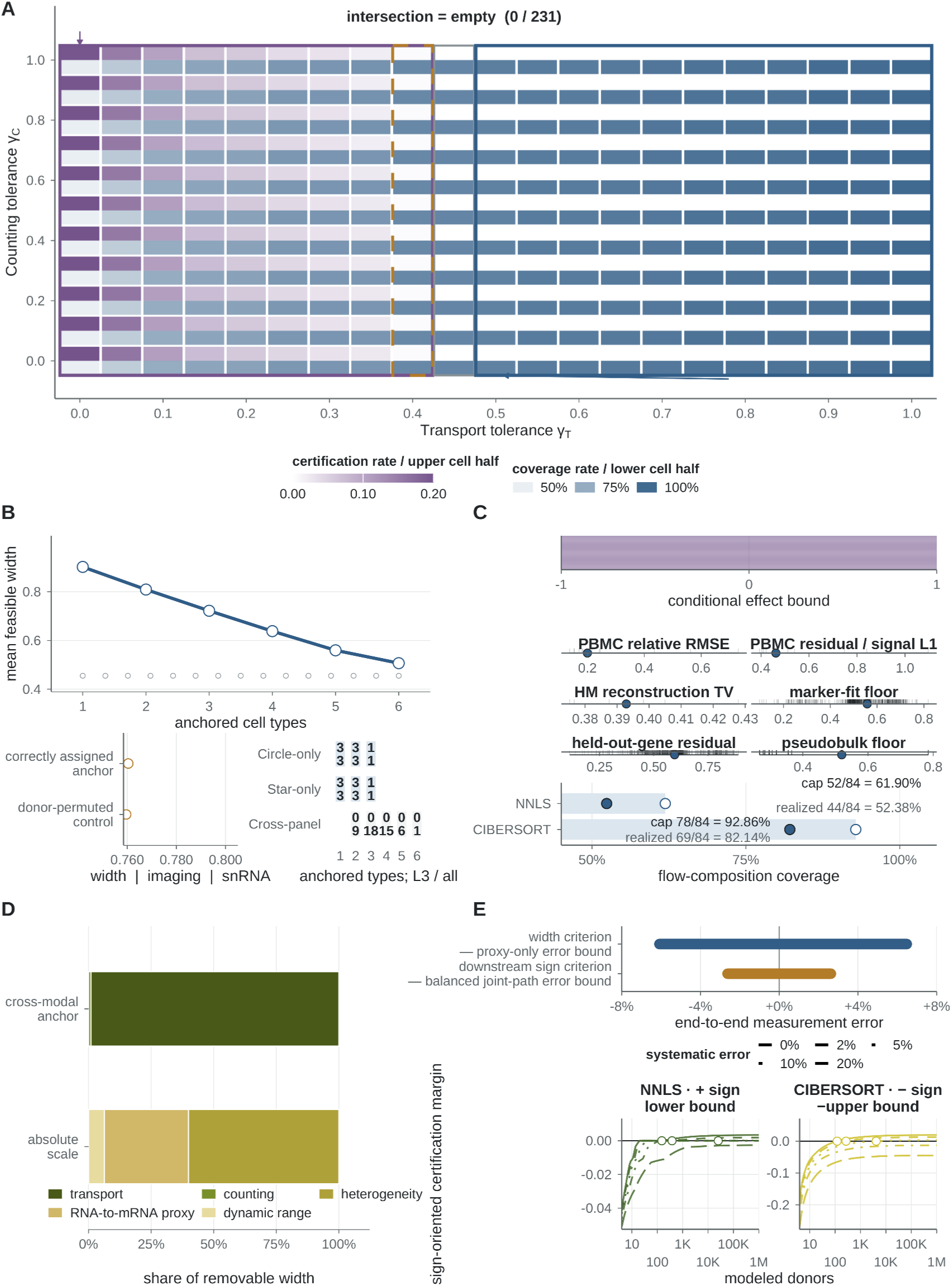
Three ceilings, and the price of measurement precision. **(A)** Cross-modal anchor certification over the 231-point transport × counting grid: no cell meets both thresholds. At full counting tolerance (*γ_C_* = 1), the first raw certificate requires *γ_T_ ≤* 0.40, where snRNA coverage (0.8837) already fails the 90% threshold; certification and coverage trade directly along that axis (0.1972/0.5104 at *γ_T_* = 0; 0/0.9201 at *γ_T_* = 0.50). **(B)** Anchoring more types contracts the feasible width monotonically (0.9022, 0.8094, 0.7221, 0.6383, 0.5595, 0.5067 for one to six types on the common roster of 7 blocks and 6 donors), while at full tolerance all 15 pairwise sign certifications are zero. The donor-permuted control contracts to 0.7596 against 0.7604 for the assigned anchor, with imaging coverage undiminished and snRNA coverage 0.9931. Only single-panel combinations of up to three types retain a complete complementary validation panel; types are ordered by width reduction (record-level results in Additional file 2: Fig. S4). **(C)** Three ceilings: identification (conditional sharp bounds are [*−* 1, 1] at all 30 scan points, so the certificate ceiling is zero), model (item-wise residual diagnostics, medians marked) and algorithmic (frozen NNLS fixes 32 of 84 composition cells at zero, capping flow-composition coverage at 61.90%, 52.38% realized; CIBERSORT fixes 6 of 84, cap 92.86%, realized 82.14%). **(D)** Shapley attribution of removable width: 99.01% cross-modality transport and 0.99% counting error; in the absolute-scale system 60.1% donor/leaf heterogeneity, 33.6% total-RNA-to-mRNA proxy, 6.3% dynamic range. **(E)** Precision frontier: the width criterion is met at proxy error +6.5%/−6.1% with other sources at main settings; fixing the sign along the equal joint-scaling path requires ±2.6% with 97.4% of the log-scale heterogeneity and dynamic-range envelopes removed. Modeled donor requirements are 151, 381 and 25,168 (NNLS) and 120, 269 and 4,152 (CIBERSORT) at 0%, 2% and 5% systematic error, unattainable at 10%; the estimators certify opposite signs. NNLS, non-negative least squares.

The **algorithmic ceiling** is the estimator’s own. With the identical 637-gene space and leaf mapping, frozen NNLS fixes 32 of 84 composition cells at exactly zero, capping achievable flow-composition coverage at 61.90% (52.38% realized). CIBERSORT fixes only 6 of 84, raising the cap to 92.86% (82.14% realized). Changing the estimator therefore moves the algorithmic ceiling (by 26/84 structurally, 25/84 realized) without moving the identification ceiling at all: a better estimator recovers coverage lost to boundary zeros, and it supplies none of the missing identification information. The ceilings do not share coordinates: the precision thresholds below live on measurement-error axes and cannot be plotted against the 61.90%/92.86% coverage caps.

Calibrated cross-modal anchors are the one information class that contracts while retaining coverage. On the Huuki-Myers common roster (7 blocks, 6 donors), anchoring types one at a time contracts the feasible width monotonically, from 0.9022 with one anchored type through 0.8094, 0.7221, 0.6383 and 0.5595 to 0.5067 with six (Additional file 2: Fig. S3C). Yet at the full prespecified tolerance range all 15 pairwise sign certifications are zero. With the counting tolerance kept at its full value (γ_C = 1), tightening the assumed transport tolerance produces raw certificates for γ_T ≤ 0.40, reaching a rate of 0.1972 at γ_T = 0 and 0.604 for individual type pairs. At γ_T = 0.40 the aggregated held-out snRNA coverage is 0.8837, below the 90% coverage threshold. Along the transport axis, certification and coverage trade directly (0.1972 certification with 0.5104 coverage at γ_T = 0; 0 with 0.9201 at γ_T = 0.50).

Across all 231 grid cells, none meets both the certification and coverage thresholds. Both thresholds are evaluated after aggregation across samples, donors and all 15 type-pair identities. Individual sample-by-pair records that are simultaneously certified and covered do occur (129 of 62,370) but are neither broad enough across type-pair identities nor consistent enough across donors to satisfy the grid-cell validity criteria after aggregation (Additional file 2: Fig. S4). The marginal-value ordering of anchor types (EndoMural > Micro > Inhib > OligoOPC > Astro > Excit) is a width-reduction ordering on this roster, not biological importance (Additional file 2: Fig. S3). A Shapley decomposition [33] of the removable interval width attributes 99.01% to cross-slide, cross-modality transport and 0.99% to counting error. The binding constraint is the fidelity with which a section-level measurement transports to the bulk aliquot.

Donor misassignment preserves anchor contraction and held-out imaging coverage. A donor-permuted anchor control, with the same anchor-width multiset and entirely misassigned donors, contracts the width to 0.7596 against 0.7604 for the correctly assigned anchor, a difference below 0.001. Held-out imaging coverage does not drop, and snRNA coverage falls only from 1.0000 to 0.9931. Contraction and coverage therefore distinguish narrower from wider feasible sets and leave misassigned anchor information undetected: on this resource, a benchmark scoring only those two quantities would accept the same contraction after every donor assignment had been made wrong. The anchoring depth that the strongest oracle class can verify is also structurally capped. Only single-panel combinations of up to three types leave a complete complementary imaging panel to verify against, so the narrower cross-panel and six-type combinations are exactly the ones that cannot be verified.

Absolute quantification is the one class that closes the proportion-versus-RNA-contribution gauge in principle, and its price can be stated as a specification. Under the current PBMC measurement class, the prespecified width ratio under the main settings is 0.7786 and the downstream T-cell sign target, cell fraction minus RNA contribution, crosses zero. Inverting the prespecified uncertainty model yields the requirements. The width criterion (ratio ≤ 0.50) is reachable by tightening one source: total-RNA-to-mRNA proxy error within +6.5%/−6.1% while the other sources remain at their main settings. Fixing the downstream sign requires near-point joint information: along the prespecified equal joint-scaling path, proxy error within ±2.6% while 97.4% of the current donor-heterogeneity and dynamic-range envelopes, on the log scale on which they are parameterized, is simultaneously removed; with those two envelopes idealized to points, the proxy alone would still need to be within +5.4%/−5.2%. The bottleneck ordering is measured: Shapley width contributions are 60.1% donor/leaf heterogeneity, 33.6% total-RNA-to-mRNA proxy and 6.3% dynamic range, so the measurement with the highest expected information yield is direct mRNA-per-cell measurement across donors, not deeper sequencing.

The two systems therefore demand different measurement improvements: transport from section to aliquot in the cross-modal system, donor heterogeneity and the total-RNA-to-mRNA conversion in the absolute-scale system.

For NNLS, modeled donor requirements for the sign target are 151, 381 and 25,168 donors at 0%, 2% and 5% end-to-end systematic error, reached when the lower bound rises above zero, so the certified sign is positive. CIBERSORT reaches its own sign target at 120, 269 and 4,152 donors, but by the opposite route: its upper bound falls below zero and the certified sign is negative. The two donor counts therefore price opposite conclusions and are not interchangeable. The NNLS figure of 25,168 states what 5% systematic error costs in donors, and at 10% neither estimator reaches its sign target at any donor count. The donor frontier thus separates a sample-size requirement from a systematic-error limit: below the limit, recruitment can make an estimator-specific sign decisive, and above it no recruitment can.

### A measurement contract, and what it costs to satisfy it

The measurement contract is executable today, and fitdrop implements it together with the precision frontier. The contract has four clauses (manifest exemplar in Fig. 5D). First, **operator-state reporting**: an adaptive reference is closer to a trained model checkpoint than to a static matrix, so a composition estimate should carry a reference-state manifest. The manifest lists source accessions and cell ontology, terminal retained types, the edit history of the reference, selected feature set, learned transformations and weights, software versions [34], and a hash of the serialized operator state [35]. Second, **endpoint-matched sensitivity analysis**: when a reference edit is part of an analysis, run both operator histories at the fixed endpoint and report their disagreement, exactly as one would report a batch-sensitivity analysis. Third, **conditional compositions as the common scale, with the completion class declared**: re-normalize onto retained types and state the class over which conclusions are claimed, because the witness shows that the conditional target itself is not identified without one. Fourth, **equivalence bounds for claims of interchangeability** [36]: report the bound the data support [37] (here, 20.9% and 10.4% of baseline RMSE).

**fitdrop 0.2.0** implements the contract and the frontier. It emits the operator manifest, the per-deletion spectrum of frozen–refit differences with its sign map, and the supportable equivalence bound for any reference-estimator pair. It also provides `precision_frontier()`, a callback engine that inverts a user-supplied uncertainty model into system-specific precision requirements: single-source thresholds, equal-proportion joint thresholds, Shapley width attribution and donor-N curves. Its core contains no system-specific code. Two vignettes, the

PBMC absolute-scale analysis and the Huuki-Myers transport analysis, reproduce their headline numbers from the released engine (width threshold κ = 1.0648, joint width threshold u = 0.6426, joint sign threshold u = 0.0263; 231-point grid, raw-scale γ_T* = 0.40, transport share 99.01%). The interface serves any system that can supply a compliant `eval_bounds` callback, parameterize its uncertainty sources as interval envelopes, and satisfy the monotonicity or grid-only conditions (Methods). Estimator boundary zeros are flagged as an algorithmic ceiling and excluded from the inversion.

## Discussion

An incomplete reference fails twice to define the analysis: it need not define a unique estimator, and it need not identify a unique target. Non-commutation is the first failure (same endpoint, different estimator): the identical reduced reference yields different compositions depending on the history of the operator state, because the reference is training data as well as design matrix. Non-injectivity is the second (same observation, different target): observationally equivalent completions disagree about the quantity being estimated. The striking observation is that the *same* reduced reference gives different answers depending on when the operator was learned from it, and that even perfect knowledge of the observed system would not settle which answer points at the truth. Resolving that ambiguity takes information that meets a precision specification, and the four-quantity criterion scores whatever information a method supplies.

A conventional confidence interval on a composition estimate captures sampling uncertainty, conditioned on one operator history and one completion assumption [38, 39], neither of which the interval carries. We identify the two conditioned-away layers: operator-provenance uncertainty (the spread of estimates over valid operator histories ending at the same endpoint) and completion uncertainty (the spread of targets over admissible completions). In the cohort analyses, moving between two valid operator histories was sufficient to reverse significant associations, a shift that per-path sampling intervals, each conditional on its own operator history, cannot reveal.

Prior omission studies carry out heterogeneous reference edits [6, 15, 16, 23]. None we found crosses frozen and rebuilt operator states while holding the observed terminal endpoint fixed, and a modular framework [24] capable of expressing the contrast does not report it. Empirically, the bias caused by missing types is well documented [6, 15, 16]. Formal identification theory sits in adjacent territories: necessary and sufficient conditions for identifying the mixing matrix in blind separation of well-grounded sources [40], feasible-set (rotational-ambiguity) analysis of non-negative factorizations in chemometrics [41, 42], calibrated sampling uncertainty under correct model specification [43], and partial identification as a general inferential frame [44–46].

Residual-based recovery operates with the number of missing types known and reports an empirical one-to-one match between the recovered component and a factor, with no identifiability guarantee [15]. Our witness shows the unknown-completion regime is where uniqueness fails. The reference-choice sensitivity literature varies the reference [19, 20]. Our design holds it fixed and varies only its history.

The provenance results apply to any estimator whose operator state depends on the reference, and across estimators we found this state traveling through at least three distinct carriers. The first carrier is static calibration parameters learned from reference-derived pseudo-bulk. In Bisque, re-learning only the transformation reproduces the rebuilt output to numerical identity. The second is the bulk-dependent state generated during iterative fitting. In MuSiC’s identical-support deletions, every static learned component is bit-identical between paths and the asymmetry is localized to the iterative re-solution. The third is discrete feature-support turnover (DNA-methylation IDOL-style selection). Marker selection whose scores depend on the full column set, calibration learned from reference-derived pseudo-bulk, and reference-dependent weights are all instances. Across five estimators, non-commutation is pervasive but varies in magnitude and direction.

Completion non-identification is broader and binds every reference-based method, including fixed-operator ones. Theorem 1 itself is an existence construction at an explicit observation. It establishes that nothing in positivity, rank or conditioning forbids maximal ambiguity. How far a given observation’s set extends is what the conditional sharp bounds measure.

Each external benchmark measures a specific quantity, and the history comparisons are read against that quantity. Constructed-truth mixtures share donors (not cells) between reference and mixtures, so donor-level effects are common to both arms. Flow-sorted RNA has nine samples, and flow fractions are cell fractions while deconvolution without cell-size correction estimates RNA-mass fractions [47], a systematic gap shared by both arms and not path-specific, although it need not cancel exactly in the loss contrast. Orthogonal imaging benchmarks composition. Each probe panel labels three of six broad types (composition is conditional within panels), sections pass morphology quality control, and its authors explicitly caution that deconvolution may track RNA contribution rather than cell numbers [31]. That caution is why we prespecified unweighted conditional cell composition as the primary target and cell-size weightings as sensitivities.

The constructive endpoint is invariance. Practitioners instinctively ask which path is correct. The imaging benchmark shows the answer changes with deleted type, estimator and assay, and the witness shows no observed-data selector can be uniformly right over an unrestricted completion class. The productive question is which conclusions (ranks, effect directions, phenotype associations) remain invariant across valid operator histories and admissible completions. Where a conclusion is invariant, it is robust to the two hidden inputs. Where it is not, external information must resolve it, and that information can be priced.

The evaluated information classes differ sharply in what they buy. Replication and additional uncalibrated views leave these ambiguities unchanged (the Proposition, and the exact-zero multi-view instantiation). A constraint on the missing profile excludes no zero-exposure witness, so it moves an inherited bound only when paired with information that forces the hidden type’s exposure to be positive, which sharpens the escape-route list in Theorem 2. External profile libraries sharply narrow estimator-output envelopes, but the evaluated library did so at the expense of coverage. The envelope measures the spread of the library. Calibrated cross-modal anchors are the first class that buys width, yet on the best available resource they never buy a valid sign certificate, and the anchor configurations narrow enough to matter are exactly the ones the strongest oracle cannot verify. Absolute quantification closes the proportion-versus-RNA-contribution gauge in principle, at a measured price that differs by system and does not transfer between them. None of these classes validly certifies a sign today, but the criterion and the specification apply to each of them.

Benchmarking an estimator and benchmarking the information supplied to it are distinct tasks. Error against a measured target ranks histories within a setting and does not establish identification. The donor-permuted control shows that even contraction with held-out coverage need not verify sample-specific information: changing every donor assignment left the interval width essentially unchanged and imaging coverage undiminished, so the benchmark validated narrowing but could not validate alignment. An identification benchmark must therefore distinguish informative restrictions from incorrect restrictions that also narrow the feasible set.

Methods that recover missing profiles from residual structure [15] or model unknown components under explicit assumptions [8–10] make two distinct claims, that their outputs are accurate under a declared model and that their added restrictions support narrower conclusions, and only the second is an identification claim. It is scored against a declared oracle by interval shrinkage, coverage, the fraction of oracle-evaluable targets that receive a sign certificate, and the fraction that receive a wrong-sign certificate. A method that reports its certification–coverage frontier is making a falsifiable claim about information. An estimator-output envelope measures sensitivity to the evaluated completions, conditional sharp bounds measure what the admissible class identifies, and reporting both prevents stability of outputs from being mistaken for adequacy of information.

The RNA and DNA-methylation results expose a measurement problem conditioned on the reference and independent of the assay. Reference history was carried by continuous calibration, by endogenous solver weights and by discrete feature selection, while completion ambiguity concerns which latent components the observations admit. Whenever a reference determines both the learned measurement state and the latent categories an estimator admits, reproducing the measurement map, identifying the target over the declared completion class and pricing the precision that would make the conclusion decisive are three separate tasks. Operator-state manifests and endpoint-matched histories address the first, conditional identified sets the second, and precision frontiers the third. This separation transfers to reference-based mixture estimators in other omics layers and does not depend on any one of the five implementations.

Local truth, the measured benchmark of one tissue, preparation and deletion, and restricted completion models rank operator histories only within their declared domains, because no observed-data selector is universally correct over the unrestricted admissible completion class. Comparisons across independent reference systems, or across versions that change both the matrix and the learned state, must demonstrate comparability [48].

When a remedy for incompleteness is proposed, ask for its four quantities against a held-out oracle. When new measurement is planned to resolve a composition question, compute the precision it must achieve first, which `precision_frontier()` does before any data are collected. A proportion vector or an atlas accession alone does not reproduce the measurement. Reproducibility requires a measurement contract: the observed bulk, the terminal reference and its ontology, the operator state and its history, the estimand and the completion class. Report which conclusions remain invariant across the declared operator histories and admissible completions.

## Conclusions

Two distinct failures of uniqueness attend an incomplete deconvolution reference. In adaptive estimators, rebuilding after a reference edit can alter the learned or endogenous measurement state even when the terminal matrix and bulk are fixed: fit-then-drop and drop-then-fit are different estimators wearing the same name. Which history lands closer to an external benchmark varies with the deleted type, the estimator and the assay. Separately, the retained-composition target is not uniquely identified by the observed bulk and incomplete reference over an unrestricted completion class: admissible completions of the same observations reverse retained-type rankings. The first failure invalidates default comparability across operator histories. The second means the canonical target requires a stated completion class. The second failure does not yield to more data of the same kind: under the stated completion class, shared completion structure and additional uncalibrated views add nothing, external profile libraries contract far more than they cover, and calibrated anchors contract without validly certifying a sign.

The criterion scores any proposed remedy on shrinkage, coverage, certification and false certification against each family of held-out oracles. The specification treats identification as a measurement problem with a computable price. Here, absolute quantification closes the proportion-versus-RNA-contribution gauge in principle. Fixing the downstream sign in the PBMC absolute-scale system requires roughly 2.6% end-to-end proxy accuracy together with near-total removal of the donor-heterogeneity and dynamic-range envelopes, and transport is the binding constraint of the separate Huuki-Myers cross-modal anchor system. Composition estimates should carry operator provenance, be interpreted under an explicit completion class, be stress-tested for invariance over the declared contract, and state equivalence bounds before any claim of interchangeability.

## Methods

### Estimators and operator states

Five reference-based estimators were run as implemented in the released analysis code. Non-negative least squares (NNLS) [49] ran with sum-to-one normalization. An SVR-based signature estimator (nu-SVR [50] over a nu grid, signature-wide z-scoring, no quantile normalization) is referred to throughout as CIBERSORT [27] because it is our re-implementation of that method. These results apply to the re-implementation, while the official CIBERSORT distribution applies quantile normalization. The reference-based branch of BisqueRNA (per-gene affine calibration learned from reference-derived donor pseudo-bulk) [26], MuSiC [23] and SCDC [51] complete the panel. Four of the five share a marker-selection stage whose per-column scores depend on all reference columns; BisqueRNA’s reference-based branch performs no marker selection, which makes it the cleanest isolation system for the calibration channel. For each deletion, the frozen path retains the operator state learned on the full reference (feature support, centers, scales, calibration targets, weights, hyperparameters) restricted to retained columns, and removes the deleted column only at the solve step. The refit path re-learns the full operator state from the reduced reference. The controlled intermediate arm for the calibration estimator re-learns only the transformation, holding gene space, basis, and bulk fixed. The commuting negative control pre-locks every learned component; its prediction F ≡ R was verified exactly (maximum difference 0) in all 32 checked conditions, using officially released estimator implementations. An endpoint audit records, per condition, the dimensions, ordering and hashes of the terminal reduced reference and bulk consumed by both paths. It also records whether feature support differed.

### Paired deletion experiments

A total of 784 deletion checkpoints (17 tissue-estimator groups; cornea [52–57], DLPFC [31], lung [58] and blood [59] sources; the blood panel is labelled PBMC throughout, its reference comprises eight classes including granulocytes and erythroid/platelet cells, and its cohorts are whole-blood or leukocyte preparations) yielded 308,617 deletion-level sample solutions (349,728 including full-reference runs). Frozen-refit distances are reported as Aitchison distances and as total variation (half-L1) on compositions, as labeled. An independent re-execution of the full pipeline reproduced all 54 reported quantities exactly.

### Imaging-benchmark evaluation (Huuki-Myers DLPFC)

The benchmark data comprised 110 post-QC bulk RNA-seq libraries from aliquots of 19 tissue blocks nested in 10 donors across six library preparations, a donor-matched snRNA-seq reference (56,447 nuclei and 7 broad types after removing ambiguous nuclei), and RNAScope/IF imaging composition at the block level [31] (two probe panels of three named types each; per-panel conditional composition after removing the residual `Other` category; no cross-panel six-type composition was constructed, and dual-panel blocks contribute the mean of the two panel losses). The entire analysis was prespecified in a dated decision record, released with the package, before any comparison with the imaging measurements. Ensembl [60] was the primary gene-ID scale (intersection 17,804). Oligo+OPC were merged to match the imaging panel, with single deletions of Oligo or OPC evaluated only for operator divergence (excluded from truth evaluation) and OligoOPC deleted jointly for truth evaluation. Primary truth was unweighted within-panel conditional cell composition, with nuclear-area and AKT3-based cell-size weightings as prespecified sensitivities. Co-primary strata were Total×RiboZeroGold and Total×polyA, with the remaining four preparations as replication strata. Official BisqueRNA 1.0.5 (`markers = NULL`, `use.overlap = FALSE`) and MuSiC 1.0.0 (`cell_size = NULL` primary) ran with the parameters specified in the dated decision record. The sample was the paired observational unit and the donor was the bootstrap cluster [61, 62], 10,000 resamples, fixed seed. The estimator was the median across samples of the paired TV difference (rebuilt − frozen). Two truth-only blocks without bulk counterparts were excluded by identity, not renamed. One sample in which the rebuilt MuSiC path assigned zero mass to all surviving panel types was excluded from both paths under a prespecified complete-pair rule (five scenario instances; all documented). Nine donors contribute paired evaluations (one truth-only donor contributes none). Family-level inference used max-|t| simultaneous donor-bootstrap intervals, Holm-adjusted bootstrap P values [63], an exact donor sign-flip global test over all 24 primary contrasts, and leave-one-donor-out re-analysis of the four headline cells [64, 65]. A post hoc direct astrocyte-versus-inhibitory contrast was specified after the family-level analysis (10,000 resamples, seed 20260807; family-level analyses used seed 20260808). No success criterion was set. Per-block and per-contrast tables underlying every number in the Results are included in the released outputs.

### Evaluations with constructed compositions and external measurements

For the constructed-composition evaluations, we generated 1,500 Dirichlet pseudo-mixtures per tissue (500 cells each) from a 50/50 cell-level split (mixtures from held-out cells; donors shared between arms). The first flow-measured RNA cohort was GSE107572, nine blood-cell mixtures (PBMC with 3– 6% admixed granulocytes from the same donor) with matched flow cytometry [12]; flow types mapped to six reference compartments (prespecified before analysis); estimates and benchmark re-normalized on mapped retained types before computing TV. The original prespecified estimand was the difference of medians, and the unified paired-difference estimand median(TV_R − TV_F) is additionally reported for cross-cohort comparison. The second flow-measured RNA cohort was GSE107011, 13 PBMC samples of which 12 had matched flow data after one source-study flow-QC exclusion, with bulk RNA-seq and raw flow proportions from the original publication [66]. Its 29 flow leaves were mapped to seven of eight reference compartments before comparison with flow cytometry (Erythroid/platelets absent from flow and excluded from evaluation classes); eight deletions, subject bootstrap with 10,000 resamples; cross-cohort agreement uses the unified estimand. For DNA methylation (DNAm), we evaluated IDOL-based whole-blood deconvolution [25, 67, 68], leave-one-type-out with frozen and refit arms, evaluated on an independent test set of six whole-blood samples with flow-measured proportions [68] (six deletion conditions, 36 units); cluster bootstrap by complete enumeration (6^6). For each analysis the estimand, metric, and decision criterion were prespecified in a dated decision record before results were inspected; the record is released with the package.

### Synthetic deletion spectrum

The spectrum comprised 400 dictionary instances (60 features, K = 6 visible plus M = 2 permanently hidden columns; gamma-distributed profiles with a shared-component coherence parameter; Dirichlet ground truth; multiplicative noise), with 6 deletions each. Both paths saw only the visible reference. Sign discrimination used AUC of five per-condition covariates.

### Two-completion witness and theorems

The witness and Theorems 1 and 2 are constructed in exact rational arithmetic; both completions are column-stochastic and reproduce the observation exactly with simplex-interior solutions. A released self-contained script performs exact-arithmetic checks of the displayed witnesses, constants, lattice points, the identified-set parametric family, and representative embeddings (80 checks); the universal neighborhood, dimension-extension, rank, identified-set and full-class minimax claims are established analytically in the released theorem note. The full construction and analytic proofs are provided in Additional file 2: Note 1; the exact-verification script and its full 80-check output are provided in Additional file 3: Data S1.

### Real-cohort associations

The primary cohorts were GSE47460 (474 lung samples analyzed here, two platforms) [69], GSE157103 (126 whole-blood leukocyte samples) [70] and GSE77938 (50 corneal samples; 16 discovery, 34 replication) [71]. For each path, we applied two-sided Wilcoxon tests [72] of estimated proportion versus phenotype per deletion × retained type, with BH-FDR [73] within each estimator × deletion × path family of retained types; frozen-refit pairs were compared on the sign of the rank-biserial effect [74] and on significance status, and an opposite-signed pair with at least one FDR-significant path is counted as a significant reversal. The cornea cohort’s discovery/replication split was used only for a prespecified sign-consistency summary of path-inconsistent associations, with no subcohort selection. The association analysis was covariate-unadjusted and targets operator-history dependence in the estimated associations. For the systematic sweep, all 23 remaining cohorts were screened under a prespecified inclusion rule (unambiguous binary primary phenotype per the series’ own design; n ≥ 20 and ≥ 8 per arm; tissue with an available reference). Screening ran in two prespecified stages: stage 1 used only the metadata distributed with the series files (one cohort qualified: GSE112155 [75]), and stage 2 ran under a prespecified amendment permitting official Gene Expression Omnibus (GEO) series-metadata retrieval [76], applied uniformly to all 22 prior exclusions (six qualified: GSE102556 [77], GSE102741 [78], GSE174409 [79], GSE68719 [80], GSE166190 [81], GSE171110 [82]). Each included cohort carries a phenotype manifest recording the metadata fields and mapping used; every exclusion carries a documented reason. All qualifying cohorts were analyzed under the identical association protocol with the tissue-matched reference. The repeated-draw structure of GSE166190 is documented; it is excluded from primary family-level inference, and the ten-cohort totals are reported descriptively.

### Reference-completion analysis

Sample-wise feasible-set analysis: per-sample feasibility classes *X_i_* ={*x ∈ Δ* : *d_TV_*(*y_i_, A_−d_ x*)*≤ τ_i_*} over the closed simplex *Δ*, with *τ_i_* the sample’s full-reference best-achievable reconstruction residual (model-misfit floor, r_full,i) plus a single F/R-independent slack term equal to the larger of a held-out-gene residual (genes split at random, prespecified seed) and the constructed-truth pseudobulk floor (95th percentile, P95); λ = 1. Because *c on v*(*A_−d_*)*⊆ c on v*(*A_full_*), this implemented budget is at least as tight as its reduced-reference variant; enlarging the budget can only widen the feasible intervals, so the both-signs saturation result is conservative with respect to that choice. Per-sample LP endpoints [*l_ik_, u_ik_*] were computed for each retained type (linear programs solved with SciPy’s `linprog`, method `highs-ds` [83], the HiGHS dual simplex [84]). Association bounds are *β_min_* = *med_i∈case,j∈control_*(*l_ik_ −u_jk_*) and *β_max_* = *med*(*u_ik_ −l_jk_*), which are sharp and exact over the samplewise product class (coordinatewise monotonicity of the median of pairwise differences; corners jointly attainable; the feasible values fill the closed interval; proof in Additional file 2: Note 2). A fixed scalar median convention is used throughout. Linear programs were solved in floating point with primal and dual feasibility tolerances of 10⁻⁸; each returned endpoint was substituted back into the original constraints and accepted only if the largest constraint violation was at most 5 × 10⁻⁸, and all 44,856 bound calculations completed without solver failure. States are classified, using a sign tolerance of 10⁻⁷, as positive sign-invariant, negative sign-invariant, sign-reversal feasible, zero-boundary or solver-failure. External-profile library analysis: for each tissue × deleted type, a prespecified library of type-matched external donor pseudobulk profiles was assembled, and each profile’s completed reference was analyzed with the single prespecified refit-on-completed strategy. Retained-type associations are computed from the full sample × type output matrix per completed reference. Envelope states require witnessed values (`library_flip_witnessed` needs an evaluated positive and an evaluated negative effect; `library_touches_zero` needs an evaluated near-zero effect), with a stated priority rule; coarse-proxy profiles (DLPFC vascular and leptomeningeal cells, VLMC) are recorded as `library_unavailable` rather than substituted. The repeated-draws cohort is excluded from main counts and analyzed only as a subject-mean supplementary set (Additional file 2: Fig. S2; Additional file 4: Data S2). Counts are reported descriptively by cohort and deletion.

### Reference operations (merge and cross-atlas transfer)

Merge: from the identical cells and donors on a fixed gene universe, Oligo and OPC were relabeled and the unique six-type terminal reference rebuilt natively. The seven-type-learned and six-type-learned operator states were each solved on the byte-identical terminal reference (hashes recorded). For MuSiC, two levels are reported: the native operational comparison (seven-type solve with outputs summed versus native six-type solve) and a locked-weight counterfactual (final seven-type gene weights applied once on the six-type reference), validated by exact replay of the six-type native output from six-type weights. The counterfactual is a mechanism diagnostic, not an official MuSiC path. Cross-atlas transfer: the donor-matched benchmark reference and the official versioned Allen Institute DFC atlas [30] (accession, version and file hash recorded) were restricted to a strict six-type common core (Astro, Micro, Oligo, OPC, Excit, Inhib; vascular classes excluded rather than approximately mapped; both label mappings prespecified before the cross-atlas comparison).

The full 2×2 of operator source × reference matrix was run for Bisque, with MuSiC native diagonals plus locked-weight off-diagonals as mechanism diagnostics. Distances are per-sample TV summarized by medians; factor-swap contrasts are reported as observed, not as additive attributions. This is a transfer experiment between two independent reference systems, not a serial version study.

### Between-path inference in real cohorts

Beyond per-path Wilcoxon/BH testing, the between-path effect difference was estimated directly per association pair as the Hodges–Lehmann location shift under each path with subject-level stratified bootstrap (10,000 resamples, fixed seed), yielding nominal 90/95% intervals and within-cohort BH q-values. Family-level inference adds per-cohort vector omnibus tests (max-|t| over the cohort’s pairs), within-cohort max-|t| simultaneous intervals, and global BH. Primary family-level inference is computed over the 952 pairs of the nine non-longitudinal cohorts, with ten-cohort tallies reported descriptively. The repeated-draw cohort was additionally re-analyzed under subject-cluster percentile bootstrap (10,000 resamples; its four negative-group clusters place percentile calibration in doubt [85, 86]), first-draw-only analysis, subject-mean aggregation, and two subject-level exact tests: permutation of the phenotype labels over all 12,650 complete assignments of four Negative labels among the 25 subjects, retaining all 98 draws and the sample-weighted statistic, and a size-stratified clustered Wilcoxon exact test in which labels are permuted within draw-count strata with stratum group counts fixed (336 assignments) [87]. All modes use BH within estimator × deletion × path. The cohort enters only as a longitudinal supplementary analysis (Additional file 2: Fig. S2; Additional file 1: Table S3).

### Coverage and certification calibration

For each derived interval S_ψ = [L, U] and prespecified oracle value ψ*: coverage = Pr{L ≤ ψ* ≤ U}; certification = Pr{0 ∉ [L, U]}; false certification = Pr{0 ∉ [L, U] and sign([L, U]) ≠ sign(ψ*)}. The shrinkage ratio is defined against the non-informative baseline ([−1, 1], width 2, for effect intervals; the no-yield interval, width 1, for absolute-scale targets). Primary rates use oracle-evaluable records as the denominator, with all-planned rates recorded alongside; missing or unavailable combinations are retained as explicit states, never encoded as zero. Three objects are kept distinct throughout: conditional sharp bounds (the identified set given the linear mixture model, the prespecified budget τ and the completion class A[n, γ], the class built from n library profiles under total-variation inflation γ, whose construction, grids and frozen metrics are given in the deposited prespecification records); the external-profile output envelope (the min–max hull of estimator outputs over actual library profiles, which is a sensitivity object rather than an identified set); and record-level binomial spread (Wilson 95% intervals [88] on aggregate rates over evaluable records, which share donors, samples and profiles and are therefore descriptive of the record set rather than of independent trials; these intervals are attached to rates and never to LP endpoints). Three oracle families are never merged: constructed truth (known generating proportions; scores point-estimator TV error only, because the estimator outputs there carry no matching interval bounds), orthogonal external measurement truth (GSE107011 flow, 29 leaves mapped to the prespecified seven types), and full-reference continuity (the effect after restoring the deleted column, or a held-out profile effect, which is a consistency target rather than independent biological truth).

The prespecified scan grid is n ∈ {2, 4, 8, 16, all} × γ ∈ {0, 0.01, 0.025, 0.05, 0.10, 0.20}, fixed programmatically before comparison with the reference values; held-out profiles never enter their own training library; the absolute-scale scan retains all 18,522 points × 7 ψ targets (129,654 records). All rates are computed directly from the released per-record tables.

### Three-ceiling decomposition

The identification ceiling is read from the conditional sharp bounds. The model ceiling is reported as item-wise residual diagnostics (full-reference relative RMSE; residual-to-signal L1 ratio; Huuki-Myers reconstruction TV tolerance; cohort marker-fit floor, held-out-gene residual and pseudobulk floor), which live on different scales and are not aggregated into one number. The algorithmic ceiling contrasts frozen NNLS and CIBERSORT on the identical 637-gene space and 29→7 leaf mapping: composition cells fixed at exactly zero by the frozen estimator (boundary zeros) cap the flow-composition coverage attainable under any positive finite yield interval; structural caps and realized coverages are reported per estimator.

The estimator contrast is diagnostic of the algorithmic layer only. No per-estimator re-inversion of the precision frontier is performed beyond the released scan.

### Shared-completion proposition and pilot

The Proposition, its joint zero-slice condition (Z), a product-form sufficient condition (Z*), the annihilation argument for shared or externally calibrated missing profiles, and the boundary list of assumption types that do delete the zero slice are stated and proved in Additional file 2: Note 3. The PBMC pilot for this Proposition evaluates two model classes at prespecified hyperparameters over 384 optimization extrema: an ambient class consistent with (Z) at its zero-exposure-floor setting, evaluated also at positive floors, and a rank-one cohort-shared class. Cells whose optimization gap exceeded the prespecified threshold are reported as unresolved computational bounds and are never counted as evidence.

### Multi-view and anchor analyses (Huuki-Myers)

All view and anchor analyses operate on the same tissue blocks under a prespecified operator class in which each block-by-view observation carries its own unknown positive diagonal RNA-yield operator, with no constraint linking these operators across views or blocks. The design has four view-count conditions plus one anchor condition: it compares feasible widths from one, two, three and six views, and from one view plus one three-type imaging panel used as a partial anchor. Equality of the multi-view widths is the constructed consequence of the operator class, a numerical instantiation of the impossibility statement rather than an independent test of it. The anchor frontier enumerates anchored-type combinations on the common roster (7 blocks, 6 donors); certification denominators aggregate donor-balanced within the 15 type-pair identities (panel-anchor instances → donor equal weight → pair-identity equal weight; 18 instances over 11 blocks and 8 donors, dual-panel blocks not collapsed). Imaging-oracle support is recorded per combination (L3: a complete complementary panel held out; L2: partial; L1: none); L3 support exists only for single-panel combinations of up to three types. Shapley attribution decomposes removable interval width between transport tolerance and counting error; it is a width attribution, not a biological variance decomposition.

The donor-permuted negative control preserves the anchor-width multiset while misassigning all donors; the prespecified contraction-validity criterion passed, and the negative-control specificity criterion reported alongside it is post hoc, defined after the permutation was run. Bootstrap CIs derive from a single prespecified seed. Where released tables differ in trailing digits because the same bootstrap stream was consumed in different orders, the canonical CI source is declared alongside the tables.

### Precision inversion (absolute scale)

The prespecified uncertainty model for absolute per-type RNA yields has three sources (member-leaf × donor heterogeneity, total-RNA-to-mRNA proxy error, and a dynamic-range penalty), each parameterized as an interval envelope with prespecified settings. The main target is the median across the 12 flow-QC-matched PBMC donors of the conditional T-cell cell-number fraction implied by the NNLS RNA contributions and the per-type yields; the width criterion is a feasible-interval width ratio ≤ 0.50 for this target relative to the no-yield width of 1. The downstream sign target ψ is the across-donor median of the T-cell cell fraction minus the T-cell RNA contribution, and the sign criterion is a ψ interval excluding zero; the seven per-type ψ targets used for coverage are defined in the same way for each mapped type. Single-source thresholds hold the other sources at their main settings; the equal joint scaling moves the proxy factor and the two log-scale envelope exponents together along one path, and the reported ±2.6% is the proxy value at which that path first fixes the sign. Donor-N requirements are modeled design thresholds under stated assumptions (donor independence, stable log-yield variance, no additional batch structure, centers fixed at their prespecified values), not extrapolations from the observed donors. fitdrop 0.2.0 implements the inversion as a callback engine: the caller supplies an `eval_bounds` function, uncertainty sources parameterized as interval envelopes via `precision_parameter_grid(interval_envelope = TRUE)`, and either verified monotonicity or grid-only evaluation. Non-compliant inputs stop with an explicit error rather than being silently approximated. Estimator boundary zeros are reported as an algorithmic-ceiling flag and excluded from the inversion. The package passes R CMD check [89]. The released tests reproduce the PBMC and Huuki-Myers vignette headline numbers.

### Null recalibration

Structural-layer significance was re-evaluated by replacing the uniform receiver null with the realized receiver-margin null via exact enumeration over 104 units, with a prespecified effect-size criterion; empirical type-I calibration used 1,000 no-effect replicates for each of the 104 realized-margin units.

## Data and code availability

All datasets are public, and dataset and reference accessions are listed in Additional file 1: Table S1. Analysis code, the fitdrop R package (0.2.0, including `precision_frontier()` and the PBMC and Huuki-Myers vignettes), the dated decision record with prespecified criteria, the coverage-calibration, anchor-frontier and precision-inversion per-record tables, and per-sample outputs are deposited on Zenodo [90]; the record is listed under Availability of data and materials.

## Use of large language models

During the preparation of this work, the authors used Claude (Anthropic) and ChatGPT (OpenAI) for methodological consistency checks and to improve the readability and language of the final manuscript. The authors reviewed and edited all output and take full responsibility for the content of the published work.

## Declarations

## Ethics approval and consent to participate

Not applicable. This study analyzed only publicly available, de-identified data from the repositories listed in Additional file 1: Table S1; no new data involving human participants or animals were generated.

## Consent for publication

Not applicable.

## Availability of data and materials

All datasets analyzed in this study are publicly available; every accession, version and landing URL is recorded in Additional file 1: Table S1. Analysis code, the fitdrop R package (version 0.2.0, including `precision_frontier()` and the PBMC and Huuki-Myers vignettes), the dated decision record with prespecified criteria, the coverage-calibration, anchor-frontier and precision-inversion per-record tables, and per-sample outputs are deposited on Zenodo (https://doi.org/10.5281/zenodo.22300711).

## Authors’ contributions

HJ conceived and designed the study, performed the analyses and wrote the manuscript. FG, PL and YW contributed to the study design, interpreted the results and revised the manuscript. Ying Jie, YL and Yang Jiang supervised the study and revised the manuscript. All authors read and approved the final manuscript.

## Acknowledgements

Not applicable.

## Additional files

- **Additional file 1.** File name: `Additional_file_1_Supplementary_Tables.xlsx`; Format: XLSX; Title: Supplementary Tables; Description: Tables S1–S3, containing dataset and reference accessions, cohort screening and re-screening decisions, and GSE166190 longitudinal sensitivity results.
- **Additional file 2.** File name: `Additional_file_2_Supplementary_Notes_and_Figures.pdf`; Format: PDF; Title: Supplementary Notes and Figures; Description: Supplementary Notes 1–3 and Figs. S1–S4 with their captions. Note 3 states and proves the shared-completion Proposition, its joint zero-slice condition (Z), the product-form sufficient condition (Z*), and the boundary list of assumption classes that do delete the zero slice.
- **Additional file 3.** File name: `Additional_file_3_Data_S1.zip`; Format: ZIP; Title: Supplementary Data S1: exact theorem verification; Description: The exact-verification script and its full 80-check output.
- **Additional file 4.** File name: `Additional_file_4_Data_S2.zip`; Format: ZIP; Title: Supplementary Data S2: completion-library records; Description: Profile-fit, profile-effect, envelope and failure records for the completion-library analysis, with analysis scope retained.

