## Additional file 2: Supplementary Notes and Figures for "Incomplete references leave bulk deconvolution targets non-identifiable, but identification has a computable precision price"

### Contents


| Section | Page |
| --- | --- |
| Supplementary Note 1. Complete construction for non-identification under admissible completion | 3-9 |
| Supplementary Note 2. Sharpness of the Hodges–Lehmann feasible-effect interval | 10-11 |
| Supplementary Note 3. The shared-completion proposition: replication without compelled exposure is not identification | 12-15 |
| Fig. S1 | 16 |
| Fig. S1 caption | 17 |
| Fig. S2 | 18 |
| Fig. S2 caption | 19 |
| Fig. S3 | 20 |
| Fig. S3 caption | 21 |
| Fig. S4 | 22 |
| Fig. S4 caption | 23 |

### Supplementary Note 1. Complete construction for non-identification under admissible completion

#### N1.1 Observed object, completion class and estimand

For an integer $m \geq 1$, let

$$\Delta_{m-1} = \{v \in \mathbb{R}_+^m : \mathbf{1}^\top v = 1\}, \qquad \Delta_{m-1}^\circ = \{v \in \Delta_{m-1} : v_j > 0 \text{ for all } j\}. \tag{N1.1}$$

The observed input is $O = (y, A_{-d})$, where $y \in \Delta_{P-1}$ is the observed bulk composition and $A_{-d} \in \mathbb{R}_+^{P\times K}$ is the incomplete reference, with every column of $A_{-d}$ summing to one. A complete world is a pair $W = (a, x)$, where $a \in \Delta_{P-1}$ is the missing reference column and $x \in \Delta_K^\circ$ is the composition over the $K$ retained types and the missing type. The world must reconstruct the observation exactly:

$$y = [A_{-d}, a]x. \tag{N1.2}$$

The admissible completion class is therefore

$$\mathcal{C}(O) = \{(a, x) : a \geq 0,\ \mathbf{1}^\top a = 1,\ x \in \Delta_K^\circ,\ y = [A_{-d}, a]x\}. \tag{N1.3}$$

The world contains both the completion column and its generating composition. This distinction prevents a solver-dependent coefficient vector from being substituted for the estimand when a completed system is underdetermined. The estimand is the retained-type conditional composition

$$T(W) = \frac{x_{1:K}}{\sum_{j=1}^K x_j} \in \Delta_{K-1}^\circ, \tag{N1.4}$$

and distance between compositions is total variation,

$$d_{\mathrm{TV}}(u, v) = \frac{1}{2}\|u - v\|_1. \tag{N1.5}$$

Throughout this note, the claims concern the unrestricted admissible completion class $\mathcal{C}(O)$ in (N1.3); anchor references, cross-sample structure, restrictions on the missing-column shape, or external truth define narrower classes.

#### N1.2 Theorems 1 and 2

**Theorem 1 (maximal non-identification under admissible completion).** The following statements hold at three distinct quantifier levels.

1. There exists a strictly positive observation $O = (y, A_{-d})$ with $P = 3$ and $K = 2$ for which

$$\{T(W) : W \in \mathcal{C}(O)\} = \Delta_1^\circ. \tag{N1.6}$$

   For every target $(t, 1-t)$ with $0 < t < 1$, a member of the explicit realizing family in Proposition N1.1 attains that target. Each completed reference in this explicit family is nonsingular, its coefficient vector is unique, and its 1-norm condition number is at most $1313/485$. At the same observation, opposite retained-type rankings persist on two relative-open neighborhoods of strictly positive completion columns whose completed references remain nonsingular and uniformly well conditioned.

2. For every $P \geq 3$ and $K \geq 2$, there exists a strictly positive two-world embedding with a common observed input and opposite rankings of the first two retained types.

3. In the full-column-rank feasible regime $P \geq K + 1$, a separate strictly positive two-world embedding can be chosen so that both completed references have column rank $K + 1$.

**Theorem 2 (full-class decision consequences at the Theorem 1 observation).** At the observation in the first part of Theorem 1, fix

$$q_F = (3/5, 2/5), \qquad q_R = (3/10, 7/10), \qquad \delta = d_{\mathrm{TV}}(q_F, q_R) = 3/10, \tag{N1.7}$$

and define $R_H(W) = d_{\mathrm{TV}}(q_H, T(W))$ for $H \in \{F, R\}$. For binary actions, oracle-relative regret is

$$R_H(W) - \min\{R_F(W), R_R(W)\}. \tag{N1.8}$$

The deterministic binary oracle-relative minimax regret over the full class $\mathcal{C}(O)$ is

$$\inf_{g} \sup_{W\in\mathcal{C}(O)} \left[R_{g(O)}(W) - \min\{R_F(W), R_R(W)\}\right] = 3/10. \tag{N1.9}$$

The expected oracle-relative minimax regret for a randomized binary selector is

$$\inf_{p\in[0,1]} \sup_{W\in\mathcal{C}(O)} \mathrm{Reg}_p(W) = 3/20. \tag{N1.10}$$

For an observed-only composition-valued output $q \in \Delta_1$, the oracle-relative minimax regret is

$$\inf_{q\in\Delta_1} \sup_{W\in\mathcal{C}(O)} [d_{\mathrm{TV}}(q, T(W)) - \min\{R_F(W), R_R(W)\}] = 3/20, \tag{N1.11}$$

whereas its composition-valued minimax raw risk is

$$\inf_{q\in\Delta_1} \sup_{W\in\mathcal{C}(O)} d_{\mathrm{TV}}(q, T(W)) = 1/2. \tag{N1.12}$$

##### N1.3 A strictly positive two-world witness

Define the column-stochastic smoothing matrix

$$S = \frac{97}{100} I_3 + \frac{1}{100}\mathbf{1}\mathbf{1}^\top = \begin{pmatrix} 49/50 & 1/100 & 1/100 \\ 1/100 & 49/50 & 1/100 \\ 1/100 & 1/100 & 49/50 \end{pmatrix}. \tag{N1.13}$$

For every three-dimensional composition $v$, $Sv = (97/100)v + (1/100)\mathbf{1}$. Thus $S$ preserves column sums and maps every non-negative composition into the strict interior. Its eigenvalues are $1, 97/100, 97/100$, so it is invertible. Set $b_1 = Se_1$, $b_2 = Se_2$, and

$$A_{-d} = S[e_1, e_2] = \begin{pmatrix} 49/50 & 1/100 \\ 1/100 & 49/50 \\ 1/100 & 1/100 \end{pmatrix}, \qquad y = S(3/10, 1/5, 1/2)^\top = (301/1000, 51/250, 99/200)^\top. \tag{N1.14}$$

The two worlds are

$$\begin{aligned} W_1 :\quad & a_1^\star = Se_3 = (1/100, 1/100, 49/50)^\top, \quad x^{(1)} = (3/10, 1/5, 1/2)^\top, \\ W_2 :\quad & a_2^\star = S(3/10, 1/10, 3/5)^\top = (301/1000, 107/1000, 74/125)^\top, \quad x^{(2)} = (1/20, 7/60, 5/6)^\top. \end{aligned} \tag{N1.15}$$

The unsmoothed standard-basis construction is the preimage of this witness under the invertible map $S$; the formal witness used here is its strictly positive smoothed image.

**Lemma N1.1 (strictly positive observational equivalence and ranking reversal).** Both $W_1$ and $W_2$ belong to $\mathcal{C}(O)$, and their retained-type conditional compositions have opposite strict rankings.

**Proof.** Every entry of $A_{-d}$, $y$, $a_i^\star$, and $x^{(i)}$ is strictly positive. Every reference column, each completion column, and $y$ sums to one, and each $x^{(i)}$ lies in $\Delta_2^\circ$. In the unsmoothed coordinates,

$$[e_1, e_2, e_3]x^{(1)} = (3/10, 1/5, 1/2)^\top, \qquad [e_1, e_2, (3/10, 1/10, 3/5)^\top]x^{(2)} = (3/10, 1/5, 1/2)^\top. \tag{N1.16}$$

Multiplication by $S$ yields $[A_{-d}, a_i^\star]x^{(i)} = y$ for both worlds. Conditionalizing the first two coordinates gives

$$T(W_1) = (3/5, 2/5), \qquad T(W_2) = (3/10, 7/10). \tag{N1.17}$$

Hence $T_1(W_1) > T_2(W_1)$ and $T_1(W_2) < T_2(W_2)$, while

$$d_{\mathrm{TV}}(T(W_1), T(W_2)) = 3/10. \tag{N1.18}$$

These identities establish Lemma N1.1.

##### N1.4 The witness is well conditioned

**Lemma N1.2 (conditioning at the witness centers).** The observed incomplete reference and both completed references in Lemma N1.1 are well conditioned.

**Proof.** The squared 2-norm condition number of the observed reference is

$$\kappa_2(A_{-d})^2 = \frac{9803}{9409}, \qquad \kappa_2(A_{-d}) = \sqrt{\frac{9803}{9409}} = 1.020723\ldots. \tag{N1.19}$$

The exact 1-norm condition numbers of the two square completed references are

$$\kappa_1([A_{-d}, a_1^\star]) = \frac{101}{97} = 1.041237\ldots, \qquad \kappa_1([A_{-d}, a_2^\star]) = \frac{233}{97} = 2.402062\ldots. \tag{N1.20}$$

Thus the ranking reversal occurs at strictly positive, nonsingular, well-conditioned completed references.

##### N1.5 Ranking reversal persists on relative-open neighborhoods

Let $a_i^\star$ be the two center columns in (N1.15). In the affine plane of column-stochastic completion vectors, define

$$U_i = \{a_i^\star + h(u,v) : h(u,v) = (u, v, -u-v)^\top,\ \|h(u,v)\|_\infty < \rho\}, \qquad \rho = \frac{1}{200}. \tag{N1.21}$$

All physical completion columns in $U_1$ and $U_2$ are strictly positive. Because $\mathbf{1}^\top h = 0$,

$$S^{-1}h = \frac{100}{97}h, \qquad e = \frac{100}{97}\rho = \frac{1}{194}, \qquad \widetilde{a}_i = a_i^{(0)} + S^{-1}h, \tag{N1.22}$$

where $a_1^{(0)} = e_3$ and $a_2^{(0)} = (3/10, 1/10, 3/5)^\top$. The exact coefficient vector for a completion $a = a_i^\star + h$ is

$$x_3 = \frac{1/2}{\widetilde{a}_{i3}}, \qquad x_1 = \frac{3}{10} - \widetilde{a}_{i1}x_3, \qquad x_2 = \frac{1}{5} - \widetilde{a}_{i2}x_3. \tag{N1.23}$$

**Lemma N1.3 (relative-open persistence).** Every $a \in U_i$ has a unique admissible coefficient vector $x(a)$ given by (N1.23). The first retained type ranks above the second throughout $U_1$, the second ranks above the first throughout $U_2$, and the completed references are uniformly well conditioned.

**Proof.** The perturbation bound in (N1.22) keeps $\widetilde{a}_{i3}$ strictly positive, so $[e_1, e_2, \widetilde{a}_i]$ is nonsingular and (N1.23) is its unique coefficient vector. Direct worst-direction bounds over $\|h\|_\infty < \rho$ give, for every $a \in U_1$,

$$x_1 > \frac{287}{965}, \qquad x_2 > \frac{381}{1930}, \qquad x_1 - x_2 > \frac{183}{1930} > 0, \tag{N1.24}$$

and, for every $a \in U_2$,

$$x_1 > \frac{251}{5770}, \qquad x_2 > \frac{322}{2885}, \qquad x_2 - x_1 > \frac{333}{5870} > 0. \tag{N1.25}$$

Equation (N1.23) also gives $x_3 > 0$. Exact reconstruction and the unit column sums imply $\sum_j x_j = 1$, so $x(a)$ lies in $\Delta_2^\circ$. Because conditionalization divides $x_1 - x_2$ by a positive denominator, the signs in (N1.24) and (N1.25) establish the stated ranking directions.

For uniform conditioning, write $[A_{-d}, a] = S[e_1, e_2, \widetilde{a}]$. On the closures of the two neighborhoods,

$$\sup_{\overline{U}_1} \|[e_1, e_2, \widetilde{a}]^{-1}\|_1 = \frac{195}{193}, \qquad \sup_{\overline{U}_2} \|[e_1, e_2, \widetilde{a}]^{-1}\|_1 = \frac{1363}{577}. \tag{N1.26}$$

The physical completed matrices are column stochastic, so their 1-norm is one, and $\|S^{-1}\|_1 = 101/97$. Consequently,

$$\sup_{a\in\overline{U}_1} \kappa_1([A_{-d}, a]) \le \frac{19695}{18721} < \frac{19796}{18721}, \qquad \sup_{a\in\overline{U}_2} \kappa_1([A_{-d}, a]) \le \frac{137663}{55969} < \frac{138168}{55969}. \tag{N1.27}$$

These bounds establish Lemma N1.3. As a finite exact-arithmetic check, the accompanying script evaluates 1,141 rational lattice points in each neighborhood and preserves admissibility and the corresponding strict ranking at every point.

##### N1.6 The identified set is the full open simplex

**Proposition N1.1 (maximal identified set at the base observation).** For the fixed $P = 3, K = 2$ observation in (N1.14), the identified set is exactly $\Delta_1^\circ$. Every target in this set is attained by a nonsingular member of an explicit realizing family whose 1-norm condition number is at most $1313/485$.

**Proof.** For any $t \in (0,1)$, define

$$x(t) = \left(\frac{t}{10}, \frac{1-t}{10}, \frac{9}{10}\right), \qquad \widetilde{a}(t) = \left(\frac{3-t}{9}, \frac{1+t}{9}, \frac{5}{9}\right), \qquad a(t) = S\widetilde{a}(t). \tag{N1.28}$$

These vectors are strictly positive and have the required unit sums. Moreover,

$$[e_1, e_2, \widetilde{a}(t)]x(t) = (3/10, 1/5, 1/2)^\top, \qquad [A_{-d}, a(t)]x(t) = y, \qquad T(W(t)) = (t, 1-t). \tag{N1.29}$$

The third diagonal entry of $[e_1, e_2, \widetilde{a}(t)]$ is $5/9$, so the completed matrix is nonsingular and $x(t)$ is its unique coefficient vector. For this explicit realizing family,

$$\|[e_1, e_2, \widetilde{a}(t)]^{-1}\|_1 = \frac{13}{5}, \qquad \kappa_1([A_{-d}, a(t)]) \le \frac{13}{5}\frac{101}{97} = \frac{1313}{485} \quad \text{for every } t \in (0,1). \tag{N1.30}$$

Equation (N1.29) shows that every $(t, 1-t) \in \Delta_1^\circ$ is attained by a member of this explicit family. Conversely, every admissible $x \in \Delta_2^\circ$ has $x_1, x_2 > 0$, so its retained conditional composition belongs to $\Delta_1^\circ$. The two inclusions yield

$$\{T(W) : W \in \mathcal{C}(O)\} = \Delta_1^\circ. \tag{N1.31}$$

##### N1.7 Dimension embeddings and rank regimes

#### N1.7a Strictly positive two-world embeddings for every $P \ge 3, K \ge 2$

**Proposition N1.2 (dimension embedding of two-world ranking reversal).** For every $P \ge 3$ and $K \ge 2$, the two worlds in Lemma N1.1 admit a strictly positive embedding with a common observation and opposite rankings of the first two retained types.

**Proof.** Define the column-stochastic lifting matrix $H_P \in \mathbb{R}^{P\times 3}$ by

$$(H_P)_{rj} = \frac{1}{2P} + \frac{1}{2}\mathbf{1}\{r = j\}, \qquad r = 1, \dots, P, \quad j = 1, 2, 3. \tag{N1.32}$$

Its leading $3 \times 3$ minor is $\frac{1}{2}I_3 + \frac{1}{2P}J_3$, with

$$\det\left(\tfrac{1}{2}I_3 + \tfrac{1}{2P}J_3\right) = \frac{P+3}{8P} > 0. \tag{N1.33}$$

Hence $\operatorname{rank}(H_P) = 3$. Let

$$\eta = \frac{1}{4K}, \qquad \tau = (K-2)\eta, \qquad \lambda = 1 - \tau = \frac{3K+2}{4K} > 0. \tag{N1.34}$$

Using $b_1 = Se_1$ and $b_2 = Se_2$, define the shared observation and the two completion columns by

$$y^{(P)} = H_P y, \qquad A_{-d}^{(P,K)} = \left[ H_P b_1, H_P b_2, \underbrace{H_P y, \dots, H_P y}_{K-2 \text{ columns}} \right], \qquad a_i^{(P)} = H_P a_i^\star. \tag{N1.35}$$

Embed the two coefficient vectors as

$$x_i^{(P,K)} = \left( \lambda x_{i1}, \lambda x_{i2}, \underbrace{\eta, \dots, \eta}_{K-2 \text{ retained coordinates}}, \lambda x_{i3} \right). \tag{N1.36}$$

All physical columns, the bulk, and the weights are strictly positive and sum to one. For both worlds,

$$[A_{-d}^{(P,K)}, a_i^{(P)}]x_i^{(P,K)} = \lambda H_P y + (K-2)\eta H_P y = H_P y. \tag{N1.37}$$

After conditioning on the retained coordinates, the difference between the first two coordinates has a positive denominator and numerator $\lambda(x_{i1} - x_{i2})$. The opposite strict rankings from Lemma N1.1 therefore persist for every stated $P$ and $K$. This construction asserts two-world non-identification and ranking reversal; it does not assert a full-simplex identified set for general $K$.

##### N1.7b Full-column-rank embeddings when $P \geq K+1$

**Proposition N1.3 (full-column-rank feasible regime).** For every $K \geq 2$ and $P \geq K+1$, a separate strictly positive embedding has two observationally equivalent worlds with opposite first-two-type rankings and completed-reference rank $K+1$.

**Proof.** Define

$$G_P = \frac{1}{2} I_P + \frac{1}{2P} \mathbf{1}\mathbf{1}^\top. \tag{N1.38}$$

Before smoothing, take the retained reference columns and the two completion columns to be

$$\overline{A}_{-d}^{(P,K)} = [e_1, e_2, e_4, \ldots, e_{K+1}], \qquad c_1 = e_3, \qquad c_2 = (3/10, 1/10, 3/5, 0, \ldots, 0)^\top. \tag{N1.39}$$

With $\eta$ and $\lambda$ from (N1.34), define the shared raw bulk by

$$\overline{y}^{(P,K)} = \lambda(3/10, 1/5, 1/2, 0, \ldots, 0)^\top + \eta \sum_{j=4}^{K+1} e_j. \tag{N1.40}$$

The two raw worlds reconstruct $\overline{y}^{(P,K)}$ with the coefficient vectors in (N1.36). Apply $G_P$ to every reference column, completion column, and the bulk:

$$A_{-d}^{(P,K)} = G_P \overline{A}_{-d}^{(P,K)}, \qquad a_i^{(P)} = G_P c_i, \qquad y^{(P,K)} = G_P \overline{y}^{(P,K)}. \tag{N1.41}$$

The smoothed reference columns and bulk are strictly positive and sum to one, the weights lie in the simplex interior, and exact reconstruction and the opposite rankings are preserved. Let $\mathcal{R} = (1, 2, 4, \ldots, K+1, 3)$ denote the row order selecting the square minors. Their absolute determinants are

$$\left|\det\left([\overline{A}_{-d}^{(P,K)}, c_1]_{\mathcal{R}}\right)\right| = 1, \qquad \left|\det\left([\overline{A}_{-d}^{(P,K)}, c_2]_{\mathcal{R}}\right)\right| = 3/5. \tag{N1.42}$$

Both raw completed references therefore have rank $K+1$. Since $G_P$ is invertible, smoothing preserves this rank.

##### N1.8 Full-class minimax consequences

At the base observation, let $q_F$, $q_R$, and $\delta$ be as in (N1.7). Lemma N1.1 supplies two worlds satisfying

$$T(W_1) = q_F, \qquad T(W_2) = q_R, \qquad (R_F(W_1), R_R(W_1)) = (0, \delta), \qquad (R_F(W_2), R_R(W_2)) = (\delta, 0). \tag{N1.43}$$

#### Deterministic binary oracle-relative regret

**Proposition N1.4 (deterministic binary oracle-relative minimax regret).** The value in (N1.9) is exactly $\delta = 3/10$.

**Proof.** An observed-only selector must take the same action in $W_1$ and $W_2$ because both worlds generate the same $O$. Equation (N1.43) therefore gives the lower bound $\delta$. For any $W \in \mathcal{C}(O)$ and either fixed action $H \in \{F, R\}$,

$$0 \leq R_H(W) - \min\{R_F(W), R_R(W)\} \leq |R_F(W) - R_R(W)| \leq d_{\mathrm{TV}}(q_F, q_R) = \delta. \tag{N1.44}$$

The final inequality is the reverse triangle inequality. It gives a full-class upper bound for either action, and hence

$$\inf_g \sup_{W \in \mathcal{C}(O)} \left[R_{g(O)}(W) - \min\{R_F(W), R_R(W)\}\right] = \delta = 3/10. \tag{N1.45}$$

#### Randomized binary expected oracle-relative regret

**Proposition N1.5 (randomized binary expected oracle-relative minimax regret).** The value in (N1.10) is exactly $\delta/2 = 3/20$.

**Proof.** Let an observed-only rule choose $F$ with probability $p \in [0, 1]$. Its expected oracle-relative regrets in $W_1$ and $W_2$ are, respectively, $(1-p)\delta$ and $p\delta$. Thus

$$\sup_{W \in \mathcal{C}(O)} \mathrm{Reg}_p(W) \geq \delta \max\{p, 1-p\} \geq \delta/2. \tag{N1.46}$$

For $p = 1/2$ and every $W \in \mathcal{C}(O)$,

$$\mathrm{Reg}_{1/2}(W) = \frac{1}{2}|R_F(W) - R_R(W)| \leq \delta/2. \tag{N1.47}$$

Therefore

$$\inf_{p \in [0,1]} \sup_{W \in \mathcal{C}(O)} \mathrm{Reg}_p(W) = \delta/2 = 3/20. \tag{N1.48}$$

#### Composition-valued oracle-relative regret and composition-valued raw risk

**Proposition N1.6 (composition-valued oracle-relative minimax regret and raw minimax risk).** Over observed-only outputs $q \in \Delta_1$, the oracle-relative value in (N1.11) is $\delta/2 = 3/20$, while the raw-risk value in (N1.12) is $1/2$.

**Proof.** Define composition-valued oracle-relative regret by

$$\mathrm{Reg}(q, W) = d_{\mathrm{TV}}(q, T(W)) - \min\{R_F(W), R_R(W)\}. \tag{N1.49}$$

The oracle benchmark is zero in the corresponding witness world. The triangle inequality therefore gives

$$\max_{i=1,2} \mathrm{Reg}(q, W_i) = \max\{d_{\mathrm{TV}}(q, q_F), d_{\mathrm{TV}}(q, q_R)\} \geq \delta/2. \tag{N1.50}$$

The midpoint is

$$m = \frac{q_F + q_R}{2} = (9/20, 11/20). \tag{N1.51}$$

For every $W \in \mathcal{C}(O)$,

$$d_{\mathrm{TV}}(m, T(W)) \leq R_F(W) + \delta/2, \qquad d_{\mathrm{TV}}(m, T(W)) \leq R_R(W) + \delta/2. \tag{N1.52}$$

Subtracting the smaller benchmark gives $\mathrm{Reg}(m, W) \leq \delta/2$. Combined with (N1.50), this yields

$$\inf_{q \in \Delta_1} \sup_{W \in \mathcal{C}(O)} \mathrm{Reg}(q, W) = \delta/2 = 3/20. \tag{N1.53}$$

For raw risk, Proposition N1.1 identifies every $T(W)$ with $(t, 1-t)$ for $t \in (0,1)$. Writing $q = (r, 1-r)$ gives $d_{\mathrm{TV}}(q, T(W)) = |r - t|$, so the supremum is $\max\{r, 1-r\}$ and is minimized at $r = 1/2$. Hence

$$\inf_{q \in \Delta_1} \sup_{W \in \mathcal{C}(O)} d_{\mathrm{TV}}(q, T(W)) = 1/2. \tag{N1.54}$$

#### N1.9 Analytic scope and exact verification

**Exact verification.** The accompanying exact-arithmetic script verifies the displayed witnesses and rational constants, 2,282 finite lattice points, five representative values of $t$, and 54 representative dimensional embeddings (80 checks). Universality over the relative-open neighborhoods, all $t \in (0,1)$, the stated dimension regimes, and the full admissible completion class is established by the analytic inequalities and constructions given above.

The script and its matching output are provided in Data S1. Running `python3 verify_theorem_20260808.py` produces the terminal line `ALL CHECKS PASS | checks=80`.

##### N1.10 Relation to operator provenance

Theorem 1 does not require an adaptive operator: completion non-identification persists when the operator is fixed. Conversely, operator provenance does not follow from Theorem 1; $q_F$ and $q_R$ are fixed candidate outputs produced by two histories at the same observed input. Theorem 2 connects the two layers only for this fixed observation, this candidate pair, and the full admissible completion class: changing the unobserved completion changes which candidate is closer to the retained-type conditional estimand.

##### N1.11 Relation to yield calibration

The absolute-scale system of the main text converts RNA contributions into cell fractions through per-type RNA yields. Write $d \in (0, \infty)^{K+1}$ for the yields (RNA mass per cell) of the $K$ retained types and the missing type. A completed composition $x$ of RNA contributions corresponds to the cell fractions

$$c_k(x, d) = \frac{x_k/d_k}{\sum_{j=1}^{K+1} x_j/d_j}, \qquad k = 1, \ldots, K+1. \tag{N1.55}$$

Exact yields identify this conversion: for fixed positive $d$, $x \mapsto c(x, d)$ is a bijection of $\Delta_K^\circ$ onto itself. The yields enter only after a completed world has been chosen, so they leave the completion step of (N1.3) untouched.

**Corollary N1.1 (exact yields leave the Theorem 1 ambiguity intact).** Assign one exact yield vector $d$ to both witness worlds of (N1.15). Then $W_1$ and $W_2$ still reproduce the observation $y$ of (N1.14), and their retained-type conditional cell fractions differ for every positive $d$. With $d = \mathbf{1}$ the cell fractions coincide with the RNA contributions, and the retained targets are $(3/5, 2/5)$ under $W_1$ and $(3/10, 7/10)$ under $W_2$.

**Proof.** The data-fit constraint (N1.2) involves only the completed reference and the RNA contributions, so it holds for both worlds whatever $d$ is chosen (Lemma N1.1). By (N1.55), the retained-type conditional cell fraction of the first type is $c_1/(c_1 + c_2) = (x_1/d_1)/(x_1/d_1 + x_2/d_2)$. The two worlds give the same value only if $x_1^{(1)} x_2^{(2)} = x_2^{(1)} x_1^{(2)}$, that is $(3/10)(7/60) = (1/5)(1/20)$, or $7/200 = 1/100$, which is false. For $d = \mathbf{1}$ the conditional fractions reduce to $x_1/(x_1 + x_2)$, namely $3/5$ and $3/10$. □

Yield calibration therefore determines the conversion from RNA contribution to cell number, whose precision requirements the main-text frontier computes; selecting among admissible completions requires additional information.

### Supplementary Note 2. Sharpness of the Hodges–Lehmann feasible-effect interval

#### N2.1 Setup and proposition

For each sample $i$, the sample-wise feasible-set analysis (Methods) defines the feasibility class

$$X_i = \{x \in \Delta :\ d_{\mathrm{TV}}(y_i, A_{-d}x) \le \tau_i\}. \tag{N2.1}$$

Each $X_i$ is assumed to be nonempty, closed, and convex and to lie in the closed simplex $\Delta$. Because the closed simplex is compact, every $X_i$ is compact. For a fixed retained type $k$, define the coordinate endpoints

$$l_{ik} = \min_{x \in X_i} x_k, \qquad u_{ik} = \max_{x \in X_i} x_k. \tag{N2.2}$$

For the finite collection of observed samples, with nonempty, disjoint group index sets $I_{\mathrm{case}}$ and $I_{\mathrm{control}}$, the samplewise product class and its Hodges–Lehmann location shift are

$$\mathcal{X} = \prod_i X_i, \qquad \beta(x) = \mathrm{med}_{i \in I_{\mathrm{case}},\, j \in I_{\mathrm{control}}}(x_{ik} - x_{jk}). \tag{N2.3}$$

We use the fixed scalar median convention of the main text throughout: for an odd number of pairwise differences, the median is the middle order statistic; for an even number, it is the arithmetic mean of the two middle order statistics.

**Proposition N2.1 (sharp Hodges–Lehmann interval).** The image of the samplewise product class under the Hodges–Lehmann location shift is

$$\beta(\mathcal{X}) = [\beta_{\min}, \beta_{\max}], \tag{N2.4}$$

where

$$\beta_{\min} = \mathrm{med}_{i \in I_{\mathrm{case}},\, j \in I_{\mathrm{control}}}(l_{ik} - u_{jk}), \qquad \beta_{\max} = \mathrm{med}_{i \in I_{\mathrm{case}},\, j \in I_{\mathrm{control}}}(u_{ik} - l_{jk}). \tag{N2.5}$$

#### N2.2 Coordinate ranges of the samplewise classes

The coordinate map $x \mapsto x_k$ is linear and continuous. The image of the nonempty convex set $X_i$ under this map is therefore a nonempty convex subset of the real line, and the image of the compact set $X_i$ is compact. Hence

$$\{x_k : x \in X_i\} = [l_{ik}, u_{ik}]. \tag{N2.6}$$

In particular, both endpoints in (N2.2) are attained by elements of $X_i$, and every value between them is also attained. The total-variation constraint in (N2.1) has a linear-programming representation, so these attained coordinate extrema are exactly the per-sample LP endpoints used in the sample-wise feasible-set analysis.

##### N2.3 Coordinatewise monotonicity and extremal corners

Increasing a case coordinate $x_{ik}$ weakly increases every pairwise difference involving that case sample and leaves all other pairwise differences unchanged. Increasing a control coordinate $x_{jk}$ weakly decreases every pairwise difference involving that control sample and leaves all other pairwise differences unchanged. Every scalar order statistic is coordinatewise nondecreasing in its inputs, and the even-sample median is the mean of two such order statistics. Consequently, $\beta(x)$ is nondecreasing in each case coordinate and nonincreasing in each control coordinate. Thus, for every $x \in \mathcal{X}$,

$$\beta_{\min} \leq \beta(x) \leq \beta_{\max}. \tag{N2.7}$$

The lower and upper bounds are jointly attainable. For the lower corner, choose for every case sample $i \in I_{\text{case}}$ an element of $X_i$ attaining $x_{ik} = l_{ik}$, and choose for every control sample $j \in I_{\text{control}}$ an element of $X_j$ attaining $x_{jk} = u_{jk}$. Because $\mathcal{X} = \prod_i X_i$, these per-sample choices are independent and assemble into one element of $\mathcal{X}$. At that element, all pairwise differences equal $l_{ik} - u_{jk}$, so $\beta(x) = \beta_{\min}$. For the upper corner, independently choose $x_{ik} = u_{ik}$ for every case sample and $x_{jk} = l_{jk}$ for every control sample. These choices likewise assemble into one element of $\mathcal{X}$, at which all pairwise differences equal $u_{ik} - l_{jk}$ and $\beta(x) = \beta_{\max}$. This product-class construction establishes the global minimum and maximum, rather than only separate attainability of the coordinate endpoints.

##### N2.4 Sharpness of the full interval

Each pairwise difference $x_{ik} - x_{jk}$ is affine in $x$. Every order statistic of finitely many real-valued inputs is continuous, and the fixed scalar median is either one order statistic or the mean of two adjacent order statistics. Therefore $\beta$ is continuous on $\mathcal{X}$.

The finite product $\mathcal{X}$ is compact and convex because each $X_i$ is compact and convex; in particular, $\mathcal{X}$ is connected. Its continuous image $\beta(\mathcal{X})$ is therefore a compact, connected subset of the real line and hence a closed interval. Section N2.3 shows that the interval has minimum $\beta_{\min}$ and maximum $\beta_{\max}$. It follows that every value between these endpoints is attained and that (N2.4) is the complete sharp identified range.

##### N2.5 Computational realization and scope

The per-sample linear programs yield $l_{ik}$ and $u_{ik}$, and Proposition N2.1 converts those coordinate endpoints into the effect-level sharp interval in (N2.4)–(N2.5). As reported in the Methods, the LPs are solved in floating-point arithmetic with primal and dual feasibility tolerances of $10^{-8}$, and each returned endpoint is substituted back into the original constraints and accepted only if the largest constraint violation is at most $5 \times 10^{-8}$.

The sharp endpoint formulas apply to the samplewise product class $\mathcal{X} = \prod_i X_i$ used in the sample-wise feasible-set analysis. Completion classes that impose cross-sample coupling need not admit the two joint extremal corners and require a separate bound.

#### Supplementary Note 3. The shared-completion proposition: replication without compelled exposure is not identification

##### N3.1 Setting: a joint model with a cohort-shared completion profile

Let

$$\mathcal{X} = \prod_{i=1}^{n} \mathcal{X}_i, \qquad \mathcal{H} \subseteq \mathbb{R}_+^n, \qquad \mathcal{A} \neq \varnothing \tag{N3.1}$$

denote, respectively, the joint retained-coordinate space over the $n$ samples, the admissible class of hidden-exposure vectors $h = (h_1, \ldots, h_n)$, and the admissible class of cohort-shared completion profiles $a$. All data-fit requirements, normalization, non-negativity, and any within- or cross-sample constraints are collected into a single feasibility set

$$\mathcal{F}_{\mathrm{sh}} \subseteq \mathcal{X} \times \mathcal{H} \times \mathcal{A}. \tag{N3.2}$$

In the additive residual model this set takes the form

$$\mathcal{F}_{\mathrm{sh}} = \{(x, h, a) \in \mathcal{C} : \rho_i(y_i,\ A_{-d} x_i + h_i a) \leq \tau_i,\ i = 1, \ldots, n\}, \tag{N3.3}$$

where $\mathcal{C}$ collects all non-data-fit constraints and $\rho_i$ is a general acceptance function, so that the statement covers exact fit, norm residual budgets, and any other deterministic feasibility criterion alike.

The zero-exposure submodel has per-sample feasible sets

$$\mathcal{X}_i^{(0)} = \{x_i : \rho_i(y_i, A_{-d} x_i) \leq \tau_i \text{ and the within-sample constraints of the zero-exposure model hold}\}, \tag{N3.4}$$

with joint class

$$\varnothing \neq \mathcal{X}^{(0)} = \prod_{i=1}^{n} \mathcal{X}_i^{(0)}. \tag{N3.5}$$

The shared model's projection onto the target coordinates, and its joint zero slice, are

$$\mathcal{P}_{\mathrm{sh}} = \mathrm{proj}_x\, \mathcal{F}_{\mathrm{sh}} = \{x : \exists (h, a) \text{ with } (x, h, a) \in \mathcal{F}_{\mathrm{sh}}\}, \qquad \mathcal{Z}_{\mathrm{sh}} = \mathrm{proj}_x(\mathcal{F}_{\mathrm{sh}} \cap \{(x, h, a) : h = 0_n\}). \tag{N3.6}$$

##### N3.2 The joint zero-slice condition (Z)

The statement "the zero-exposure submodel embeds into the shared model through $h = 0$" is made precise by the following condition, which is both necessary and sufficient for that embedding:

$$\forall x \in \mathcal{X}^{(0)}, \quad \exists\, a_x \in \mathcal{A} \quad \text{such that} \quad (x, 0_n, a_x) \in \mathcal{F}_{\mathrm{sh}}; \tag{Z}$$

equivalently, $\mathcal{X}^{(0)} \subseteq \mathcal{Z}_{\mathrm{sh}}$. The profile $a_x$ must be one and the same across all $n$ samples for the given joint witness $x = (x_1, \ldots, x_n)$, but it may vary with the witness.

Condition (Z) is deliberately joint. Coordinatewise admissibility of zero exposure ($0 \in H_i$ for every $i$) implies (Z) when $\mathcal{H} = \prod_i H_i$ and no other cross-sample or $x$–$h$–$a$ coupling is present; under joint constraints, coordinatewise zeros need not make the all-zero vector $0_n$ feasible (Sect. N3.6). Condition (Z) is also specifically the condition for embedding *through the zero slice*: the weaker projection inclusion $\mathcal{X}^{(0)} \subseteq \mathcal{P}_{\mathrm{sh}}$ could in principle hold through lifts with $h \neq 0_n$, so (Z) is sufficient but not necessary for that inclusion; it is necessary and sufficient for the zero-slice embedding that the Proposition uses.

##### N3.3 The Proposition and its proof

**Proposition (shared completion structure; replication without compelled exposure is not identification).** Let $\varnothing \neq \mathcal{X}^{(0)} = \prod_{i=1}^{n} \mathcal{X}_i^{(0)}$ be the joint feasible class under the zero-exposure submodel, and let $\mathcal{F}_{\mathrm{sh}} \subseteq \mathcal{X} \times \mathcal{H} \times \mathcal{A}$ be a model with a cohort-shared completion profile. Assume condition (Z). Then

$$\mathcal{X}^{(0)} \subseteq \mathrm{proj}_x\, \mathcal{F}_{\mathrm{sh}}. \tag{N3.7}$$

Consequently, for every scalar target $\psi : \mathcal{X} \to \overline{\mathbb{R}}$,

$$\inf_{x\in\operatorname{proj}_x \mathcal{F}_{\rm sh}} \psi(x) \;\le\; \inf_{x\in\mathcal{X}^{(0)}} \psi(x), \qquad \sup_{x\in\operatorname{proj}_x \mathcal{F}_{\rm sh}} \psi(x) \;\ge\; \sup_{x\in\mathcal{X}^{(0)}} \psi(x). \tag{N3.8}$$

Thus replication through a shared profile cannot remove any feasible target value, extremal witness, or extremal sequence inherited from the zero-exposure submodel: the shared-profile constraint tightens no bound inherited from that submodel.

**Proof.** Fix any $x \in \mathcal{X}^{(0)}$. By (Z) there is a single profile $a_x \in \mathcal{A}$, shared across all $n$ samples, such that $(x, 0_n, a_x) \in \mathcal{F}_{\rm sh}$. By the definition of the projection, $x \in \operatorname{proj}_x \mathcal{F}_{\rm sh}$. Since $x$ was arbitrary, the inclusion (N3.7) holds. For any scalar $\psi$, the infimum is antitone and the supremum is monotone under set inclusion; applying these order properties to (N3.7) gives (N3.8). If an endpoint over $\mathcal{X}^{(0)}$ is attained, its optimizer itself remains feasible in the shared model; if it is not attained, every extremal sequence from $\mathcal{X}^{(0)}$ remains a feasible sequence in the shared projection. □

**Corollary N3.1 (identified sets).** Writing $\mathcal{I}_0(\psi) = \{\psi(x) : x \in \mathcal{X}^{(0)}\}$ and $\mathcal{I}_{\rm sh}(\psi) = \{\psi(x) : x \in \mathcal{P}_{\rm sh}\}$, the inclusion (N3.7) gives

$$\mathcal{I}_0(\psi) \subseteq \mathcal{I}_{\rm sh}(\psi). \tag{N3.9}$$

In particular, if the zero-exposure submodel already contains a pair of feasible witnesses with $\psi(x^-) < 0 < \psi(x^+)$, the shared model contains the same pair, and no sign determination can emerge from sharing the profile; and if the zero-exposure target image already fills the maximal admissible range, the shared model's image cannot be narrower. Only the superset relation is asserted; the two identified sets need not be equal.

##### N3.4 The annihilation argument: externally calibrated profiles add nothing on the zero slice

In the additive model (N3.3), $h_i = 0$ makes the product $h_i a$ vanish for every profile $a$. On the zero slice, therefore, the completion profile never enters the data-fit constraints, and the entire content of condition (Z) reduces to joint admissibility of $(x, 0_n, a)$ in $\mathcal{C}$.

This yields the sharpest form of the negative result. Suppose the profile is calibrated externally and exactly, shrinking $\mathcal{A}$ to a single point $\{a^\dagger\}$. If the full zero slice remains admissible, condition (Z) holds with $a_x = a^\dagger$ for every witness, and the Proposition applies verbatim: even fixing the missing profile exactly excludes no zero-exposure witness, because $0 \cdot a^\dagger = 0$. Under (Z), constraints on the missing profile alone remove no zero-exposure witness and therefore move no bound inherited from the zero slice. External calibration of $a$ moves such a bound only when it is coupled to information that forces the hidden type's exposure to appear in the observations; that is, information that deletes the joint zero slice (Sect. N3.6). Accordingly, "the profile is externally calibrated" is neither a counterexample to the Proposition nor, by itself, an escape from it: the dividing line is whether the zero slice survives, not whether $a$ is known.

##### N3.5 A product-form sufficient condition (Z*)

Condition (Z) is stated witness-by-witness. A simpler structural condition implies it in one step. Suppose the non-fit constraints factor as a product,

$$\mathcal{C} = \mathcal{C}_x \times \mathcal{H} \times \mathcal{A}, \qquad \mathcal{X}^{(0)} \subseteq \mathcal{C}_x, \qquad 0_n \in \mathcal{H}, \qquad \mathcal{A} \neq \varnothing. \tag{Z*}$$

Then a single uniform profile works for all witnesses: pick any $a_\star \in \mathcal{A}$; for every $x \in \mathcal{X}^{(0)}$, the point $(x, 0_n, a_\star)$ satisfies the product constraint by construction and satisfies the data-fit constraints because $h = 0_n$ reduces them to the zero-exposure constraints that $x$ already satisfies. Hence (Z) holds, in the stronger, uniform-profile form $\exists a_\star\, \forall x$, whenever (Z*) holds. Condition (Z*) is what typical "unstructured completion" model classes satisfy: no coupling between the target coordinates, the exposure vector, and the profile beyond the data-fit terms, exposures free to vanish jointly, and at least one admissible profile.

Under (Z*) with $\mathcal{H} = \prod_i H_i$, the coordinatewise condition $0 \in H_i$ for all $i$ is equivalent to $0_n \in \mathcal{H}$; the product form is sufficient for this equivalence, not necessary.

##### N3.6 Boundary: which assumption classes do delete the zero slice

The Proposition's zero-slice guarantee ceases where its hypothesis fails: when the joint zero slice is deleted, some zero-exposure witness admits no shared-profile lift with $h = 0_n$, and projection inclusion, if it holds at all, then requires a separate proof through nonzero-exposure lifts. The following assumption classes delete the zero slice. For each, the conclusion of the Proposition no longer applies and the class may contract the bounds, but whether contraction actually occurs is a separate question requiring its own proof in each model.

1. **Per-sample exposure floors.** Requiring $h_i \geq \eta > 0$ for all $i$ (or for sufficiently many samples) makes $0_n$ infeasible. A worked example shows the resulting contraction can be strict and exactly quantified. Take all quantities scalar: $y_i = 1$, $A_{-d} = 1$, $\mathcal{A} = \{1\}$, with the exact-fit and mass constraint $1 = x_i + h_i$ and $x_i, h_i \geq 0$. The zero-exposure submodel is the single point $\mathcal{X}^{(0)} = \{(1, \ldots, 1)\}$, so for the target $\psi(x) = n^{-1} \sum_i x_i$ the zero-exposure sharp upper bound is 1. Imposing $h_i \geq \eta > 0$ with $0 < \eta \leq 1$ (feasibility requires $\eta \leq 1$, since $x_i = 1 - h_i \geq 0$) forces $x_i \leq 1 - \eta$ for every sample, and the bound $\psi \leq 1 - \eta$ is attained at $x_i = 1 - \eta$: the exposure floor lowers the sharp upper bound from 1 to $1 - \eta$. This is the sufficient example cited in the main text and in the Fig. 6C legend.
2. **Cross-sample exposure constraints.** Aggregate lower bounds such as $n^{-1} \sum_i h_i \geq \eta > 0$, weighted versions $\sum_i w_i h_i = c > 0$, or covariate models forcing a positive exposure scale delete $0_n$ from $\mathcal{H}$ even when every coordinate marginally admits zero. In the worked example above, replacing the per-sample floor by the average-exposure bound $n^{-1} \sum_i h_i \geq \eta$ yields the same strict contraction of the upper bound.
3. **Hidden-type-specific anchors and absolute-mass constraints.** External measurements that bear on the hidden type itself (an anchor that measures the hidden type's abundance, or an absolute-mass constraint tying total measured material to the completed composition) couple the observations to $h$ and can make $h = 0_n$ inconsistent with the anchor values.
4. **Joint constraints on $(a, h, x)$.** Calibration, moment, rank, normalization or transport conditions that require $\sum_i h_i a$ to be nonzero, or that leave some $x \in \mathcal{X}^{(0)}$ with no admissible common profile, violate (Z) directly. Sharing $a$ across samples does not by itself create such a failure; an additional constraint must actually delete the zero slice.
5. **Changed baseline constraints.** If the shared model simultaneously tightens the residual budgets, adds smoothing, regression or grouping constraints across the $x_i$, changes the normalization, or changes the estimand, then the zero-exposure product class need not embed; but any resulting contraction is attributable to those added restrictions, not to profile sharing itself.

Two further scope limits apply on the conclusion side rather than the hypothesis side. The target must depend only on the projected coordinates $x_{1:n}$: the projection inclusion transfers bounds for $\psi(x)$, not for functionals of the profile shape, the hidden mass, or joint functionals of $(x, h, a)$. And the comparison must concern feasible target sets or their endpoints, not solver incumbents, approximate outer envelopes, or statistical precision, to which the set-inclusion argument does not speak.

#### N3.7 Strictly positive but vanishing exposure: a closure version

If the model requires $h_i > 0$ with no uniform lower bound, the exact witness embedding no longer applies, since $h = 0_n$ is infeasible. A closure version survives: if

$$\mathcal{X}^{(0)} \subseteq \overline{\mathcal{P}_{\mathrm{sh}}} \tag{N3.10}$$

and $\psi$ is continuous, then the endpoint inequalities (N3.8) continue to hold, by taking for each zero-exposure witness a convergent sequence of feasible shared-model points and passing to the limit. The closure version preserves endpoints and strict-margin sign witnesses, but not the statement that the original witness is itself feasible.

The closure hypothesis (N3.10) must be verified, not assumed: requiring merely $h_i > 0$ does not automatically imply it. Normalization, non-negativity boundaries, equality constraints or cross-sample coupling can all prevent zero-exposure witnesses from being approximated by strictly positive-exposure feasible points.

#### N3.8 Relation to Theorems 1 and 2

The Proposition is logically independent of Theorems 1 and 2 (Additional file 2: Note 1), and is not a corollary of either.

Theorem 1 is an existence construction: it exhibits a specific single-sample observation whose identified set of retained-type conditional compositions fills the open simplex. Its admissible worlds carry strictly positive hidden mass, so they do not supply the zero slice that condition (Z) requires, and its realizing family varies the completion profile with the target, so it does not assemble into a shared-profile multi-sample witness. Conversely, the Proposition constructs no observation and controls no conditioning: it is a set-monotonicity statement for arbitrary $n$, arbitrary zero-exposure submodels and arbitrary scalar targets, contingent on (Z). The only valid bridge is conditional: if some baseline class with Theorem-1-type ambiguity is separately proven to embed into a shared model's projection, the ambiguity transfers by inclusion; and it is the separately proven inclusion, not Theorem 1, that carries the argument.

With respect to Theorem 2, the inclusion (N3.7) transfers *lower* bounds only: risk or regret witnesses feasible in the zero-exposure submodel remain feasible in the shared model, so worst-case quantities cannot decrease; but the shared class may contain additional worlds, so Theorem 2's exact minimax equalities do not transfer by inclusion alone.

##### N3.9 Scope, and the position of the PBMC pilot

The Proposition applies to comparisons in which (i) the baseline is genuinely the zero-exposure joint product class $\mathcal{X}^{(0)} = \prod_i \mathcal{X}_i^{(0)}$; (ii) the shared and zero-exposure models use the same data-acceptance rules, residual budgets and meaning of $x$ at $h = 0_n$; (iii) condition (Z) holds for every joint witness in the zero class, not merely for one feasible point; (iv) the target depends only on $x_{1:n}$; and (v) the objects compared are feasible target sets or their endpoints. It does not assert that more samples never help, that all cohort-shared completion models are non-identifying, or that multi-sample designs are useless: wherever the zero slice is deleted (Sect. N3.6), the question of contraction is reopened and must be settled by its own analysis.

The PBMC pilot reported in the main text and Methods is an empirical instantiation in one real cohort, logically separate from the Proposition: the pilot's zero-exposure-floor scenario is consistent with condition (Z) and exhibits the predicted absence of contraction, while its positive-floor scenarios lie outside the Proposition's hypothesis; their observed non-contraction is an additional empirical failure of that model class, not a consequence of this Note. The pilot's regression checks that the shared class contains the baseline class at zero exposure floor are implementation tests of (Z), not part of the proof. The pilot's computational closure is bounded: of its 384 optimization extrema, 273 were solved, 107 remained unresolved at the prespecified gap and 4 were infeasible, and all 36 main-profile extrema of its rank-one class carried gaps above the threshold; its incumbents are feasible sign witnesses, not sharp global extrema. Conversely, nothing in the pilot, including those unresolved cells reported as computational bounds, enters the mathematical argument above.

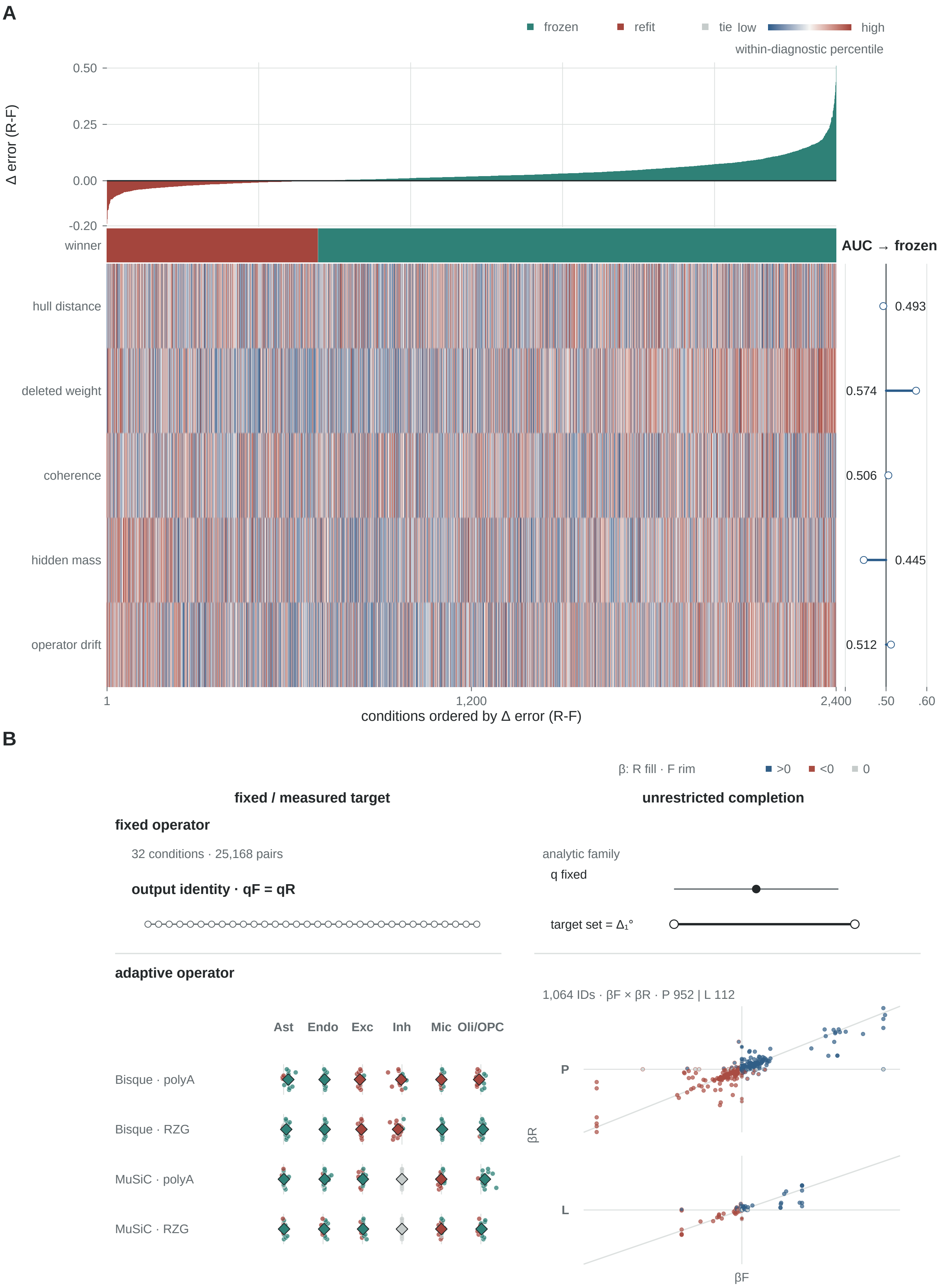
A
frozen
refit
tie
low
high
within-diagnostic percentile
Δ error (R-F)
0.50
0.25
0.00
-0.20
winner
AUC → frozen
hull distance
deleted weight
coherence
hidden mass
operator drift
0.493
0.574
0.506
0.445
0.512
1
1,200
2,400
.50
.60
conditions ordered by Δ error (R-F)
B
β: R fill · F rim
>0
<0
0
fixed / measured target
unrestricted completion
fixed operator
32 conditions · 25,168 pairs
output identity · qF = qR
analytic family
q fixed
target set = Δ₁°
adaptive operator
1,064 IDs · βF × βR · P 952 | L 112
Ast
Endo
Exc
Inh
Mic
Oli/OPC
Bisque · polyA
Bisque · RZG
MuSiC · polyA
MuSiC · RZG
P
L
βR
βF

#### Fig. S1 caption

**Fig. S1. Common scalar diagnostics show near-chance average-case discrimination.** **(A)** The synthetic deletion spectrum comprises 2,400 deletion conditions (400 dictionaries × six deletions) evaluated by five scalar diagnostics: hull distance, deleted weight, coherence, hidden mass and operator drift. Conditions are ordered by Δ error $(R - F)$, positive when frozen has the lower error (71% of conditions) and negative when refit does (29%). Areas under the receiver operating characteristic curve are 0.493 for hull distance, 0.574 for deleted weight, 0.506 for coherence, 0.445 for hidden mass and 0.512 for operator drift, where 0.5 is chance. Teal denotes frozen-favoring and brick red refit-favoring conditions; the heat field is the within-diagnostic percentile. In the side strip, the open circle is the AUC estimate and the horizontal line runs from 0.5 to it as a reading aid, not an interval. **(B)** The $2 \times 2$ panel is a logical index that reuses real fixed-operator, adaptive-operator, measured-target and unrestricted-completion objects; it creates no new sample size. Fixed output identity establishes an operator seam but does not identify the target. For association effects, refit is the fill and frozen the rim; blue denotes positive $\beta$, orange negative $\beta$ and gray zero, distinct from the winner key in A. In the adaptive-operator quadrant, the 24 cells are four estimator-by-preparation rows (BisqueRNA and MuSiC, each with polyA and RiboZeroGold) crossed with six deleted types, and the 336 pairs are the 14 frozen-versus-refit paired observations behind each cell. Each small circle is one paired Δ loss, displaced vertically only to avoid overlap; each diamond is the cell median; the short vertical tick marks $\Delta = 0$. Fill follows the winner key. Ast, astrocyte; AUC, area under the curve; Endo, endothelial/mural; Exc, excitatory neuron; Inh, inhibitory neuron; Mic, microglia; Oli/OPC, oligodendrocyte/oligodendrocyte precursor cell; RZG, RiboZeroGold.

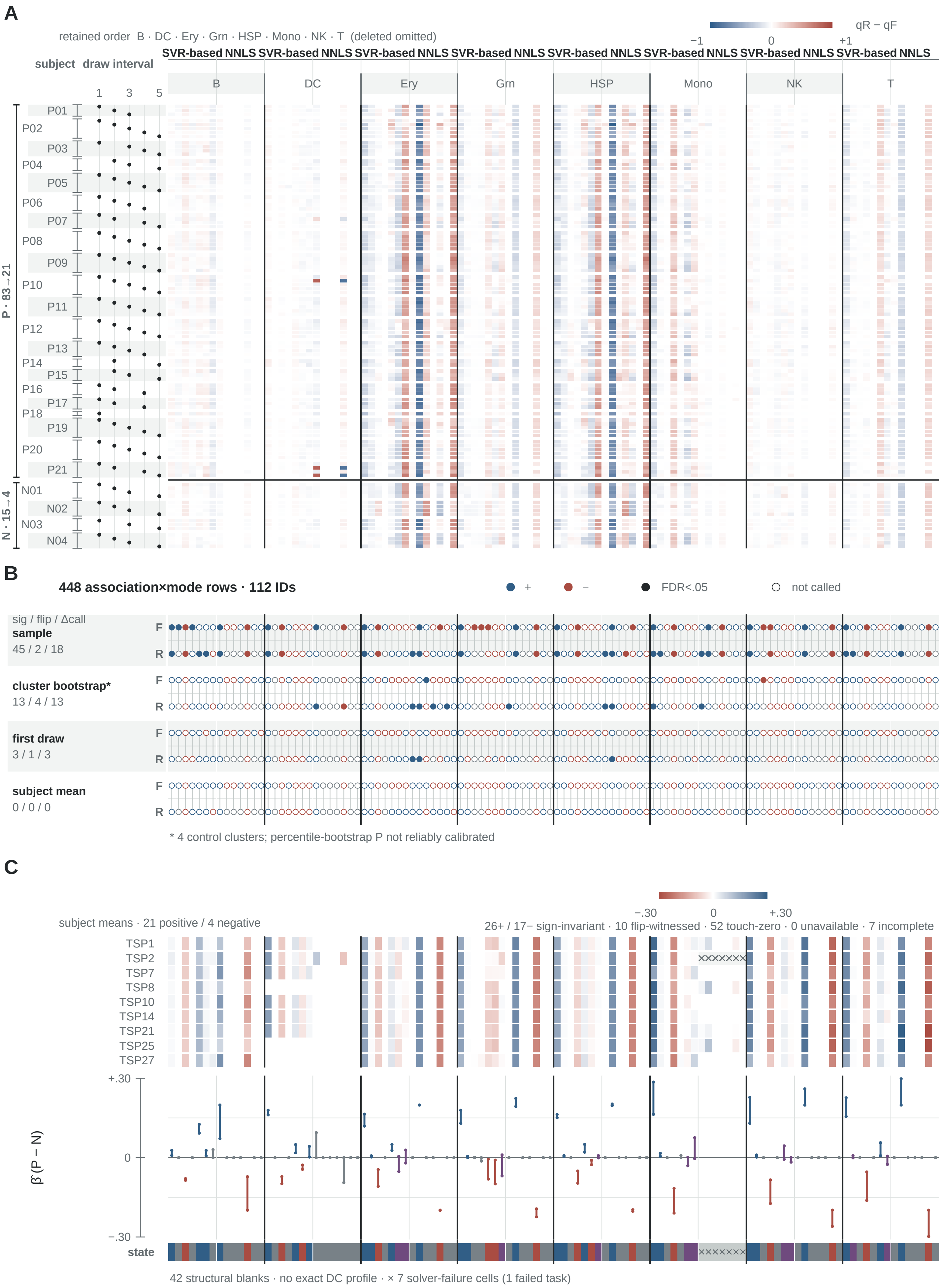
A
retained order B · DC · Ery · Grn · HSP · Mono · NK · T (deleted omitted)
qR − qF
−1
0
+1
SVR-based NNLS
subject draw interval
1 3 5
B
DC
Ery
Grn
HSP
Mono
NK
T
P01
P02
P03
P04
P05
P06
P07
P08
P09
P10
P11
P12
P13
P14
P15
P16
P17
P18
P19
P20
P21
N01
N02
N03
N04
P · 83 → 21
N · 15 → 4
B
448 association×mode rows · 112 IDs
+
−
FDR<.05
not called
sig / flip / Δcall
sample
45 / 2 / 18
cluster bootstrap*
13 / 4 / 13
first draw
3 / 1 / 3
subject mean
0 / 0 / 0
F
R
* 4 control clusters; percentile-bootstrap P not reliably calibrated
C
subject means · 21 positive / 4 negative
−.30
0
+.30
26+ / 17− sign-invariant · 10 flip-witnessed · 52 touch-zero · 0 unavailable · 7 incomplete
TSP1
TSP2
TSP7
TSP8
TSP10
TSP14
TSP21
TSP25
TSP27
+.30
0
−.30
β̃ (P − N)
state
42 structural blanks · no exact DC profile · × 7 solver-failure cells (1 failed task)

#### Fig. S2 caption

**Fig. S2. Repeated draws do not constitute a tenth independent cohort. (A)** Frozen-to-refit composition displacement in longitudinal GSE166190 is shown as $(q_R - q_F)$ for 10,976 paired cells across 98 draws from 25 subjects, comprising 21 positive-group and four negative-group subjects. Columns follow deletion, estimator and retained-type order. **(B)** Outcomes for the same 112 association IDs are compared under sample-level analysis, subject-cluster percentile bootstrap, first-draw-only analysis and subject-mean aggregation. Registers give significant associations, direction flips (opposite signs with at least one path significant) and significance-status changes: 45/2/18, 13/4/13, 3/1/3 and 0/0/0, respectively. Percentile-bootstrap calibration is unreliable with four negative-group clusters, so two subject-level exact tests were also run and are reported in Additional file 1: Table S3: permutation of the phenotype labels over all 12,650 complete assignments of four negative labels among the 25 subjects gives 27/2/13, and the size-stratified clustered Wilcoxon exact test (336 assignments) gives 0/0/0. **(C)** After subject-mean aggregation, 959 profile effects form 112 envelopes: 105 complete, comprising 26 positive and 17 negative sign-invariant, 10 flip-witnessed, 52 touch-zero and zero unavailable, plus seven incomplete. Forty-two structural blanks denote the absence of an exact dendritic-cell profile. One failed task expands to seven solver-failure cells; none is imputed as zero. This subject-mean supplementary analysis is not pooled with the 12,204 profile effects of the nine-cohort primary analysis. Blue and orange denote positive and negative association effects, and purple denotes witnessed flips. The blue–white–brick field in A encodes signed $(q_R - q_F)$ displacement, with positive values meaning larger refit proportions, not path superiority. B, B cell; DC, dendritic cell; Ery, erythroid cell; FDR, false discovery rate; Grn, granulocyte; HSP, hematopoietic stem/progenitor cell; Mono, monocyte; NK, natural killer cell; T, T cell.

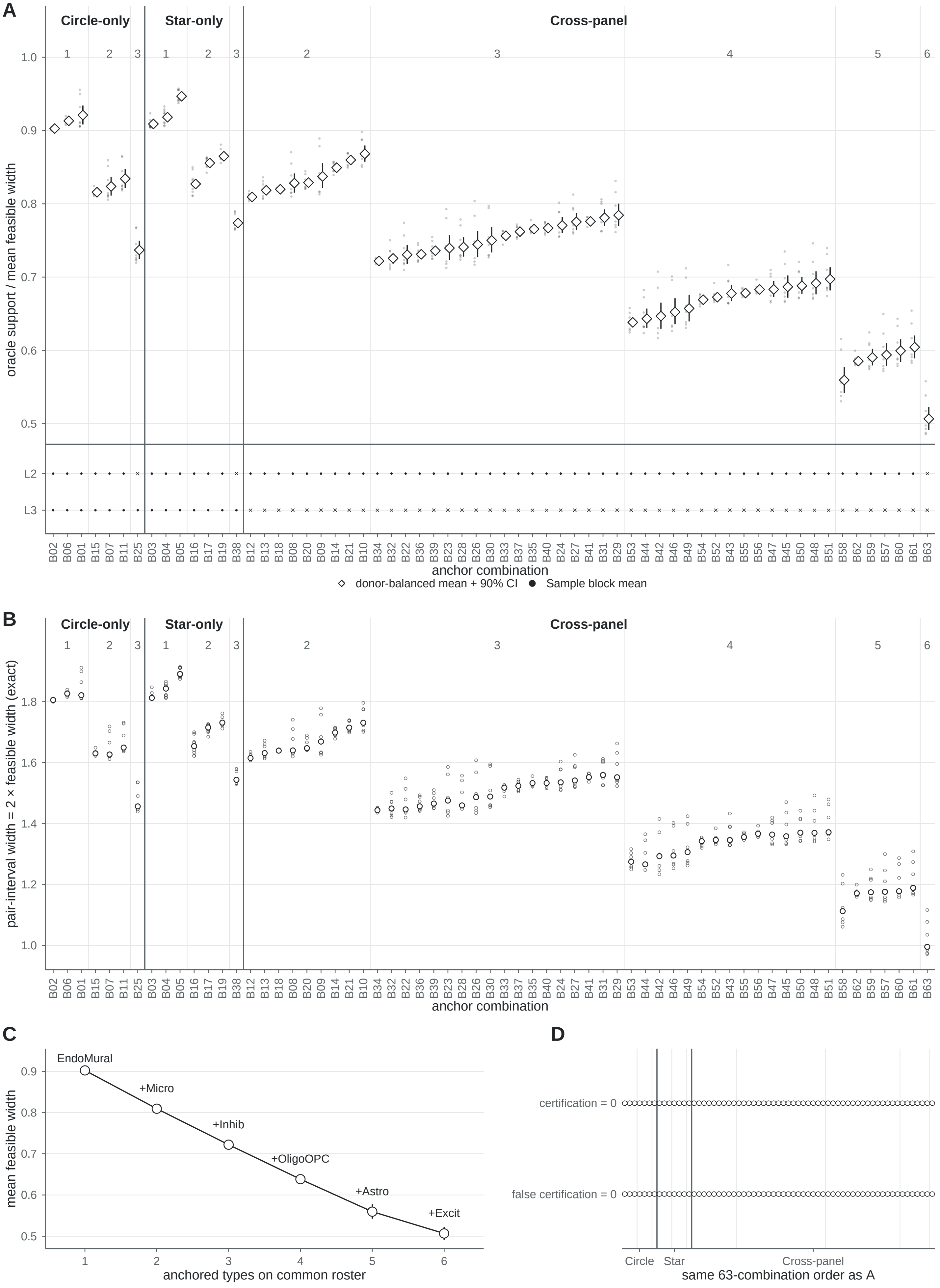
A
Circle-only
Star-only
Cross-panel
oracle support / mean feasible width
L2
L3
anchor combination
donor-balanced mean + 90% CI
Sample block mean
B
pair-interval width = 2 × feasible width (exact)
anchor combination
C
mean feasible width
EndoMural
+Micro
+Inhib
+OligoOPC
+Astro
+Excit
anchored types on common roster
D
certification = 0
false certification = 0
Circle
Star
Cross-panel
same 63-combination order as A

#### Fig. S3 caption

**Fig. S3. The anchor-combination field: a stable value ordering, and a verifiable depth of three types. (A)** All 63 anchored-type combinations are shown by number of anchored types (one to six) and panel scope (Circle-only 3/3/1, Star-only 3/3/1 and cross-panel 0/9/18/15/6/1 at k = 1…6), each with its mean feasible width, 90% donor-bootstrap interval and by-block records; the strip below marks oracle support per combination. Exactly 14 combinations (seven Circle-only and seven Star-only, no cross-panel combination) admit the strongest held-out imaging check, because only single-panel anchoring leaves a complete complementary panel to verify against. **(B)** Pair-interval widths for the same 63 combinations: each plotted point is one Sample block, so the spread is between blocks within a combination, and the vertical axis is on the pairwise-difference scale, exactly twice the coordinate feasible-width scale of the other panels. **(C)** The marginal-value ordering on the common roster of 7 blocks and 6 donors, one type added at a time; it is a width-reduction ordering, not a statement of biological importance, and combinations measured on different rosters are not ranked against one another. **(D)** Certification and false certification are zero for all 63 combinations over the full prespecified tolerance range.

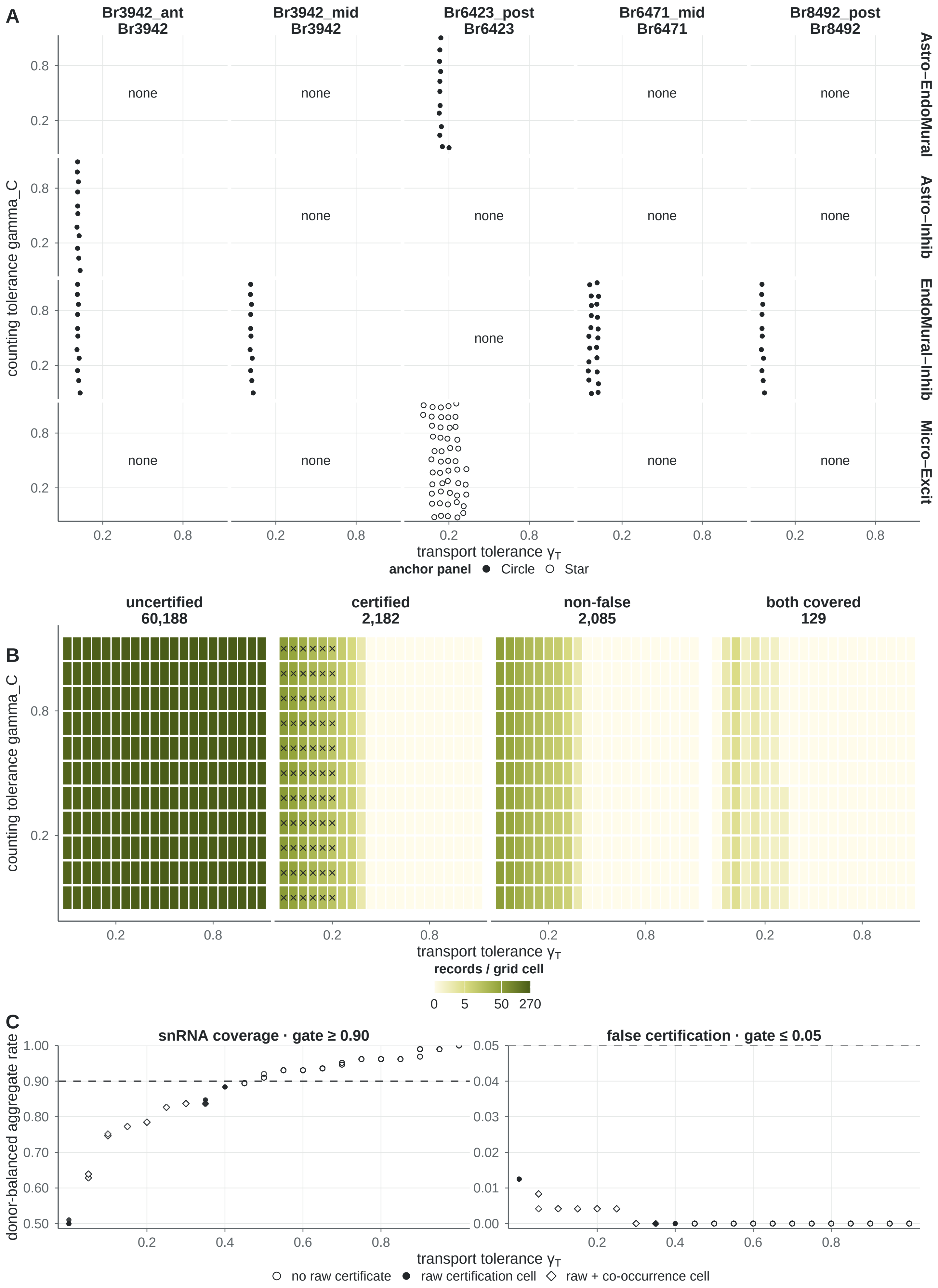

A
Br3942_ant Br3942
Br3942_mid Br3942
Br6423_post Br6423
Br6471_mid Br6471
Br8492_post Br8492
Astro–EndoMural
Astro–Inhib
EndoMural–Inhib
Micro–Excit
none
counting tolerance gamma_C
transport tolerance γT
anchor panel ● Circle ○ Star
B
uncertified 60,188
certified 2,182
non-false 2,085
both covered 129
counting tolerance gamma_C
transport tolerance γT
records / grid cell
0 5 50 270
C
snRNA coverage · gate ≥ 0.90
false certification · gate ≤ 0.05
donor-balanced aggregate rate
transport tolerance γT
○ no raw certificate ● raw certification cell ◇ raw + co-occurrence cell

#### Fig. S4 caption

**Fig. S4. Why the certification–coverage intersection is empty at the aggregate level, and what survives beneath it. (A)** The 129 certified records whose two type coordinates are both covered by the held-out snRNA check, resolved by sample, donor, type pair and grid point; they fall in 5 samples, 4 donors, 71 grid points and 4 type-pair identities over transport tolerances 0.05 to 0.35, with anchor panel marked. **(B)** Record accounting over the tolerance grid: of 62,370 sign records, 2,182 are certified, 2,085 of these are not false certifications, and 129 have both types covered. **(C)** Aggregate validity after donor- and pair-balanced aggregation across samples, donors and all 15 type-pair identities: snRNA coverage against its 0.90 threshold and false certification against its 0.05 threshold, by transport tolerance. The aggregate is zero because the record-level co-occurrences neither span enough type-pair identities nor recur consistently across donors, so the 129 records are not valid certifications; the empty intersection is a statement about the aggregate, not a claim that every record fails. Records that are unavailable or structurally undefined are drawn as explicit states and never as zero; libraries, panel instances and grid rows are not independent sample sizes.
