## Additional file 3: Data S1 for "Incomplete references leave bulk deconvolution targets non-identifiable, but identification has a computable precision price": README.md

# Supplementary Data S1

This archive contains the exact-arithmetic theorem verifier and the single matching standard output captured from a fresh run on 2026-08-09.

## Files

`verify_theorem_20260808.py` is the verification script and uses only the Python standard library. `verify_theorem_20260808.txt` is the matching run output with 80 `PASS |` lines and final line `ALL CHECKS PASS | checks=80`.

## Reproduction

```bash
python3 verify_theorem_20260808.py > verify_theorem_20260808.txt
```

## SHA-256

- `verify_theorem_20260808.py`: `43c0855ff21c92a0f4868480e67f1e34926da7a5de74411d4b7711bf44bb5901`
- `verify_theorem_20260808.txt`: `0af4800da59a137a8c100a67cced50a7261e80ffa4fd517680c98f3db533e03f`
