## Additional file 3: Data S1 for "Incomplete references leave bulk deconvolution targets non-identifiable, but identification has a computable precision price": verify_theorem_20260808.py

```
#!/usr/bin/env python3
"""Exact verification for THEOREM_NOTE_2026-08-08.md.

The script performs exact-arithmetic checks of the displayed witnesses,
constants, finite lattice points, and representative embeddings. Universal
claims are proved analytically in the note. Decimal strings are display-only
conversions of already checked exact rationals.
"""

from fractions import Fraction as F


ZERO, ONE = F(0), F(1)
PASS_COUNT = 0


def pass_check(label, condition):
    global PASS_COUNT
    if not condition:
        raise AssertionError(label)
    PASS_COUNT += 1
    print(f"PASS | {label}")


def fmt(q, digits=6):
    return f"{float(q):.{digits}f}"


def transpose(a):
    return [list(row) for row in zip(*a)]


def matmul(a, b):
    bt = transpose(b)
    return [[sum((x * y for x, y in zip(row, col)), ZERO) for col in bt] for row in a]


def matvec(a, x):
    return [sum((u * v for u, v in zip(row, x)), ZERO) for row in a]


def columns(*cols):
    return [list(row) for row in zip(*cols)]


def inverse(a):
    n = len(a)
    aug = [row[:] + [F(int(i == j)) for j in range(n)] for i, row in enumerate(a)]
    for col in range(n):
        pivot = next(row for row in range(col, n) if aug[row][col] != 0)
        aug[col], aug[pivot] = aug[pivot], aug[col]
        scale = aug[col][col]
        aug[col] = [x / scale for x in aug[col]]
        for row in range(n):
            if row != col:
                scale = aug[row][col]
                aug[row] = [x - scale * y for x, y in zip(aug[row], aug[col])]
    return [row[n:] for row in aug]


def determinant(a):
    n = len(a)
    work = [row[:] for row in a]
    value = ONE
    for col in range(n):
        pivot = next((row for row in range(col, n) if work[row][col] != 0), None)
        if pivot is None:
            return ZERO
        if pivot != col:
            work[col], work[pivot] = work[pivot], work[col]
            value = -value
        pivot_value = work[col][col]
        value *= pivot_value
        work[col] = [x / pivot_value for x in work[col]]
        for row in range(col + 1, n):
            scale = work[row][col]
            work[row] = [x - scale * y for x, y in zip(work[row], work[col])]
    return value


def norm1(a):
    return max(sum((abs(a[i][j]) for i in range(len(a))), ZERO) for j in range(len(a[0])))


def cond1(a):
    return norm1(a) * norm1(inverse(a))


def matrix_rank(a):
    work = [row[:] for row in a]
    rows, cols = len(work), len(work[0])
    pivot_row = 0
    for col in range(cols):
        pivot = next((row for row in range(pivot_row, rows) if work[row][col] != 0), None)
        if pivot is None:
            continue
        work[pivot_row], work[pivot] = work[pivot], work[pivot_row]
        scale = work[pivot_row][col]
        work[pivot_row] = [x / scale for x in work[pivot_row]]
        for row in range(rows):
            if row != pivot_row and work[row][col] != 0:
                scale = work[row][col]
                work[row] = [x - scale * y for x, y in zip(work[row], work[pivot_row])]
        pivot_row += 1
        if pivot_row == rows:
            break
    return pivot_row


def tv(x, y):
    return sum((abs(a - b) for a, b in zip(x, y)), ZERO) / 2


def conditional_retained(weights, retained_count=2):
    denom = sum(weights[:retained_count], ZERO)
    return [x / denom for x in weights[:retained_count]]


def all_positive(x):
    return all(v > 0 for v in x)


def flatten(a):
    return [x for row in a for x in row]


def lift_matrix(feature_count):
    eps = F(1, 2 * feature_count)
    return [
        [eps + (F(1, 2) if row == col else ZERO) for col in range(3)]
        for row in range(feature_count)
    ]


def global_smoothing(feature_count):
    eps = F(1, 2 * feature_count)
    return [
        [eps + (F(1, 2) if row == col else ZERO) for col in range(feature_count)]
        for row in range(feature_count)
    ]


def solve_from_completion(z):
    """Solve [B_OBS,z]w=Y via the exact preimage under S."""
    a = matvec(S_INV, z)
    hidden = F(1, 2) / a[2]
    return [F(3, 10) - a[0] * hidden, F(1, 5) - a[1] * hidden, hidden]


print("== C1: strictly positive robust witness ==")
S = [
    [F(49, 50), F(1, 100), F(1, 100)],
    [F(1, 100), F(49, 50), F(1, 100)],
    [F(1, 100), F(1, 100), F(49, 50)],
]
S_INV = inverse(S)
E1, E2, E3 = [ONE, ZERO, ZERO], [ZERO, ONE, ZERO], [ZERO, ZERO, ONE]
BASE_Y = [F(3, 10), F(1, 5), F(1, 2)]
BASE_Z2 = [F(3, 10), F(1, 10), F(3, 5)]
B1, B2 = matvec(S, E1), matvec(S, E2)
Y = matvec(S, BASE_Y)
Z1, Z2 = matvec(S, E3), matvec(S, BASE_Z2)
W1 = [F(3, 10), F(1, 5), F(1, 2)]
W2 = [F(1, 20), F(7, 60), F(5, 6)]
FULL1, FULL2 = columns(B1, B2, Z1), columns(B1, B2, Z2)

pass_check("column-stochastic smoothing matrix is invertible", matmul(S_INV, S) == columns(E1, E2, E3))
pass_check("all observed-reference, bulk, completion, and weight entries are strictly positive",
           all_positive(flatten(columns(B1, B2))) and all_positive(Y + Z1 + Z2 + W1 + W2))
pass_check("all reference/completion columns and bulk sum to one",
           all(sum(v, ZERO) == ONE for v in [B1, B2, Z1, Z2, Y]))
pass_check("both mixing vectors lie in the simplex interior",
           all_positive(W1 + W2) and sum(W1, ZERO) == ONE and sum(W2, ZERO) == ONE)
pass_check("both completed worlds reconstruct the identical observed bulk exactly",
           matvec(FULL1, W1) == Y and matvec(FULL2, W2) == Y)

T1, T2 = conditional_retained(W1), conditional_retained(W2)
pass_check("retained conditional compositions are exactly (3/5,2/5) and (3/10,7/10)",
           T1 == [F(3, 5), F(2, 5)] and T2 == [F(3, 10), F(7, 10)])
pass_check("the retained-type ranking reverses", T1[0] > T1[1] and T2[0] < T2[1])
print(f"CONST | Y={tuple(Y)} Z1={tuple(Z1)} Z2={tuple(Z2)}")
print(f"CONST | T1={tuple(T1)} T2={tuple(T2)} TV(T1,T2)={tv(T1, T2)}")

print("\n== Conditioning ==")
GRAM_OBS = matmul(transpose(columns(B1, B2)), columns(B1, B2))
pass_check("observed-reference Gram matrix has the displayed exact entries",
           GRAM_OBS == [[F(4803, 5000), F(197, 10000)],
                        [F(197, 10000), F(4803, 5000)]])
GRAM_DIAGONAL, GRAM_OFF_DIAGONAL = GRAM_OBS[0]
KAPPA2_SQUARED = ((GRAM_DIAGONAL + GRAM_OFF_DIAGONAL)
                  / (GRAM_DIAGONAL - GRAM_OFF_DIAGONAL))
pass_check("observed-reference squared 2-norm condition number is directly 9803/9409",
           KAPPA2_SQUARED == F(9803, 9409))
K1_FULL1, K1_FULL2 = cond1(FULL1), cond1(FULL2)
pass_check("completed-reference 1-norm condition numbers are 101/97 and 233/97",
           K1_FULL1 == F(101, 97) and K1_FULL2 == F(233, 97))
print(f"CONST | kappa2(A_obs)=sqrt(9803/9409)={fmt((F(9803, 9409)) ** F(1, 2))}")
print(f"CONST | kappa1(A_plus_1)={K1_FULL1}={fmt(K1_FULL1)}")
print(f"CONST | kappa1(A_plus_2)={K1_FULL2}={fmt(K1_FULL2)}")

print("\n== Open-neighborhood persistence ==")
RHO = F(1, 200)
E = F(100, 97) * RHO
W1_P1_LOWER = F(3, 10) - E / (2 * (1 - E))
W1_P2_LOWER = F(1, 5) - E / (2 * (1 - E))
W1_MARGIN_LOWER = F(1, 10) - E / (1 - E)
W2_P1_LOWER = F(3, 10) - F(1, 2) * (F(3, 10) + E) / (F(3, 5) - E)
W2_P2_LOWER = F(1, 5) - F(1, 2) * (F(1, 10) + E) / (F(3, 5) - E)
W2_MARGIN_LOWER = F(1, 2) * (F(1, 5) - 2 * E) / (F(3, 5) + E) - F(1, 10)
U1_CLOSURE_K1_BOUND = F(195, 193) * F(101, 97)
U2_CLOSURE_K1_BOUND = F(1363, 577) * F(101, 97)
U1_STATED_K1_BOUND = F(19796, 18721)
U2_STATED_K1_BOUND = F(138168, 55969)
pass_check("rho=1/200 gives exact positive coefficient and rank-margin lower bounds",
           [W1_P1_LOWER, W1_P2_LOWER, W1_MARGIN_LOWER,
            W2_P1_LOWER, W2_P2_LOWER, W2_MARGIN_LOWER]
           == [F(287, 965), F(381, 1930), F(183, 1930),
               F(251, 5770), F(322, 2885), F(333, 5870)])
pass_check("closure 1-norm bounds are strictly below the stated neighborhood bounds",
           U1_CLOSURE_K1_BOUND == F(19695, 18721)
           and U2_CLOSURE_K1_BOUND == F(137663, 55969)
           and U1_CLOSURE_K1_BOUND < U1_STATED_K1_BOUND
           and U2_CLOSURE_K1_BOUND < U2_STATED_K1_BOUND)


def scan_neighborhood(center, expected_first_wins):
    tested = 0
    step = F(1, 4000)
    for j in range(-19, 20):
        for k in range(-19, 20):
            h = [j * step, k * step, -(j + k) * step]
            if max(map(abs, h)) >= RHO:
                continue
            z = [x + dx for x, dx in zip(center, h)]
            weights = solve_from_completion(z)
            target = conditional_retained(weights)
            exact_reconstruction = matvec(columns(B1, B2, z), weights)
            if not (all_positive(z + weights) and sum(z, ZERO) == ONE
                    and sum(weights, ZERO) == ONE and exact_reconstruction == Y):
                raise AssertionError("inadmissible point in exact neighborhood scan")
            if (target[0] > target[1]) != expected_first_wins:
                raise AssertionError("rank failed in exact neighborhood scan")
            tested += 1
    return tested


SCAN1 = scan_neighborhood(Z1, True)
SCAN2 = scan_neighborhood(Z2, False)
pass_check("exact rational lattice scan preserves admissibility and opposite rankings",
           SCAN1 == 1141 and SCAN2 == 1141)
print(f"CONST | rho_inf={RHO}={fmt(RHO)} preimage_radius={E}")
print(f"CONST | analytic rank margins: U1>{W1_MARGIN_LOWER}, U2>{W2_MARGIN_LOWER}")
print(f"CONST | uniform kappa1 bounds: U1<={U1_CLOSURE_K1_BOUND}"
      f"<{U1_STATED_K1_BOUND}; U2<={U2_CLOSURE_K1_BOUND}"
      f"<{U2_STATED_K1_BOUND}")
print(f"SCAN  | U1={SCAN1} U2={SCAN2} implied_pairs={SCAN1 * SCAN2}")

print("\n== Full identified set in the two-type simplex interior ==")
IDENTIFIED_SET_T_VALUES = [F(1, 7), F(1, 3), F(1, 2), F(2, 3), F(9, 10)]
IDENTIFIED_FAMILY_K1_BOUND = F(13, 5) * F(101, 97)
for t in IDENTIFIED_SET_T_VALUES:
    weights = [t / 10, (ONE - t) / 10, F(9, 10)]
    raw_completion = [(3 - t) / 9, (ONE + t) / 9, F(5, 9)]
    completion = matvec(S, raw_completion)
    full = columns(B1, B2, completion)
    target = conditional_retained(weights)
    pass_check(f"identified-set family t={t}",
               all_positive(raw_completion + completion + weights)
               and sum(raw_completion, ZERO) == ONE
               and sum(completion, ZERO) == ONE
               and sum(weights, ZERO) == ONE
               and matvec(full, weights) == Y
               and target == [t, ONE - t]
               and norm1(inverse(columns(E1, E2, raw_completion))) == F(13, 5)
               and cond1(full) <= IDENTIFIED_FAMILY_K1_BOUND)
pass_check("identified-set family uniform 1-norm condition bound is 1313/485",
           IDENTIFIED_FAMILY_K1_BOUND == F(1313, 485))

print("\n== Arbitrary P-feature by K-retained embedding ==")

pass_check("H_P leading 3-by-3 minor equals (P+3)/(8P) for P=3..8",
           all(determinant([row[:3] for row in lift_matrix(feature_count)[:3]])
                   == F(feature_count + 3, 8 * feature_count)
                   and matrix_rank(lift_matrix(feature_count)) == 3
                   for feature_count in range(3, 9)))


def embedded_world(feature_count, retained_count, completion, base_weights):
    lift = lift_matrix(feature_count)
    mapped_b1, mapped_b2 = matvec(lift, B1), matvec(lift, B2)
    mapped_y, mapped_z = matvec(lift, Y), matvec(lift, completion)
    eta = F(1, 4 * retained_count)
    extra_count = retained_count - 2
    tau = extra_count * eta
    lam = ONE - tau
    observed = [mapped_b1, mapped_b2] + [mapped_y for _ in range(extra_count)]
    weights = [lam * base_weights[0], lam * base_weights[1]]
    weights += [eta for _ in range(extra_count)]
    weights += [lam * base_weights[2]]
    full = columns(*(observed + [mapped_z]))
    target = conditional_retained(weights, retained_count)
    return lift, observed, mapped_y, mapped_z, weights, full, target


EMBED_TESTS = 0
for feature_count in range(3, 9):
    for retained_count in range(2, 8):
        world1 = embedded_world(feature_count, retained_count, Z1, W1)
        world2 = embedded_world(feature_count, retained_count, Z2, W2)
        lift1, obs1, y1, z1, weights1, full1, target1 = world1
        lift2, obs2, y2, z2, weights2, full2, target2 = world2
        pass_check(f"embedding P={feature_count}, K={retained_count}",
                   lift1 == lift2 and obs1 == obs2 and y1 == y2
                   and all_positive(flatten(lift1) + flatten(columns(*obs1)) + y1 + z1 + z2)
                   and all(sum(col, ZERO) == ONE for col in obs1 + [z1, z2, y1])
                   and all_positive(weights1 + weights2)
                   and sum(weights1, ZERO) == ONE and sum(weights2, ZERO) == ONE
                   and matvec(full1, weights1) == y1 and matvec(full2, weights2) == y2
                   and target1[0] > target1[1] and target2[0] < target2[1])
        EMBED_TESTS += 1
print(f"SCAN  | exact embedding representatives={EMBED_TESTS} (P=3..8, K=2..7)")


def unique_embedded_world(feature_count, retained_count, base_completion, base_weights):
    """Full-column-rank embedding for feature_count >= retained_count + 1."""
    smoother = global_smoothing(feature_count)
    raw_b1 = [ONE, ZERO, ZERO] + [ZERO] * (feature_count - 3)
    raw_b2 = [ZERO, ONE, ZERO] + [ZERO] * (feature_count - 3)
    raw_z = list(base_completion) + [ZERO] * (feature_count - 3)
    eta = F(1, 4 * retained_count)
    extra_count = retained_count - 2
    tau = extra_count * eta
    lam = ONE - tau
    raw_extras = []
    for extra in range(extra_count):
        col = [ZERO] * feature_count
        col[3 + extra] = ONE
        raw_extras.append(col)
    raw_y = [lam * x for x in BASE_Y] + [eta] * extra_count
    raw_y += [ZERO] * (feature_count - len(raw_y))
    raw_observed = [raw_b1, raw_b2] + raw_extras
    observed = [matvec(smoother, col) for col in raw_observed]
    y, z = matvec(smoother, raw_y), matvec(smoother, raw_z)
    weights = [lam * base_weights[0], lam * base_weights[1]]
    weights += [eta] * extra_count
    weights += [lam * base_weights[2]]
    full = columns(*(observed + [z]))
    target = conditional_retained(weights, retained_count)
    return observed, y, z, weights, full, target


UNIQUE_EMBED_TESTS = 0
for retained_count in range(2, 8):
    for feature_count in range(retained_count + 1, retained_count + 4):
        world1 = unique_embedded_world(feature_count, retained_count, E3, W1)
        world2 = unique_embedded_world(feature_count, retained_count, BASE_Z2, W2)
        obs1, y1, z1, weights1, full1, target1 = world1
        obs2, y2, z2, weights2, full2, target2 = world2
        pass_check(f"unique embedding P={feature_count}, K={retained_count}",
                   obs1 == obs2 and y1 == y2
                   and all_positive(flatten(columns(*obs1)) + y1 + z1 + z2 + weights1 + weights2)
                   and all(sum(col, ZERO) == ONE for col in obs1 + [z1, z2, y1])
                   and sum(weights1, ZERO) == ONE and sum(weights2, ZERO) == ONE
                   and matvec(full1, weights1) == y1 and matvec(full2, weights2) == y2
                   and matrix_rank(full1) == retained_count + 1
                   and matrix_rank(full2) == retained_count + 1
                   and target1[0] > target1[1] and target2[0] < target2[1])
        UNIQUE_EMBED_TESTS += 1
print(f"SCAN  | exact full-column-rank embeddings={UNIQUE_EMBED_TESTS} "
      "(K=2..7, P=K+1..K+3)")

print("\n== C2: no universal observed-data selector ==")
Q_F = [F(3, 5), F(2, 5)]
Q_R = [F(3, 10), F(7, 10)]
DELTA = tv(Q_F, Q_R)
RISKS = {
    "W1": (tv(Q_F, T1), tv(Q_R, T1)),
    "W2": (tv(Q_F, T2), tv(Q_R, T2)),
}
pass_check("fixed q_F and q_R have opposite strict risk order in the two observationally identical worlds",
           DELTA == F(3, 10)
           and RISKS == {"W1": (ZERO, DELTA), "W2": (DELTA, ZERO)})


def regret(action, world):
    risk_f, risk_r = RISKS[world]
    selected = risk_f if action == "F" else risk_r
    return selected - min(risk_f, risk_r)


MINIMAX_DETERMINISTIC_WITNESS = min(
    max(regret(action, world) for world in RISKS) for action in ("F", "R")
)
pass_check("witness lower bound for deterministic binary-selector regret is delta",
           MINIMAX_DETERMINISTIC_WITNESS == DELTA)
pass_check("witness instance of 0 <= regret <= |R_F-R_R| <= delta",
           all(ZERO <= regret(action, world)
               <= abs(RISKS[world][0] - RISKS[world][1]) <= DELTA
               for action in ("F", "R") for world in RISKS))


def randomized_regret(probability_f, world):
    risk_f, risk_r = RISKS[world]
    return (probability_f * risk_f + (ONE - probability_f) * risk_r
            - min(risk_f, risk_r))


HALF = F(1, 2)
RANDOMIZED_HALF_REGRETS = {
    world: randomized_regret(HALF, world) for world in RISKS
}
pass_check("p=1/2 randomized binary selector has witness regret delta/2 in both worlds",
           RANDOMIZED_HALF_REGRETS == {"W1": DELTA / 2, "W2": DELTA / 2})
MIDPOINT = [(a + b) / 2 for a, b in zip(Q_F, Q_R)]
MIDPOINT_ORACLE_RELATIVE_REGRETS = {
    world: tv(MIDPOINT, target) - min(RISKS[world])
    for world, target in (("W1", T1), ("W2", T2))
}
pass_check("midpoint composition has witness oracle-relative regret delta/2",
           MIDPOINT == [F(9, 20), F(11, 20)]
           and MIDPOINT_ORACLE_RELATIVE_REGRETS
           == {"W1": DELTA / 2, "W2": DELTA / 2})
RAW_RISK_CENTER = [HALF, HALF]
RAW_RISK_CLOSURE_ENDPOINTS = ([ZERO, ONE], [ONE, ZERO])
pass_check("raw-risk center has radius 1/2 on the closure of the identified set",
           max(tv(RAW_RISK_CENTER, endpoint)
               for endpoint in RAW_RISK_CLOSURE_ENDPOINTS) == HALF)
print(f"CONST | risks W1(F,R)={RISKS['W1']} W2(F,R)={RISKS['W2']}")
print(f"CONST | delta={DELTA}; deterministic={MINIMAX_DETERMINISTIC_WITNESS}; "
      f"oracle-relative randomized/composition={DELTA / 2}; raw-risk={HALF}")

print(f"\nALL CHECKS PASS | checks={PASS_COUNT}")
```
