## Additional file 4: Data S2 for "Incomplete references leave bulk deconvolution targets non-identifiable, but identification has a computable precision price": README.md

# Supplementary Data S2

This directory contains the S2b task manifest, successful beta records, envelopes, and failure manifest.

## Files and dimensions

| File | Data rows x columns | Contents |
|---|---:|---|
| `s2b_task_manifest.tsv` | 2,049 x 18 | All planned tasks |
| `s2b_success_beta_records.tsv.gz` | 13,163 x 25 | The 2,003 successful `s2b_parts/*.tsv.gz` shards merged in file-name order with one header |
| `s2b_envelopes.tsv.gz` | 1,064 x 26 | Pair-level envelopes and six-state results |
| `s2b_failure_manifest.tsv` | 46 x 17 | Solver failures, without imputation or zero encoding |

Every file retains `analysis_scope`. The counts are:

| File | `primary` | `supplementary_subject_mean` |
|---|---:|---:|
| task manifest | 1,911 tasks | 138 tasks |
| success beta records | 12,204 beta rows | 959 beta rows |
| envelopes | 952 pairs | 112 pairs |
| failure manifest | 45 failures | 1 failure |

`s2b_success_beta_records.tsv.gz` contains 896 primary pairs with successful beta records and 112 supplementary pairs. `s2b_envelopes.tsv.gz` defines the complete 952/112 pair universe. Assembly checks confirmed 2,003 shards, one 25-column schema, 13,163 rows, and the `12,204 + 959` scope split.

Read the compressed table with:

```bash
gzip -cd s2b_success_beta_records.tsv.gz | head
```

## SHA-256

These hashes cover the data payload files and exclude this README.

| File | SHA-256 |
|---|---|
| `s2b_task_manifest.tsv` | `e65e8750a5c0e4484a4c194b2d2eaf2f7fe286452d7867450d1c68b72b485397` |
| `s2b_success_beta_records.tsv.gz` | `c3e9d5ae9a98a6267ed317363a6c94c4eb7ed539987d9cfe7bc2f657916ca9be` |
| `s2b_envelopes.tsv.gz` | `2a9198a87c83ece56b28f67d6a819ed76b245f4b2012a2623460448be2c21e12` |
| `s2b_failure_manifest.tsv` | `b6656f77188c13f2062ab1a99a86c36101c02986a5c0185f9dc650003b7d1392` |
